# Connecting single-cell transcriptomes to projectomes in mouse visual cortex

**DOI:** 10.1101/2023.11.25.568393

**Authors:** Staci A. Sorensen, Nathan W. Gouwens, Yun Wang, Matt Mallory, Agata Budzillo, Rachel Dalley, Brian Lee, Olga Gliko, Hsien-chi Kuo, Xiuli Kuang, Rusty Mann, Leila Ahmadinia, Lauren Alfiler, Fahimeh Baftizadeh, Katherine Baker, Sarah Bannick, Darren Bertagnolli, Kris Bickley, Phil Bohn, Dillan Brown, Jasmine Bomben, Krissy Brouner, Chao Chen, Kai Chen, Maggie Chvilicek, Forrest Collman, Tanya Daigle, Tim Dawes, Rebecca de Frates, Nick Dee, Maxwell DePartee, Tom Egdorf, Laila El-Hifnawi, Rachel Enstrom, Luke Esposito, Colin Farrell, Rohan Gala, Andrew Glomb, Clare Gamlin, Amanda Gary, Jeff Goldy, Hong Gu, Kristen Hadley, Mike Hawrylycz, Alex Henry, Dijon Hill, Karla E. Hirokawa, Zili Huang, Katelyn Johnson, Zoe Juneau, Sara Kebede, Lisa Kim, Changkyu Lee, Phil Lesnar, Anan Li, Andrew Glomb, Yaoyao Li, Elizabeth Liang, Katie Link, Michelle Maxwell, Medea McGraw, Delissa A. McMillen, Alice Mukora, Lindsay Ng, Thomas Ochoa, Aaron Oldre, Daniel Park, Christina Alice Pom, Zoran Popovich, Lydia Potekhina, Ram Rajanbabu, Shea Ransford, Melissa Reding, Augustin Ruiz, David Sandman, La’Akea Siverts, Kimberly A. Smith, Michelle Stoecklin, Josef Sulc, Michael Tieu, Jonathan Ting, Jessica Trinh, Sara Vargas, Dave Vumbaco, Miranda Walker, Micheal Wang, Adrian Wanner, Jack Waters, Grace Williams, Julia Wilson, Wei Xiong, Ed Lein, Jim Berg, Brian Kalmbach, Shenqin Yao, Hui Gong, Qingming Luo, Lydia Ng, Uygar Sümbül, Tim Jarsky, Zizhen Yao, Bosiljka Tasic, Hongkui Zeng

## Abstract

The mammalian brain is composed of diverse neuron types that play different functional roles. Recent single-cell RNA sequencing approaches have led to a whole brain taxonomy of transcriptomically-defined cell types, yet cell type definitions that include multiple cellular properties can offer additional insights into a neuron’s role in brain circuits. While the Patch-seq method can investigate how transcriptomic properties relate to the local morphological and electrophysiological properties of cell types, linking transcriptomic identities to long-range projections is a major unresolved challenge. To address this, we collected coordinated Patch-seq and whole brain morphology data sets of excitatory neurons in mouse visual cortex. From the Patch-seq data, we defined 16 integrated morphoelectric-transcriptomic (MET)-types; in parallel, we reconstructed the complete morphologies of 300 neurons. We unified the two data sets with a multi-step classifier, to integrate cell type assignments and interrogate cross-modality relationships. We find that transcriptomic variations within and across MET-types correspond with morphological and electrophysiological phenotypes. In addition, this variation, along with the anatomical location of the cell, can be used to predict the projection targets of individual neurons. We also shed new light on infragranular cell types and circuits, including cell-type-specific, interhemispheric projections. With this approach, we establish a comprehensive, integrated taxonomy of excitatory neuron types in mouse visual cortex and create a system for integrated, high-dimensional cell type classification that can be extended to the whole brain and potentially across species.

## Introduction

The wildly varying neuronal shapes that were originally observed with the Golgi stain provided the first clues about the cellular complexity of the brain^1,2^. Our understanding of cell types has expanded greatly since then and continues to grow exponentially with recent technological advances in anatomical tracing, single cell transcriptomic characterization, and electron microscopy^3–6^. These techniques have revealed the cellular landscape of the brain at extraordinary scale and detail. However, in many cases, individual cellular properties have been studied in isolation, and we lack knowledge of the correspondences among these properties to establish robust, integrated definitions of cell types. Since cell types defined based on multiple properties^7–9^ form a stable foundation for understanding brain organization and function, studies that link genetic, functional, and circuit properties are needed.

Patch-seq is a powerful method that enables collection of electrophysiology, morphology, and transcriptomic data from the same cell^10,11^. By mapping transcriptomic signatures of Patch-seq cells to an established taxonomy of transcriptomic types (T-types)^12^, we can annotate and refine these cell type taxonomies with additional electrophysiological and morphological properties^13–18^. For example, in the hippocampus, Patch-seq was used to identify cell-type-specific expression of connectivity-related molecules^17^. In mouse and human neocortex, Patch-seq studies revealed that the morphoelectric properties of neurons largely supported transcriptomically identified cell types in both species^13,15,16^. However, these studies also observed large phenotypic variation within certain T-types, leading to ambiguity about their potential roles in the cortical circuit^16^. Patch-seq data have also been used to establish an integrated cell type taxonomy based on the morphoelectric and transcriptomic (MET) properties of neurons^13^. In mouse visual cortex, this led to the discovery of 28 inhibitory MET-types, each with type-specific axonal laminar innervation patterns that deepened our view of cell types and created priors for the circuits that they might form^13^.

In mice, transcriptomic studies of long-range projecting excitatory neurons^12^ have relied on RNA-sequencing of retrogradely labeled neurons (Retro-seq) to assign projection target subclass identity (projection subclass, for example layer 5 (L5) intratelencephalic (IT) and L5 extratelencephalic (ET)), to T-types. However, the relationship between T-types and the full set of long-range axonal projections of individual neurons has largely been missing from transcriptomic studies. This is a major gap in our knowledge since transcriptomic characterization of excitatory cortical neurons has revealed more cell types (approximately 30 types) than had previously been captured by more traditional morphoelectric characterization (9 to 19 types)^12,19–21^. Understanding the relationship between T-types and neuronal long-range projection signatures in mammals could help explain the wider transcriptomic diversity if specific projection target properties are encoded in the transcriptome of adult mice as they are in the developing fly^22^. Studies aimed at describing the complete axonal projections of single neurons are relatively few^5,6,23–27^ and describe complex targeting patterns for individual neurons that are similar to those described for the population^28,29^. Furthermore, the detailed axonal projections of individual T-types have only been established for a small number of cell types in mouse cortex^30,31^. It remains an open question for the thousands of transcriptomic types that have now been described across the whole mouse brain^3^ and for other mammalian species^32,33^.

In this study, we first generated a Patch-seq data set for mouse visual cortical excitatory neurons and defined 16 multimodal MET-types. Using these data, we characterized how transcriptomic variation frequently corresponds with electrophysiological and morphological variation as well. In parallel, we collected a mouse visual cortical data set of the complete morphology and interareal projections of individual, excitatory neurons, which we refer to as whole neuron morphology (WNM). Local morphology from Patch-seq MET-types was used as the key link between the two datasets. MET-type labels, anatomical location, and transcriptomically-correlated morphological properties were used to build models to predict specific projection targets for individual neurons. With this approach, we provide an integrated view of excitatory, transcriptomically-defined, neuron types and their phenotypic properties including neuron type-specific interareal projection patterns.

## Results

### Taxonomy of morphoelectric and transcriptomic (MET) excitatory neuron types

We used Patch-seq to investigate the correspondences between transcriptomic identity and intrinsic electrophysiological and morphological properties of excitatory neurons in adult mouse visual cortex (VIS) (Fig. 1a, top). We recorded from neurons in acute brain slices containing VIS, collected electrical responses to a standardized set of hyperpolarizing and depolarizing current stimuli, extracted the nucleus and cytosol for single-cell RNA sequencing, and filled neurons with biocytin for later morphological reconstruction. 1,544 Patch-seq recordings of excitatory neurons were included in this study (133 L2/3 neurons were previously published^16^ and were re-analyzed as part of this study.) — 1,271 neurons from primary visual cortex (VISp) and 273 from higher visual areas (HVAs). We combined cells in VISp and HVAs to describe the excitatory cell type composition of visual cortex. Finally, we generated an image-based dendritic reconstruction for 689 of those neurons with adequate biocytin fills, then calculated a set of morphological features from those reconstructions^13,21^.

**Figure 1:**
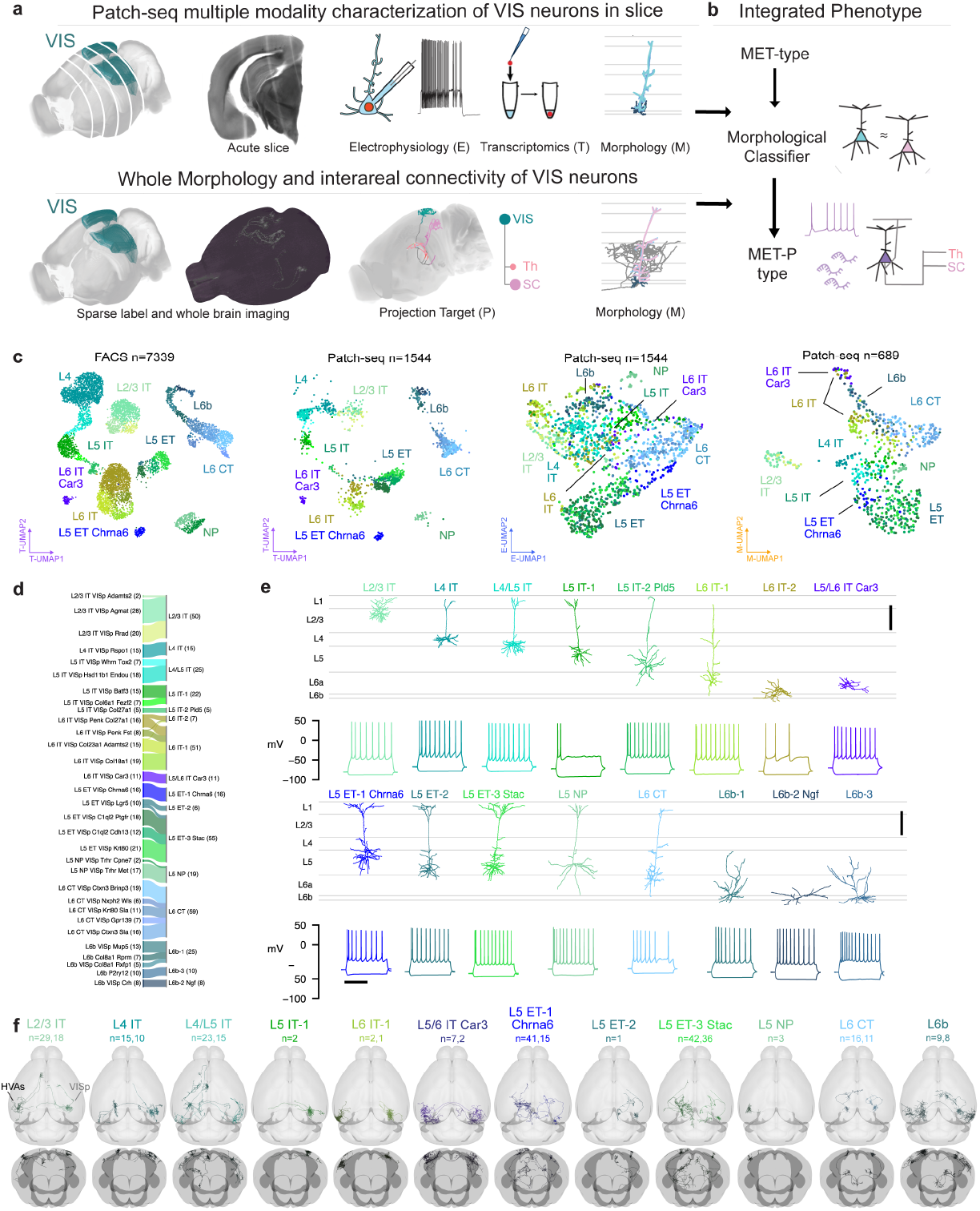
**a**, Schematic of parallel experimental strategies. **b**, Integration of the two datasets based on shared dendritic properties. **c**, UMAPs based on principal components of gene expression (left: dissociated cells, second from left: cells from Patch-seq recordings), electrophysiology features (second from right) and morphology features (right), with T-types shown in colors. Broader transcriptomic sub-classes are labeled for clarity. **d**, River plot showing the relationships between T-types (left) and assigned MET-types (right) for cells from Patch-seq recordings with all three data modalities available. **e**, Example cortical layer-aligned morphological reconstructions and electrophysiological responses for each excitatory cortical MET-type. Electrophysiology examples include responses evoked by a hyperpolarizing current step (−70 or −90 pA), and the response evoked by a rheobase + 30 pA or + 40 pA stimulus. **f**, Whole neuron morphologies (WNM) registered to the Allen CCFv3, horizontal and frontal views. Each panel shows individual WNMs located in VISp (right) and/or HVAs (left), that were classified into an integrated MET-type using local morphology features.

For experiments in which whole neuron morphology (WNM) data was collected (Fig. 1a, bottom), brains were sparsely labeled using transgenic and viral approaches, followed by whole brain fMOST imaging^34,35^ and registration to the Allen Common Coordinate Framework^36^. After that, the complete dendritic and axonal morphology of a neuron was reconstructed (Fig. 1a, bottom). The local morphologies of neurons from both data sets were aligned to an average cortical framework derived from thousands of Patch-seq experiments for integrated analyses (Fig. 1b, Extended Data Figs. 1 to 10).

Each Patch-seq neuron was assigned a transcriptomic type (T-type) using a reference taxonomy^12^ based on dissociated neurons from primary visual cortex (Fig. 1c). We visualized the transcriptomic landscape of dissociated FACS-sorted cells from the reference taxonomy^12^ and Patch-seq cells in a common Uniform Manifold Approximation and Projection (UMAP) embedding^37^ derived from the expression of 1,398 differentially expressed genes. Cells collected from Patch-seq experiments occupied similar locations within the transcriptomic UMAP as reference cells assigned to the same T-types, and relationships between transcriptomic groups were preserved between the two data sets (Fig. 1c, left two panels).

To investigate the intrinsic electrophysiological properties of these same neurons, we recorded electrical responses to a standardized current-clamp protocol^13^ and calculated a set of electrophysiological features based on those responses (Methods). We also calculated morphological features for cells that had local dendritic reconstructions. These feature sets were visualized by UMAP embedding to attain an initial visual assessment of their coherence within and diversity across T-types (Fig. 1c, right two panels). Excitatory Patch-seq neurons did not appear to be as clearly separable into distinct T-types by their intrinsic electrophysiological properties alone, but cells mapping to broader transcriptomic subclasses occupied different regions of the E-UMAP.

To establish cell types based on all three modalities, we defined MET-types using previously established techniques^13^ (Extended Data Fig. 11, Methods). We identified 16 excitatory MET-types from our data set of 384 neurons with transcriptomic and electrophysiological data and a manually-curated dendritic reconstruction (Fig. 1d, e). MET-types were named by their laminar locations, transcriptomically-derived projection types, and marker genes that distinguished them from other related MET-types (if present; see Methods and Extended Data Fig. 13). MET-types with the same layer and projection type were also distinguished with numerical suffixes. A river plot (Fig. 1d) illustrates the rlationship between T-types and MET-types. T-types frequently either merged into single MET-types (e.g., L2/3 IT VISp Rrad/Adamts2/Agmat into L2/3 IT), or had one-to-one relationships with MET-types (e.g., L4 IT VISp Rspo to L4 IT; L5 IT VISp Col27a1 to L5 IT-2 Pld5; L5 ET VISp Chrna6 to L5 ET-1 Chrna6). We also mapped Patch-seq neurons to a recently published whole brain transcriptomic taxonomy^3^ and observed a similar consolidation of those T-types into MET-types (Extended Data Fig. 12). Given the tight link between MET-type and T-type, we used T-type labels to infer MET-type labels for the remaining cells that had electrophysiological and transcriptomic data but lacked a manually-curated morphological reconstruction (Methods).

We identified eight IT MET-types (Fig. 1e, Extended Data Fig. 13): a single type per layer in L2/3 and L4, three IT MET-types in L5, two in L6, and one type that spanned infragranular layers, L5/L6 IT Car3. We also identified three ET MET-types, one L5 NP MET-type, and three L6b MET-types. Some MET-types were familiar from previous studies^19,20^, like the L2/3 IT type, which contained wide branching neurons located throughout the depth of L2/3, or the L4 IT type that had neurons with little to no apical tuft^14,38^. Other MET-types had more novel phenotypes, such as L5 IT-2 Pld5 with untufted apical dendrites, which differed from the more typical simple tufted neurons found in L5 IT-1. The L5/L6 IT Car3 type consisted of a single T-type and exhibited a stellate morphology particularly unusual for deep excitatory neurons. The L5 ET-1 Chrna6 MET-type also contained neurons from a single T-type with notably distinct gene expression patterns compared to other L5 ET neurons Extended Data Fig. 13.

When we examined the spatial distributions of MET-types across the visual cortex (Extended Data Fig. 14a-b), most MET-types had spatial distributions consistent with our sampling distribution, except for L5/L6 IT Car3 (Extended Data Fig. 14c). This MET-type was more frequently found in lateral VISp and lateral HVAs like VISl, similar to previous studies^27,39^. We also examined the cortical layer distributions of Patch-seq neurons (Extended Data Fig. 14d). MET-types were either found predominantly in a specific layer or in two neighboring layers, in accordance with previous findings^12,15,19,21,39,40^.

In order to link MET-types to neurons from the WNM data set, we developed a multi-step classification protocol. This method leveraged both dendritic morphology and projection subclass (derived transcriptomically for Patch-seq and anatomically for WNM) to generate robust MET-type predictions (Extended Data Fig. 15). These MET-type assignments enabled predictions about the axonal properties of excitatory MET-types, and, conversely, about the electrophysiological and transcriptomic properties that accompany neurons with these axonal phenotypes. Individual neurons within each MET-type had complex targeting patterns (Fig. 1f), similar to what has been described previously for other brain regions^5,24,27^. However, on average, each predicted MET-type had interareal projection patterns distinct from each other. This morphology-based cross-data set mapping provided a more complete description of neuronal cell types.

### Transcriptomic variation and cross-modal correspondence

Though several T-types were typically merged into a single MET-type, we observed that neurons within a given MET-type sometimes exhibited heterogeneous transcriptomic, electrophysiological, and morphological properties. To examine whether these variations were coordinated across properties, we performed principal component analysis (PCA) on the highly-variable genes expressed in reference FACS-derived neurons within each transcriptomically defined subclass (Methods). These transcriptomic PCs (Tx PCs) were generated in the same way as those used to define T-types in the original taxonomy^12^, and the first few Tx PCs within each subclass could usually distinguish its member T-types (Extended Data Fig. 16a). We also performed a dimensionality reduction with weighted gene co-expression network analysis (WGCNA)^41^ to identify gene modules associated with those PCs that could be linked to functionally relevant categories (Extended Data Figs. 17 and 18). To examine the correspondence between transcriptomic variation and morphoelectric properties within single neurons collected with Patch-seq, we measured the correlations between the Tx PCs and electrophysiological and morphological features. Statistically significant correlations (after adjustment for multiple comparisons) were examined to characterize the coordinated heterogeneity with MET-types or groups of MET-types.

#### IT MET-types

In the L2/3 IT MET-type, we found a number of significant correlations, especially with the second transcriptomic principal component (L2/3 IT Tx PC-2, Fig. 2a, Extended Data Fig. 19a). For example, electrophysiology features relating to the shape of the action potential were correlated with Tx PC-2 (Fig. 2b); neurons with higher Tx PC-2 values had narrower action potentials with a steeper downstroke (Fig. 2c). Morphological features, such as the depth of the neuron from the pial surface, also were correlated with this transcriptomic PC (Fig. 2d, f).

**Figure 2:**
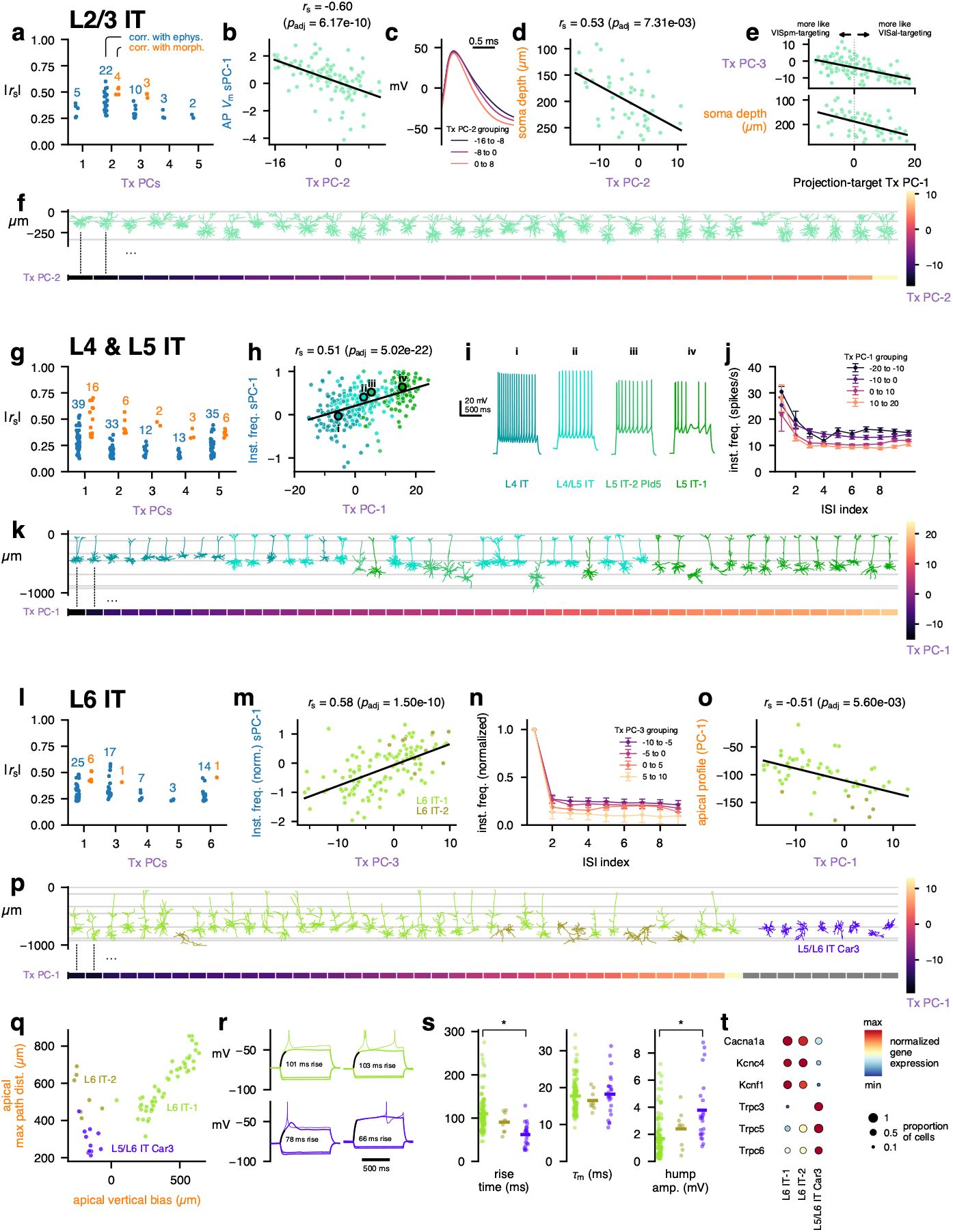
MET characterization of IT subclasses. **a**, Spearman correlations between transcriptomic principal components (Tx PCs) and electrophysiology/morphology features for the L2/3 IT subclass. Only statistically significant correlations are shown (B-H adjusted p-values < 0.05); the number of significant correlations per PC and modality are shown above the points. **b**, Relationship between L2/3 IT Tx PC-2 and the first sparse principal component (sPC-1) of the action potential (AP) waveform. **c**, Average AP waveforms for cells grouped by transcriptomic PC-2 values. **d**, Relationship between L2/3 IT Tx PC-2 and soma depth. **e**, Relationships between a PC derived from VISpm- vs VISal-targeting differentially expressed genes identified by Kim *et al.* [42] and L2/3 IT Tx PC-3 (top) and soma depth (bottom). **f**, Example L2/3 IT morphologies (top) ordered by their Tx PC-2 values (bottom). **g**, Correlations between L4 & L5 IT (L4 IT, L4/L5 IT, L5 IT-1, and L5 IT-2 Pld5) Tx PCs and electrophysiology/morphology features. **h**, Relationship between L4 & L5 IT Tx PC-1 and the sPC-1 of instantaneous firing frequency. **i**, Example responses to depolarizing current steps for cells corresponding to points i to iv in (**i**). **j**, Average instantaneous firing frequencies by interspike interval (ISI) index for cells grouped by L4 & L5 IT transcriptomic PC-1. **k**, Example L4 & L5 IT morphologies (top) ordered by their Tx PC-1 values (bottom). **l**, Correlations between L6 IT (L6 IT-1 and -2) Tx PCs and electrophysiology/morphology features. Note that L6 IT Tx PC-2 is not shown as it did not exhibit any significant correlations with electrophysiology/morphology features. **m**, Relationship between L6 IT Tx PC-3 and normalized instantaneous firing frequency sPC-1. **n**, Average instantaneous firing frequencies, normalized to the first ISI, by ISI index for cells grouped by L6 IT Tx PC-3. **o**, Relationship between L6 IT Tx PC-1 and the apical depth profile PC-1. **p**, Example L6 IT morphologies (top) ordered by their Tx PC-1 values (bottom). Example L5/L6 IT Car3 morphologies are also shown for comparison; they were not included in the L6 IT PCA as they exhibit highly distinct transcriptomic profiles and so do not have Tx PC-1 values (gray). **q**, Relationship between the apical vertical bias and maximum path distance within the apical dendrite. **r**, Example responses to subthreshold (thick line) and suprathreshold (thin line) depolarizing current steps. The interval in which the voltage rose between 10% and 90% of its steady-state value is indicated (black) and the rise time shown. Colors indicate MET-types as in (**q**). **s**, Comparison of rise times (left), membrane time constants (middle), and depolarizing “hump” amplitudes (right) between L6 IT-1, -2, and L5/L6 IT Car3 cells. **t**, Differentially expressed ion channels between L6 IT-1 and L5/L6 IT Car3. L6 IT-2 also shown for comparison.

We were interested in whether the expression of certain functionally relevant categories of genes, such as voltage-gated ion channels, varied along these same transcriptomic dimensions; however, these genes were not reliably found in the set of highly variable genes used to calculate the PCs and therefore often did not have a PC weight to examine. Therefore, we measured the correlations between the Tx PCs and the expression of voltage-gated ion channels (Extended Data Fig. 20), ligand-gated ion channels (Extended Data Fig. 21), cell-adhesion molecules (CAMs, Extended Data Fig. 22), and synaptic exocytosis-related molecules (Extended Data Fig. 23). Interestingly, for L2/3 IT neurons, few ion channels were strongly correlated with Tx PC-2, but several CAMs (e.g., *Ptprk*, *Fam19a1*) and synaptic molecules (e.g., *Rims3*, *Syt4*) were correlated with the transcriptomic gradient.

A previous study^42^ identified differentially expressed genes within L2/3 IT neurons of mouse primary visual cortex that were related to their HVA projection targets; the authors of that study identified genes that were associated with axonal projections to either VISpm or VISal. While we do not know the projection targets of the neurons in our Patch-seq data set, we tested whether the VISpm-projecting vs. VISal-projecting transcriptomic signature was related to other properties of the cells. We calculated the first PC of the projection target-specific differentially expressed genes of Kim *et al.* [42] and compared them with the Patch-seq transcriptomic PCs and other features. The projection target PC-1 was most strongly correlated with L2/3 IT Tx PC-3 (Fig. 2e). It, too, was correlated with soma depth, with deeper cells exhibiting a more VISal-targeting transcriptomic signature, consistent with the relationship between cortical depth and projection target reported by Kim *et al.* [42].

Interestingly, the L2/3 IT Tx PC-1, which represented 13.5% of the variance in the population, was not correlated with many electrophysiological or morphological features (Fig. 2a). This PC was, however, highly correlated with activity-dependent gene expression (Extended Data Fig. 24a-c), consistent with the results of Tasic *et al.* [12] who reported that activity-dependent genes were differentially expressed across the L2/3 IT T-types. Several activity-dependent genes (e.g., *Fos*, *Fosb*, and *Npas4*) exhibited increased expression in Patch-seq cells compared to the reference FACS data set^12^ (Extended Data Fig. 24e-f). We also noted relatively few Patch-seq cells mapping to the L2/3 IT VISp Adamts2 T-type and relatively more to the L2/3 IT VISp Rrad T-type^12^ (reference data set: 68% Agmat, 21% Adamts2, 11% Rrad; Patch-seq data set: 53% Agmat, 41% Rrad, 6% Adamts2). When we removed the activity-dependent signal and reclustered the reference cells (Methods), those two T-types were merged together (Extended Data Fig. 24d), suggesting that these T-types may represent high and low activity states, respectively, and that the process of preparing and recording from Patch-seq cells may increase the expression of certain genes that results in mapping to the latter type.

Next, we examined coordinated variation across modalities for the MET-types belonging to the L4 and L5 IT subclass (L4 IT, L4/L5 IT, L5 IT-1, and L5 IT-2 Pld5). Multiple significant correlations with electrophysiological and morphological features were identified (Fig. 2g, Extended Data Fig. 19b), including a relationship between instantaneous firing frequency during depolarizing current steps and Tx PC-1 (Fig. 2h). Cells of the L4 IT type typically had lower values of L4 & L5 IT Tx PC-1, while L5 IT-1 had higher values, with L4/L5 IT and L5 IT-2 Pld5 in between. This corresponded with a higher sustained firing frequency among the cells with lower Tx PC-1 values (Fig. 2i-j). Tx PC-1 was also correlated with several morphological features (Extended Data Fig. 19b), which could be observed by sorting the cells by Tx PC-1 (Fig. 2k): L4 IT cells at shallower depths with smaller apical tufts transitioned to L4/L5 IT cells and L5 IT-1 cells deeper in cortex. Interestingly, L5 IT-2 Pld5 cells (which had intermediate Tx PC-1 values) were found deeper in cortex but had less elaborate apical tufts than L4/L5 IT and L5 IT-2 cells. The L4 & L5 IT Tx PC-1 was also correlated with several voltage-gated ion channel genes (e.g., positively with *Cacna1d* and *Kcna6* and negatively with *Kcnh5*, Extended Data Fig. 20), ligand-gated ion channel genes (e.g., positively with *Grik3* and negatively with *Grik1*, Extended Data Fig. 21), and CAMs (e.g., positively with *Ilrapl2*, negatively with *Nrxn3*, Extended Data Fig. 22).

To compare transcriptomic variations with morphoelectric features for L6 IT neurons, we performed PCA on L6 IT-1 and L6 IT-2 cells, but not L5/L6 IT Car3 cells, since the latter cells are quite transcriptomically distinct and have been classified into their own subclass in recent cortical taxonomies^3,39^. L6 IT-1 and L6 IT-2 neurons had somewhat weaker correlations with electrophysiological and morphological features compared with L4 & L5 IT cells (Fig. 2l, Extended Data Fig. 19c) but did exhibit some relationships, such as with the normalized instantaneous firing frequency (Fig. 2m). Cells with higher values of L6 IT Tx PC-3 had higher normalized steady-state firing frequencies (Fig. 2n). Tx PC-3 was negatively correlated with *Kcnc2* expression and positively correlated with *Kcnq5* (Extended Data Fig. 20). L6 IT Tx PC-1 was the most correlated with morphological features, including the apical depth profile PC-1 feature (Fig. 2o); however, this PC did not clearly separate L6 IT-2 cells (with an inverted apical dendrite) from L6 IT-1 cells (Fig. 2p). This PC exhibited strong positive correlations with several CAMs (*Nptx2*, *Ptpru*, Extended Data Fig. 22) and synaptic exocytosis-related molecules (*Cplx2*, Extended Data Fig. 23).

None of the transcriptomic PCs calculated from L5/L6 IT Car3 cells alone were significantly correlated with electrophysiological or morphological features, though this could be in part due to the smaller number of cells collected for that type (*n* = 11 with triple-modality data, *n* = 13 with only E/T data). The L5/L6 IT Car3 neurons could be distinguished morphologically from other L6 IT cells by their smaller inverted apical dendrites (Fig. 2p-q). They also exhibited distinct electrophysiological features, such as a faster rise time during depolarizing current steps despite having similar membrane time constants (Fig. 2r-s). Several of the L5/L6 IT Car3 cells also exhibited large depolarizing “humps” at potentials just below the AP firing threshold (Fig. 2r-s); the timing of this hump frequently corresponded to the timing of the first AP at rheobase. The L5/L6 IT Car3 cells expressed several different ion channels compared with L6 IT-1 cells (Fig. 2t) — the L5/L6 IT Car3 cells did not express genes for P/Q-type Ca channels (*Cacna1a*), Kv3.4 channels (*Kcnc4*), or Kv5.1 channels (*Kcnf1*) but did express several Trpc channels (*Trpc3*, *Trpc5*, and *Trpc6*) that were absent in L6 IT-1 and L6 IT-2 cells.

#### ET, NP, CT, and L6b MET-types

L5 ET cells had several transcriptomic PCs that were correlated with electrophysiological and morphological features; L5 ET Tx PC-1 had stronger correlations with morphology, while L5 ET Tx PC-2 was more correlated with electrophysiology (Fig. 3a, Extended Data Fig. 25a). The L5 ET-1 Chrna6 cells, along with L5 ET-2 cells, typically did not fire AP bursts during depolarizing current steps, unlike many (but not all) of the L5 ET-3 Stac cells (Fig. 3b). This could be seen by comparing the maximum instantaneous firing frequency during a current step to the total number of APs fired during that step (Fig. 3c, left) — cells that fired bursts had very high instantaneous frequencies but only fired a few APs. The ratio between those two values was significantly correlated with Tx PC-2 (Fig. 3c). In addition, several voltage-gated ion channels were differentially expressed between L5 ET-1 Chrna6 and L5 ET-3 Stac (Fig. 3d), including a T-type Ca channel (*Cacna1h*) that was present in L5 ET-3 Stac but not L5 ET-1 Chrna6 cells. In accordance with those findings, several of these channels (e.g., *Cacna1h*, *Trpc4*, *Trpc5*) were positively correlated with Tx PC-2, along with others (such as *Hcn1*) that did not meet the criteria for differential expression but did vary along the similar gradient (Extended Data Fig. 20).

**Figure 3:**
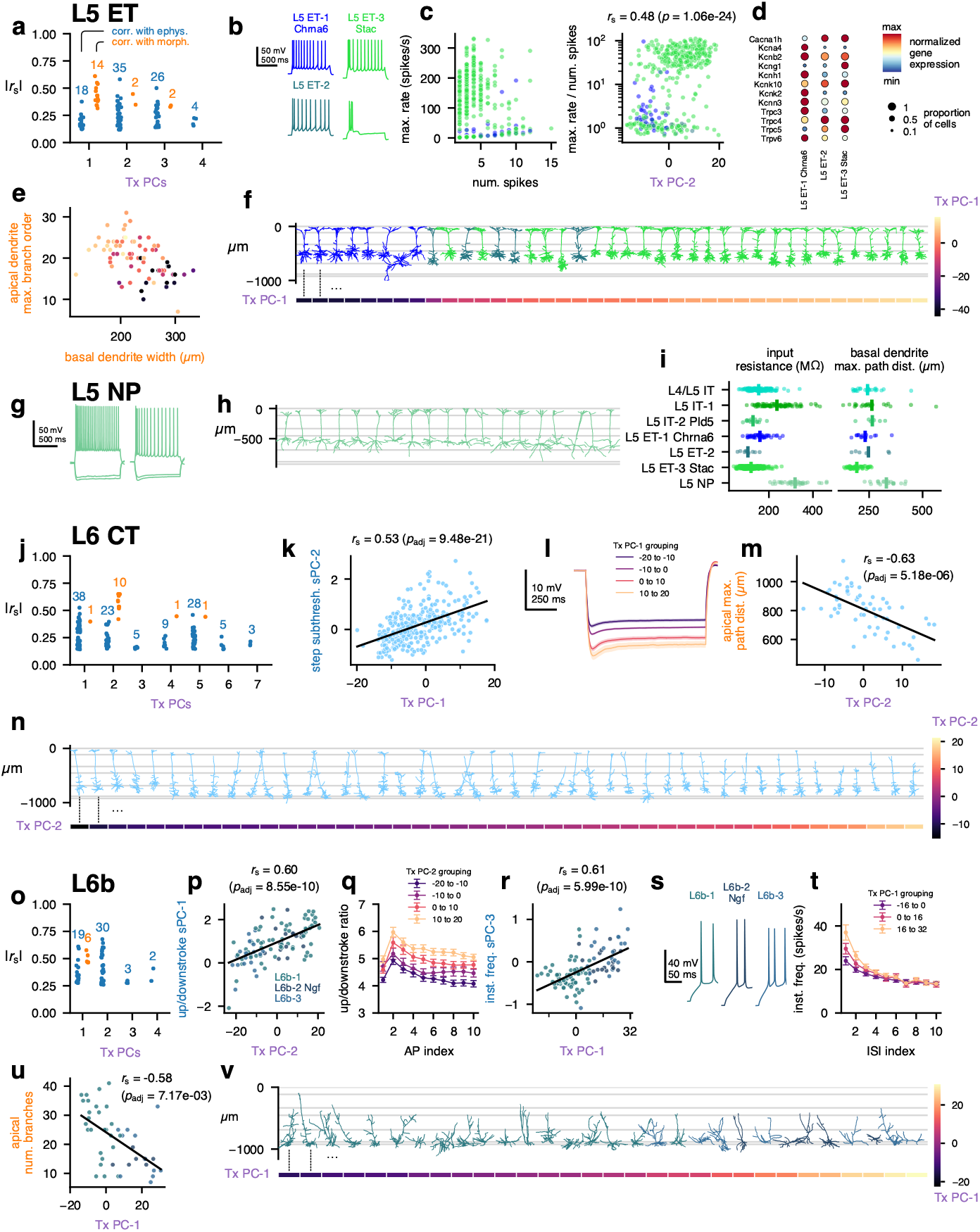
MET characterization of ET, NP, CT, and L6b subclasses. **a**, Spearman correlations between transcriptomic principal components (Tx PCs) and electrophysiology/morphology features for the L5 ET (L5 ET-1 Chrna6, L5 ET-2, and L5 ET-3 Stac) subclass. Only statistically significant correlations are shown (B-H adjusted p-values < 0.05); the number of significant correlations per PC and modality are shown above the points. **b**, Example responses to depolarizing current steps showing different initial activity, including bursting. **c**, Relationship between the maximum instantaneous firing frequency during a current step and the number of total APs in that step (left), and the relationship between that ratio and the L5 ET Tx PC-2 (right). **d**, Differentially expressed ion channels between L5 ET-1 Chrna6 and L5 ET-3 Stac. L5 ET-2 also shown for comparison. **e**, Relationship between the width of the basal dendrites and the maximum branch order of the apical dendrite. Colors indicate the L5 ET Tx PC-1 value; see (**f**) for color scale. **f**, Example L5 ET morphologies (top) ordered by their Tx PC-1 values (bottom). **g**, Example responses to hyperpolarizing and depolarizing current steps from L5 NP cells. **h**, Example L5 NP morphologies. **i**, Comparison of input resistance (left) and basal dendrite maximum path distances (right) across L5 MET-types. L5 NP cells had significant differences in input resistance to all other classes except L5 IT-1 (post hoc Dunn’s test *p* = 3.4 × 10^−30^ to 1.3 × 10^−7^ following K-W test *p* = 3.8 × 10^−61^), and significant differences in path distances to all others but L5 IT-2 Pld5 (post hoc Dunn’s test *p* = 3.1 × 10^−12^ to 0.018 following K-W test *p* = 2.3 × 10^−11^). **j**, Correlations between L6 CT Tx PCs and electrophysiology/morphology features. **k**, Relationship between L6 CT Tx PC-1 and the first sparse principal component (sPC-1) of the step subthreshold responses. **l**, Average responses to −90 pA current steps grouped by L6 CT Tx PC-1. **m**, Relationship between L6 CT Tx PC-2 and the maximum path distance within the apical dendrite. **n**, Example L6 CT morphologies (top) ordered by their Tx PC-2 values (bottom). **o**, Correlations between L6b (L6b-1, L6b-2 Ngf and L6b-3) Tx PCs and electrophysiology/morphology features. **p**, Relationship between L6b Tx PC-2 and upstroke/downstroke ratio sPC-1. **q**, Average upstroke/downstroke ratio by AP number grouped by L6b Tx PC-2. **r**, Relationship between L6b Tx PC-1 and instantaneous firing frequency sPC-3. **s**, Example initial responses to depolarizing current steps from each L6b MET-type. **t**, Average instantaneous firing frequencies versus ISI index grouped by L6b Tx PC-1. **u**, Relationship between L6b Tx PC-1 and number of branches in the apical dendrite. **v**, Example L6b morphologies (top) ordered by their Tx PC-1 values (bottom).

The morphologies of L5 ET cells varied systematically in the relationship between their apical and basal dendritic fields; cells with more branching in their apical tufts had narrower basal fields, and vice versa (Fig. 3e). This variation corresponded with Tx PC-1 (Fig. 3e, f), as cells with lower values of Tx PC-1 (which included most L5 ET-1 Chrna6 and L5 ET-2 cells) had simpler apical tufts and wider basal dendrites. Tx PC-1 exhibited strong correlations with multiple CAMs (e.g., positively with *Cdh13*, *Cntnap5a*, and *Pcdh7*; negatively with *Nrg1*, *Fat3*, and *Car4*; Extended Data Fig. 22) and synaptic molecules (e.g., *Snca* and *Stxbp6*, Extended Data Fig. 23).

The L5 NP cells did not exhibit significant correlations between transcriptomic PCs and electrophysiological or morphological features, although our ability to detect those correlations could be limited by the smaller number of L5 NP cells recorded (*n* = 19 triple modality cells, *n* = 16 with only E/T data). Still, L5 NP cells exhibited high input resistances, relatively strong spike-frequency adaptation (Fig. 3g, i), and long basal dendrites (Fig. 3h), which distinguished them from other L5 excitatory MET-types (Fig. 3i).

The transcriptomic PCs of neurons in the L6 CT MET-type were moderately correlated with electrophysiological features but more strongly correlated with morphology (Fig. 3j, Extended Data Fig. 25b). L6 CT Tx PC-1 was correlated with the responses to hyperpolarizing current steps (Fig. 3k); higher values corresponded to greater hyperpolarization from the same current stimulus (Fig. 3l). Many of the morphological features that were correlated with L6 CT Tx PC-2 were related to the size of the apical dendrite (Fig. 3m, Extended Data Fig. 25). When cells were sorted by their Tx PC-2 values, the tallest cells were found at the lower end and the smallest at the higher end (Fig. 3n). Tx PC-2 was positively correlated with CAMs such as *Cdh13* and *Nptx2* (Extended Data Fig. 22).

Cells in the L6b MET-types (L6b-1, L6b-2 Ngf, and L6b-3) had relatively strong correlations between transcriptomic PCs and electrophysiological features (Fig. 3o, Extended Data Fig. 25), especially L6b Tx PC-2, which corresponded to differences in the upstroke/downstroke ratio across APs during current steps (Fig. 3p-q). Tx PC-2 was positively correlated with *Cacna1b* and *Kcnk2* expression (Extended Data Fig. 20). L6b Tx PC-1 was correlated with additional electrophysiological features as well as morphological ones. Cells with lower values of Tx PC-1 (which were often of the L6b-1 MET-type) had wider initial interspike intervals (ISIs), while high values (typically L6b-2 Ngf and L6b-3) had faster doublets at the start of the AP train (Fig. 3r-t). Higher values of Tx PC-1 were also correlated with less branched apical dendrites, which were often oriented in directions other than toward the pia (Fig. 3u-v). Tx PC-1 was positively correlated with the ion channel genes *Kcnn3* and *Kcnt2* (Extended Data Fig. 20) along with multiple CAMs such as *Cntn5*, *Epha7*, and *Fam19a5* (Extended Data Fig. 22).

### Whole neuron morphology of predicted MET-types

A major question about transcriptomically-defined cell types is to what extent they reflect the detailed, axonal projections of a neuron. To understand the relationship between MET-types and the pattern of local and interareal axonal projections, we generated a dataset of 306 VIS cortical neurons reconstructed from whole brain fMOST images (Fig. 1a,f, Fig. 4, Fig. 5). Neurons were labeled using transgenic mouse lines and viral labeling strategies to achieve broad coverage across layers and projection neuron subclasses (Supplementary Data Table 1). Successful strategies sparsely and robustly labeled neurons either across the brain or only within VIS^27^. Where available, we also used mouse lines that selectively label T-types (i.e., Chrna6-IRES2-FlpO-WPRE-neo) and/or subclasses (i.e., Nxph4-T2A-CreERT2; Nxph4 is a marker gene for L6b subclass neurons). We prioritized neuron reconstructions in L2/3 through L6b of VISp (*n* = 161), but a smaller set of neurons was also reconstructed in HVAs (VISl *n* = 22, VISam *n* = 22, VISpm *n* = 21, VISpor *n* = 18, VISal *n* = 18, VISrl *n* = 16, VISa *n* = 14, VISpl *n* = 9, VISli *n* = 5). A subset of these neurons, 36 VISp, 3 VISa, 8 VISal, 4 VISam, 6 VISl, 2 VISli, 2 VISpm, 2 VISpor, and 10 VISrl neurons, was previously published^27^ and was re-registered and re-analyzed for this study. We examine the full morphologies of VISp neurons (Fig. 4) and then compare to HVA neurons in (Fig. 5).

**Figure 4:**
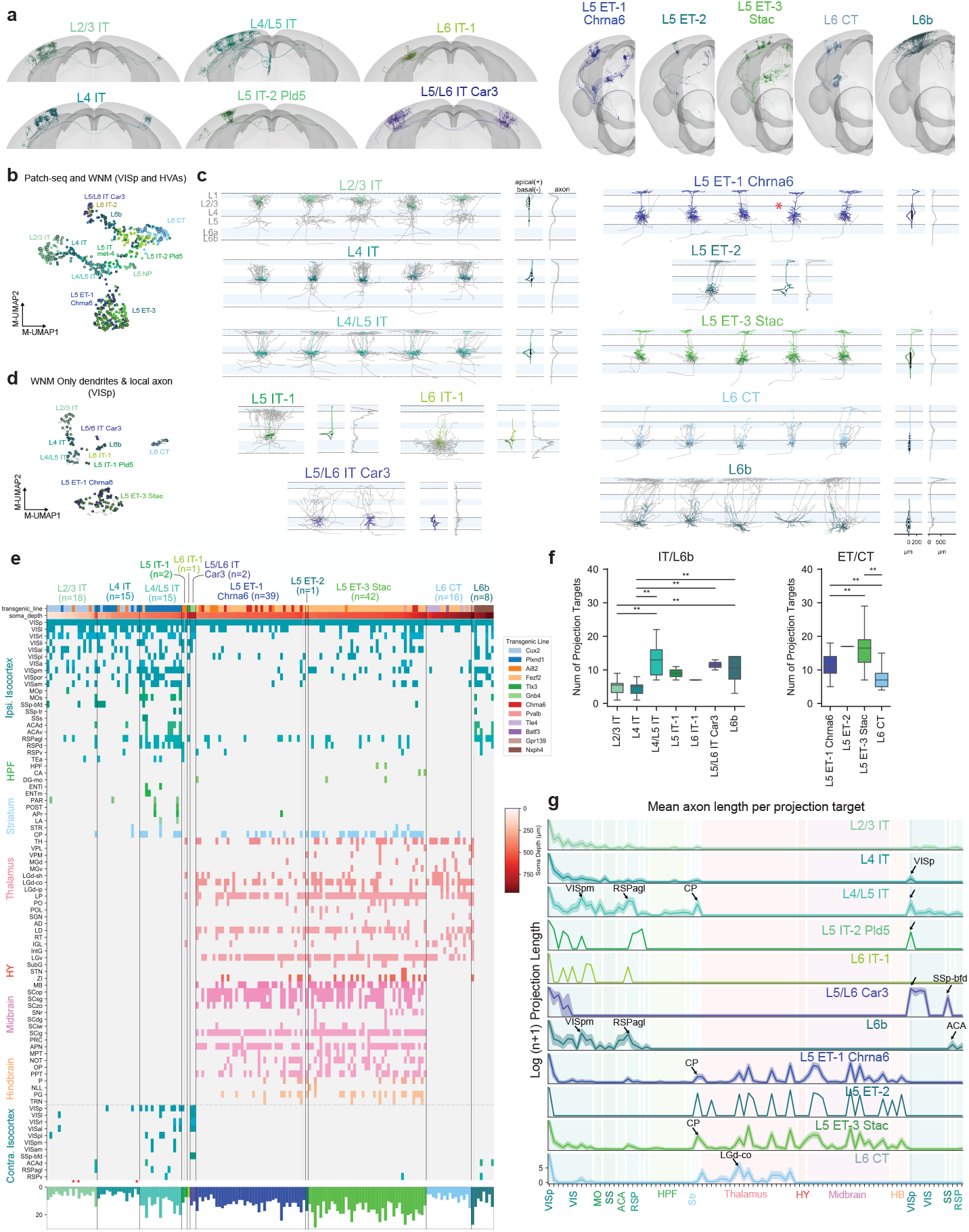
Local morphology and long-range projections of predicted VISp MET-types. **a**, Example WNMs of predicted MET-types registered to CCFv3. **b**, UMAP based on principal components of dendritic morphology and soma location, colored by MET-type. WNMs are also circled in black. **c**, Local morphologies of predicted MET-types shown in (**a**). Example morphologies were selected by calculating a pairwise similarity score for dendrites, local axon, and laminar location. Dendrites of example neurons appear in MET-type colors. Local axon appears in gray. Starred neurons indicate the reconstructions that were generated based on T-type-specific Chrna6-IRES2-FlpO mice. **d**, UMAP based on principal components of dendritic and local axon morphology and soma location, colored by MET-type (WNMs only). **e**, Binary projection target matrix ordered first by MET-type and then by normalized cortical depth. Transgenic mouse line and soma depth are also indicated with a color bar at the top. Histograms at the bottom show the total number of targets per neuron. A “target” was defined as a CCFv3 brain region containing a branch node or tip. Targets shown were contacted either by at least three neurons or at least two neurons from the same MET-type. A full projection target matrix appears in Supplementary Data Table 1. Stars indicate “local” neurons that do not project out of their soma region. **f**, Box-and-whisker plots showing the average number of targets per MET-type. To determine significant differences in the number of MET-type targets, Kruskal-Wallis tests were performed followed by post-hoc Dunn tests. **g**, Projection target histogram summaries ordered by MET-type. Mean (lines) +/− SEM (shaded regions).

**Figure 5:**
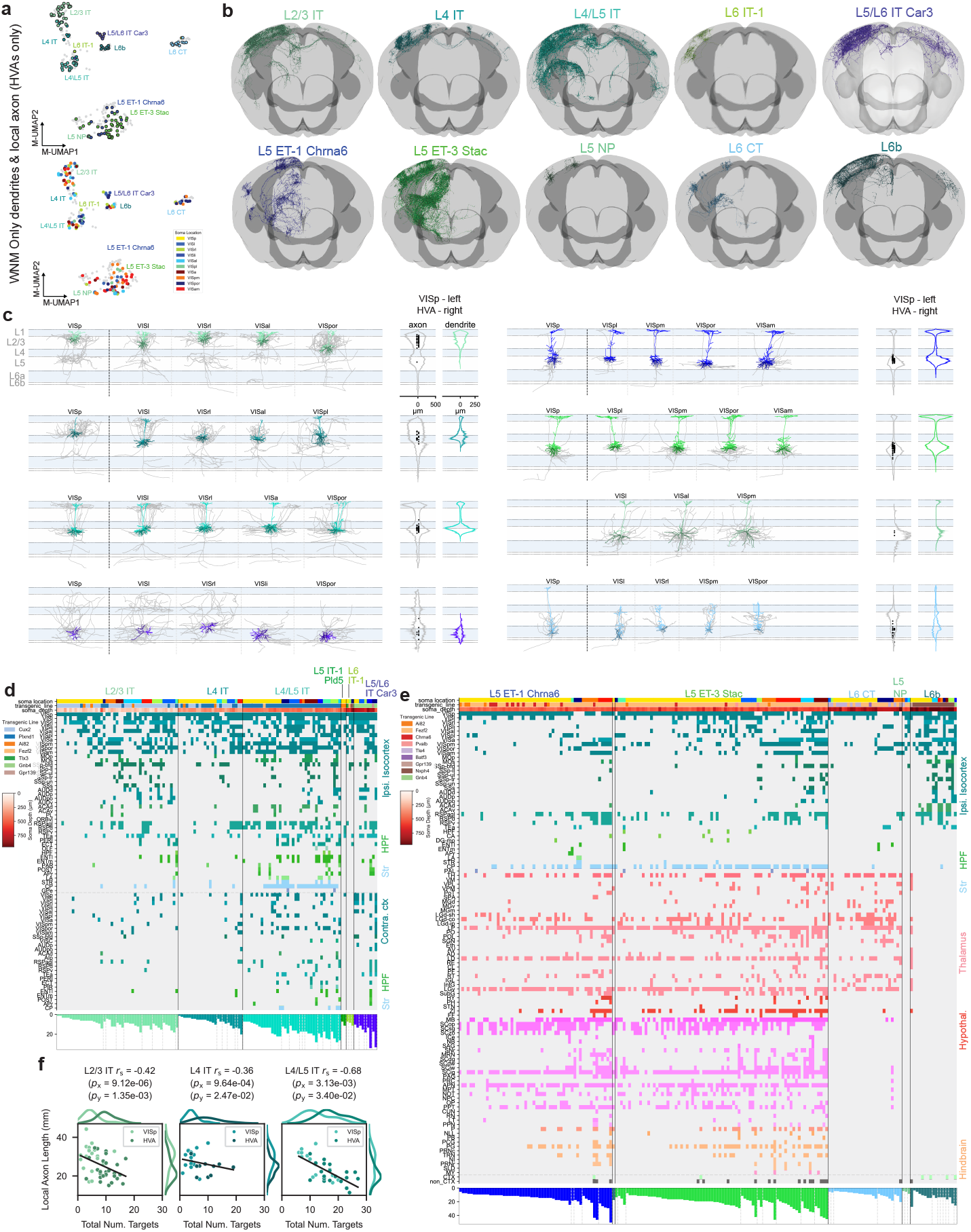
Local morphology and long-range projections of predicted MET-types in higher visual areas (HVAs), compared to VISp. **a**, Top: UMAP based on principal components of dendritic and local axon morphology. HVA neurons are colored by MET-type; VISp neurons appear in gray. Middle: VIS cortical flatmap colored by structure. Bottom: UMAP based on principal components of dendritic and local axon morphology. HVA neurons are colored by soma location; VISp neurons appear in gray. Only WNMs shown in top and bottom. **b**, WNM of example neurons registered to CCFv3 by predicted IT MET-types and non-IT MET-types. **c**, Example local morphologies of predicted IT MET-types and non-IT MET-types in different VIS brain regions. Dendrites appear in MET-type colors. Local axon, which was not collected for the Patch-seq dataset, appears in gray. Only abundantly represented MET-types are shown in the figure. **d**, Binary projection target matrices for IT MET-types. Matrix is ordered first by MET-type, then by brain region sorted by the smallest to largest number of projection targets per region. Transgenic mouse line and soma depth are also indicated with a color bar at the top of the matrix. Histograms at the bottom show the total number of targets per neuron. **e**, Binary projection target matrices for non-IT MET-types. Matrix is ordered first by MET-type, then by brain region sorted by the smallest to largest number of projection targets per region. Transgenic mouse line and soma depth are also indicated with a color bar at the top of the matrix. Histograms at the bottom show the total number of targets per neuron. **f**, Scatter plots illustrating the relationship between local axon total length and the number of projection targets for IT MET-types in VISp and HVA neurons.

#### Linking MET-types to axonal projections through dendritic morphology

To link MET-types defined with Patch-seq to WNM data, we used the classifier described above (see Fig. 1) that relied on dendritic features and projection subclass assignments from both data sets (Extended Data Fig. 15). Examples of WNMs by predicted MET-types are shown in Figure 4a. Before running the classifier, multiple processing steps were performed on the WNM data to ensure feature alignment with Patch-seq (Extended Data Fig. 15). A UMAP derived from dendritic components suggested good alignment between the two datasets (Fig. 4b); as expected, neurons that mapped to the same MET-type were grouped together, regardless of data set.

For VISp neurons, we identified six of the eight Patch-seq-defined IT MET-types in the VISp WNM dataset. The missing MET-types were likely the result of fewer whole brain morphologies from the L5 and L6 IT subclasses. We identified all three L5 ET MET-types and the L6 CT MET-type in the VISp WNM dataset. Since L6b MET-types, as mentioned, could not reliably be distinguished from their dendritic morphologies alone (Extended Data Fig. 15), we mapped WNMs to the L6b subclass rather than individual MET-types. When we visually compared MET-types across data sets, we found consistent dendritic phenotypes (Fig. 4c, Extended Data Fig. 26, Extended Data Fig. 27).

Importantly, our VISp mapping results also agreed with the type and subclass-specific transgenic mice from which these neurons were sampled (Extended Data Fig. 28). Specifically, all predicted L5/L6 IT Car3 neurons in VISp were labeled by the Gnb4 mouse line, which has previously been described to label neurons in the Car3 subclass^27^. Similarly, 10 out of 13 neurons labeled by the T-type-specific Chrna6-IRES2-FlpO mouse mapped to the L5 ET-1 Chrna6 MET-type (for comparison, in the Patch-seq data set, 24 out of 26 neurons from the Chrna6-IRES2-FlpO mouse mapped to the L5 ET VISp Chrna6 T-type). L6b neurons were all labeled by the Nxph4-CreER line, which selectively labels these neurons throughout the brain^43^.

#### Local axon of predicted MET-types

When we examined the local axon (defined as the axon within a 500 µm radius cylinder centered on the soma) of these predicted MET-types, we also saw clear differences across types (Fig. 4c, histograms to the right of each morphology panel). L2/3 IT neurons predominantly innervated superficial L1, L2/3 and L5. L4 IT neurons had the very distinct, dense, and highly columnar local axon, predominantly found in L2/3 and L4, typically observed for L4 sensory cortical neurons^38^. Predicted L4/L5 IT neurons had radially projecting axon that elaborated in L1 and superficial L5. In contrast, the one L5 IT-1-typed neuron densely innervated L2/3 and deep L5. L6 IT-1 had axonal projections largely restricted to L6, while L5/L6 IT Car3 neurons had sparse axon in all layers.

L5 ET-1 Chrna6 and L5 ET-3 Stac neurons had relatively minimal local axon distributed in L5 and, to a lesser extent, in L1. In contrast, the single L5 ET-2-typed neuron had abundant L1 axon. L6 CT neurons had very distinct local axon largely restricted to L4 to L6. The deeper neurons also had one or two axon collaterals that reached L1. L6b neurons most strongly innervated L1, which agreed with previous descriptions of these neurons^44,45^. The UMAP in Figure 4d, generated from WNM dendritic and axonal features, suggested how local axon could distinguish MET-types, in particular the neurons in L6 (L6b, L6 CT, L5/L6 IT Car3) that were grouped together in the dendritic-only UMAP (Fig. 4b). To confirm, we tested how well local axon alone could predict MET-types in the WNM data and we achieved 81% accuracy (Extended Data Fig. 29). These findings supported our MET-type assignments by demonstrating that local axon was also consistently different across MET-types.

#### Long-range projections of predicted MET-types

When we examined the long-range projections of MET-types, we found distinct rules for their projection target patterns (Fig. 4e-g) and axonal properties (e.g., total axon length) (Fig. 4g, Extended Data Fig. 30), though there was still variation across individual neurons in the specific set of targets. We defined a “target” as any CCFv3 structure containing at least one axonal branch point or ending^46^. This target definition was more inclusive than that used in previous studies requiring a minimum 1 mm axon length^24,26,27^, which resulted in a comparatively larger number of per-neuron targets identified here. For example, L2/3 IT neurons had an average of six cortical targets, mostly restricted to the HVAs (Fig. 4f). They rarely projected contralaterally, consistent with other observations^24,27^. When they did, they contacted contralateral HVAs only (Fig. 4e,g). Interestingly, we observed two “local” L2/3 neurons with axon restricted to VISp (starred in Fig. 4e). L4 IT neurons had few targets and rarely projected contralaterally. When they did, they exclusively contacted VISp (Fig. 4e,g, arrow). L4 neurons have often been considered to be “local” neurons; however, in our data set most of them did project to other brain regions (Fig. 4e,g), consistent with previous observations in the mouse^12,28^.

L4/L5 IT neurons targeted a much larger number of ipsilateral and contralateral brain regions (up to 23). Over 60% of these neurons projected contralaterally, compared to less than 20% of either L2/3 and L4 IT MET-types (Fig. 4e). Like contralateral-projecting L4 IT neurons, contralateral-projecting L4/L5 IT neurons most consistently targeted VISp (Fig. 4e, g, arrow; Extended Data Fig. 31). L4/L5 IT neurons targeted multiple HVAs on both sides of the brain, with ipsilateral VISpm targeted most commonly for this set of neurons (Extended Data Fig. 31). Of note, this was also the most common target for L2/3 neurons in a previous study^24^; however, for our L2/3 dataset, VISrl was the most common target (Extended Data Fig. 31). Variables that influence the targeting of specific cortical brain regions are discussed further below. L4/L5 IT neurons also frequently projected to ipsilateral association areas such as RSPagl. L4/L5 IT neurons, compared to L2/3 and L4 IT types, also more frequently contacted ACA and CP, as well as other sensory-motor areas (e.g., MOs) and HPF, though the trend was not significant for the latter two regions (Extended Data Fig. 32).

We only identified a single neuron in each of the L5 IT-1 and L6 IT-1 MET-types. The L5 IT-1 neuron projected to HVAs, RSP, and contralateral VISp only. The L6 IT-1 neuron was unusual in that it remained mostly within ipsilateral visual areas. More neurons are needed to characterize the axonal projection of these two types. Finally, there were few L5/L6 IT Car3 neurons found in VISp (a larger number were located in lateral HVAs; see Fig. 5); each projected to VISp, VISl, VISrl, and VISal (Fig. 4e, g, arrow) and across connectional modules^28^ to contralateral SSp-bfd (Fig. 4e). Interestingly, these neurons are unique in that they have relatively symmetrical projection target strength across brain hemispheres (as assessed by average total axon length, Fig. 4g, arrow).

L5 ET neurons had many interareal projection targets in common, regardless of MET-type, including multiple nuclei in the thalamus, superior colliculus, and the pretectal region (Fig. 4e, g). However, L5 ET-1 Chrna6 neurons on average had significantly fewer projection targets per neuron than L5 ET-3 Stac (Fig. 4f). This was largely the result of fewer cortical targets (Extended Data Fig. 33, Extended Data Fig. 30). We only identified one L5 ET-2 neuron, which had projections similar to the the other ET types. L5 ET-3 Stac neurons were more likely to target thalamic structures (like LP) and hindbrain (Fig. 4e, Extended Data Fig. 32). Projections to hindbrain structures, though, were relatively rare within the L5 ET group (Extended Data Fig. 33), but when they did occur, neurons were located in deep L5 (Fig. 4e, soma depth bar, top).

L6 CT neurons had significantly fewer targets than L5 ET types. Within cortex, they rarely projected outside of VISp. In the thalamus, they most reliably targeted LGd-core and other thalamic nuclei (Extended Data Fig. 33). L6b neurons most consistently targeted ipsilateral HVAs and RSPagl. They rarely projected contralaterally, but when they did, they targeted non-VIS cortex, RSP and ACA (Fig. 4e, g, arrow; Extended Data Fig. 33). We are not aware of previous WNM studies of these neurons.

### Higher Visual Areas (HVAs)

To examine the extent to which the properties of MET-types vary across visual areas, we plotted HVA neurons in the same dendrite and local axon feature-derived UMAP that we presented in Fig. 5a, top. In this visualization, HVA neurons also grouped cleanly by MET-type and were found in regions of the UMAP that largely overlapped with VISp neurons. For some MET-types (L4 IT, L4/L5 IT, L5 ET-1 Chrna6, and L5 ET-3 Stac), HVA neurons appeared to be concentrated within a subregion of each MET-type domain in the UMAP, suggesting their local morphologies differed from VISp neurons (Extended Data Fig. 34, Extended Data Fig. 35). To understand the effect of specific HVA region on this distribution, we plotted neurons colored by their soma location (Fig. 5a, bottom). For MET-types where we had the most coverage across HVAs (e.g., L2/3 IT, L4/L5 IT), there were no obvious region-specific clusters. Of interest, though, L4/L5 IT neurons did appear to be roughly ordered across medial-anterior to lateral-posterior structures.

Among the HVA neurons, we identified six of the eight Patch-seq defined IT MET-types (missing only L5 IT-1 and L5 IT-2 Pld5) (Fig. 5b). Importantly, multiple neurons from lateral VIS structures mapped to L5/L6 IT Car3, facilitating additional morphological characterization of these unique neurons. We also identified two of the three ET MET-types (L5 ET-1 Chrna6 and L5 ET-3 Stac) along with the L6 CT and L6b types. Additionally, we identified several L5 NP neurons. Looking across brain regions, dendritic and axonal morphologies appeared to be fairly consistent within a MET-type, with some feature variation as mentioned above, and agreed with the dendritic phenotypes observed with Patch-seq (Fig. 5c).

In order to examine differences in axonal projections between VISp and HVAs, we plotted the binary projection target matrix ordered by total number of projection targets (Fig. 5d,e). For each MET-type, VISp neurons had the fewest projection targets. When MET-types were well represented across areas, we found that most MET-types in VISp had significantly fewer axonal targets per neuron than at least one other HVA (L4/L5 IT and L6b were exceptions) (Extended Data Fig. 34, Extended Data Fig. 35). L2/3 IT and L4 IT neurons in HVAs contacted more VIS regions on average compared to VISp (Extended Data Fig. 31). L2/3 IT and L4/L5 IT HVA neurons also more frequently targeted other sensory cortical regions (somatosensory, auditory), HPF, and multiple, contralateral cortical regions (Extended Data Fig. 32). Most HVA L4/L5 IT neurons targeted the striatum, while L2/3 IT and L4 IT neurons rarely did. Interestingly, there was an inverse relationship between the amount of local axon and the number of projection targets across VISp and HVAs (Fig. 5f), which could have interesting functional implications for local versus interareal information processing across structures. L5/L6 IT Car3 neurons in VISl and VISpor had the largest number of targets in the IT division, with multiple contralateral targets (Fig. 5g, far right, bottom histogram).

Given the projection differences between HVA neurons and VISp neurons, we looked at whether there were transcriptomic differences as well between the HVA and VISp neurons. Using the Patch-seq data set, we trained a logistic regression model based on gene expression to predict HVA vs VISp location for the three most populous MET-types (L5 ET-3 Stac, L6 CT, and L4/L5 IT; see Methods). The models on average did not outperform chance predictions of VISp vs HVA location (50% chance accuracy based on equal subsampling; 95% confidence intervals for test accuracy: [50%, 75%] for L5 ET-3 Stac, [39%, 66%] for L6 CT, [28%, 66%] for L4/L5 IT), which suggested that there were not dramatic transcriptomic differences between VISp and HVA neurons for these MET-types. However, our ability to detect subtler differences may be limited by our relatively small number of HVA neurons in each MET-type.

In analyzing contralateral projections, we found that L4/L5 ITs and L5/L6 Car3 ITs had the largest proportions of contralateral-projecting neurons across MET-types (Extended Data Fig. 36a). We then compared the total axon length of the contralateral projections between these two MET-types and found that Car3 neurons had significantly more axon in contralateral cortex (Extended Data Fig. 36b,c). Furthermore, the laminar distribution patterns of these two MET-types differed markedly: L4/L5 IT neurons strongly innervated L5 and L6 (VISp and HVAs) and L1 (e.g., VISal, VISl) compared to Car3 neurons, which seemed to avoid L1 (Extended Data Fig. 36d).

L5 ET neurons also increased in projection target complexity from VISp to the HVAs (Fig. 5h,i). When these neurons had somas in anterior structures (e.g., VISa, VISal, and VISam), they had more midbrain and hindbrain targets. HVA L5 ET-3 Stac neurons contacted additional targets in VIS as well as other sensory cortical regions (Extended Data Fig. 33). Similarly, L6 CTs in VISpm and VISpor targeted additional VIS targets and other sensory thalamic nuclei (e.g., PO) relative to VISp neurons; they also more consistently targeted LP (Extended Data Fig. 33).

VISp L6b neurons had significantly more local axon than L6b neurons in HVAs (Extended Data Fig. 34), although they had similar total axon lengths. Looking at their laminar innervation patterns, we found that L6b neurons, regardless of soma location, consistently innervated cortical L1 across brain regions and hemispheres (Extended Data Fig. 36 d). L6b innervation of contralateral L1 differed again from the contralateral laminar innervation patterns described above for L4/L5 ITs and L5/L6 IT Car3 neurons. There were no significant differences in the number of targets across VIS areas for L6b neurons. As target number increased (independent of brain region), other sensory cortical areas (SS, AUD) were more likely to be targeted (Extended Data Fig. 33).

The few NP (”near projecting”) neurons in the WNM dataset were located only in HVAs and had projections restricted to VIS. Each neuron contacted 3 to 6 regions, which was low compared to other types. VISp was the only region targeted by all NP neurons.

### Linking multimodal properties across datasets

Within and across MET-types, local morphological and electrophysiological properties were highly correlated with gene expression gradients (Figs. 2 and 3). Using a morphological classifier, we could reliably link MET-type identity (and the corresponding morphoelectric-transcriptomic properties) to the complete axonal phenotype of excitatory neurons (Figs. 4 and 5). In doing this, we identified distinct patterns of interareal projections across MET-types, but still found considerable variation among the sets of targets of individual neurons belonging to the same MET-type. We hypothesized that the transcriptomic PCs that are correlated with local dendritic and electrophysiological properties might also be related to axonal properties, including projection targets.

To test this, we derived a transcriptomic correlated dendritic PC for each MET-type from the dendritic features that were highly correlated with transcriptomic variation (Fig. 6a; see also Extended Data Figs. 19 and 25). For example, soma depth and apical and basal dendrite overlap were highly correlated with the transcriptomic variation captured by L2/3 IT Tx PC-2 (Fig. 6b). Thus, these features were used to calculate the dendritic PC for this type; the same approach was applied to the other major excitatory subclasses. When we ordered dendrites by their transcriptomic-correlated dendritic PC, we saw clear differences in soma location and/or dendritic complexity across the continuum (Fig. 6b-e, top panels). Furthermore, we confirmed that our calculated transcriptomic-correlated dendritic PCs and matching Tx PCs were highly correlated for each Patch-seq subclass (Fig. 6b-e, bottom panels). Our goal in calculating transcriptomic-correlated dendritic PC for the WNM data set was to transfer this relationship across data sets and allow us to relate transcriptomic gradients to axonal properties (Fig. 6b-e, middle and bottom panels).

**Figure 6:**
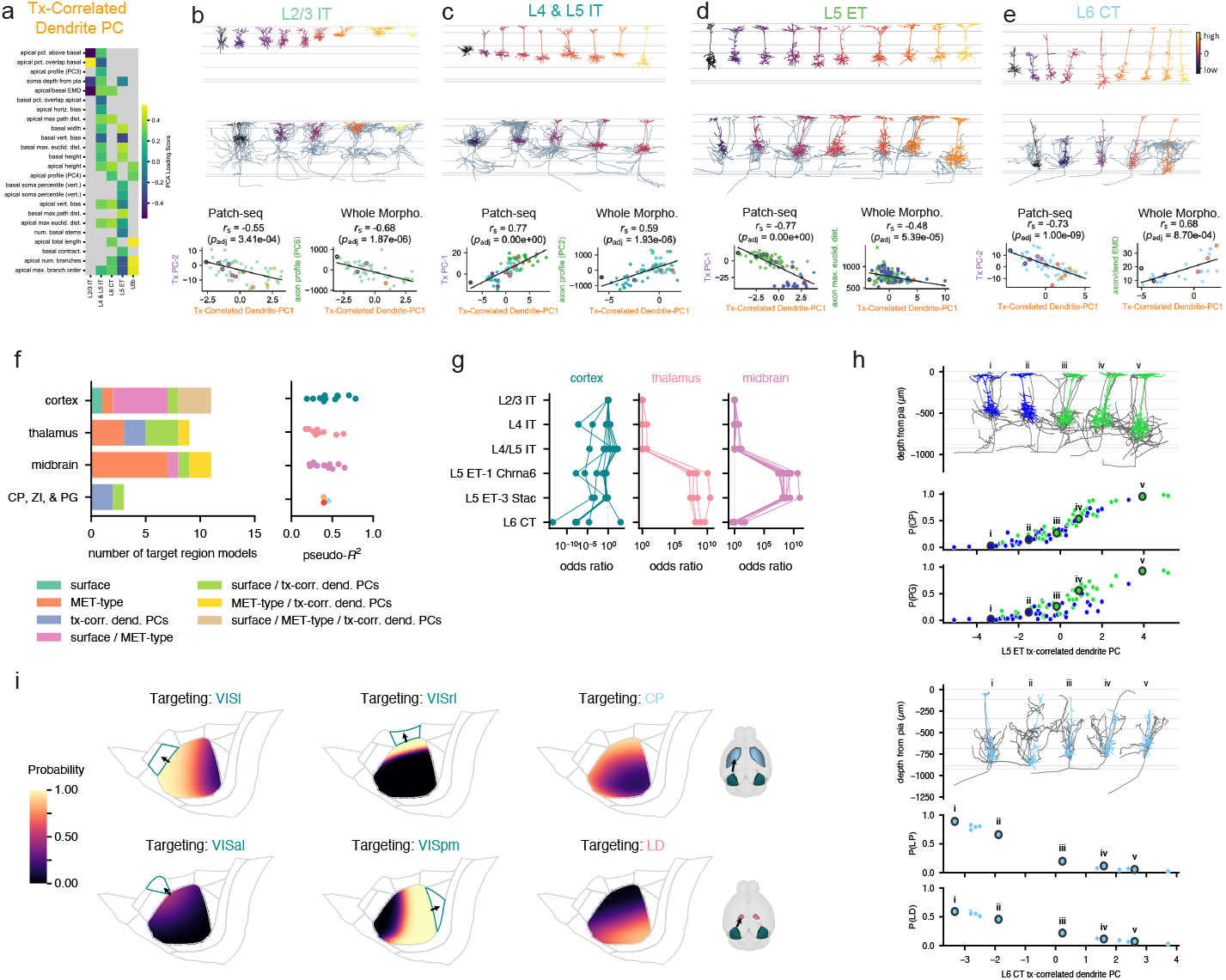
Projection target prediction using multimodal properties. **a**, Dendritic features that were highly correlated with the transcriptomic PCs described in Figs. 2 and 3 were used to calculate a dendritic PC for each MET-type. **b-e**, Morphologies from Patch-seq and WNM ordered and colored by dendritic PC value. Correlations between dendritic PC and transcriptomic PC are shown for Patch-seq neurons. Correlations between dendritic PC and local axon features are shown for WNM neurons. **f**, Types of logistic regression models used to predict whether WNM neurons project to specific targets (left) and pseudo-*R*^2^ values for the selected models (right). Models were selected by Akaike information criterion (Methods). **g**, MET-type odds ratios for models that used MET-type to predict projection targets. Higher odds ratios represent higher probabilities of projection associated with those MET-types. Odds ratios were defined relative to L2/3 IT (always set to 1). **h**, Effects of transcriptomic-correlated dendritic PCs on projection probabilities. Higher values of the L5 ET dendritic PC were associated with a higher chance of projecting to CP and PG. Lower values of the L6 CT dendritic PC were associated with a higher chance of projecting to LP and LD thalamus. **i**, Effects of cortical surface location on projections to different cortical and subcortical targets.

To test whether integrating multimodal properties of individual neurons could be used to predict their specific projection targets, we used logistic regression with the VISp WNM data set to model the probability of projection to each target area based on predicted MET-type, cortical surface location, and transcriptomic-correlated dendritic PCs (or subsets of those features, see Methods, Extended Data Figs. 37 and 38). We could identify models that outperformed a null model for each target region (Methods), and different regions could be best predicted by different combinations of features (Fig. 6f, left).

For example, nearly all cortical area targets were best predicted by models that made use of the cortical surface location (though in most cases they also made use of other features), while most midbrain targets were best predicted by models that made use of MET-type. These models did not explain all the variance in projection targets (estimated by pseudo-*R*^2^ ranging from 0.17 to 0.78, mean = 0.41, Fig. 6f, right, Methods), but examining the model properties could still provide insights into the relationships between projection targets and various cell features.

For models that used MET-type, the relative weights given to different types aligned with expectations about the types of neurons projecting to those targets (Fig. 6g). Cortical area models gave more similar weights to most MET-types (apart from L6 CT, which was overall less likely to project to those targets). Thalamus models predicted much higher projection probabilities for L5 ET and L6 CT neurons versus IT neurons, and midbrain models predicted high projection probabilities only for L5 ET neurons. Models that used transcriptomic-correlated dendritic PCs highlighted relationships between local morphology and projection targets. For example, deeper L5 ET cells with more complex tufts were predicted as being more likely to project to CP and PG (Fig. 6h). Shorter L6 CT cells were predicted to be less likely to project to the LP and LD thalamic nuclei (Fig. 6h). For most cortical targets, cells nearer to the target areas had a much higher probability of projecting there than more distant cells (Fig. 6i), which is consistent with the sections of visual space most well-represented by those HVAs^47^. For subcortical targets where cortical surface location was informative for the predictions, cells located in anterior VISp were predicted to be more likely to project to CP but less likely to project to LD thalamus (Fig. 6i).

Using these models, we predicted the probabilities that different MET-types would project to target regions across a range of VISp cortical locations and transcriptomic-correlated dendritic PC values (Fig. 7, Extended Data Fig. 39). For example, the predicted probability that L4/L5 IT MET-type neurons project to cortical targets was strongly dependent on the proximity of the neuron to those regions; in contrast, variations in the morphology along the dendritic PC axis had smaller effects on the projection probabilities (Fig. 7a). The predicted probability that L5 ET-3 Stac neurons project to cortical targets also strongly depended on their locations within VISp, although the overall probabilities were lower than L4/L5 IT neurons (Fig. 7b, left). The locations within VISp also strongly affected the probability of projecting to CP but had more modest effects on thalamic and midbrain projection probabilities. The projections to CP were also strongly influenced by the morphology of the L5 ET-3 Stac neuron (Fig. 7b, right; see also Fig. 6h), as was the probability of projecting to several other regions. We summarize the effects of location and transcriptomic-correlated dendritic PC variation in Figure 7c across the six MET-types used to fit the models. These prediction ranges illustrated the neuronal properties that were more important for accurately predicting their potential projections.

**Figure 7:**
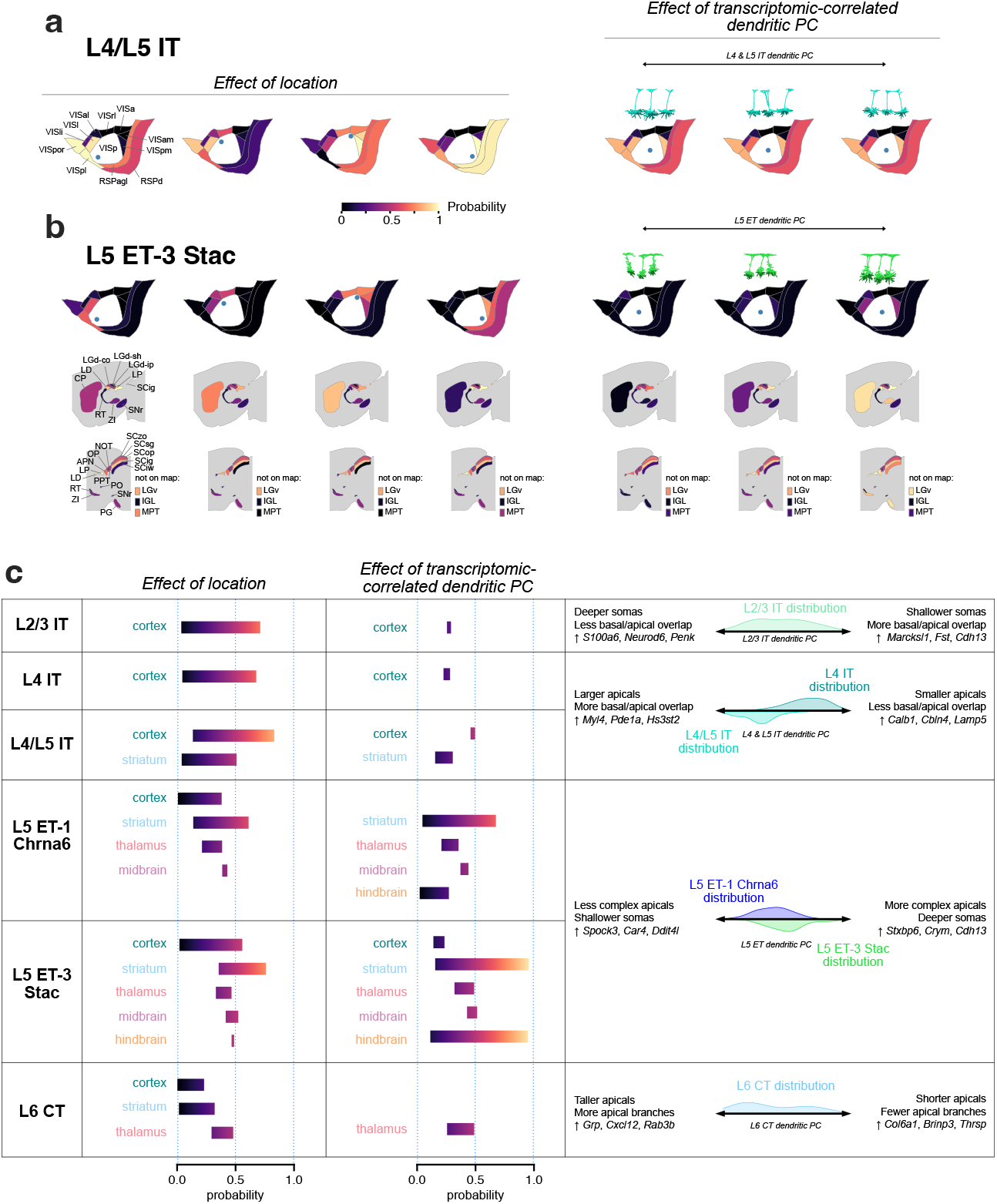
Predicting target projections for MET-types. **a**, Predicted probabilities of projecting to cortical targets for L4/L5 IT neurons. Probabilities are predicted at different VISp locations (left) and for different transcriptomic-correlated dendritic PC values (right). Example neurons (right, above) were chosen at low, medium, and high values of the dendritic PC range for projection probability calculations. **b**, Predicted probabilities of projecting to cortical and subcortical targets for L5 ET-3 Stac neurons. The effects of VISp location and dendritic PC are shown as in (**a**). **c**, Summary of the effects of location and dendritic PC variation by MET-type. Probability ranges for major structures (cortex, striatum, thalamus, midbrain, and hindbrain) were calculated by averaging the lowest and highest predictions across the regions belonging to the structure, either by varying location (second column) or dendritic PC (third column). The rightmost column summarizes the morphological features that vary along the relevant dendritic, the genes inferred to vary by the corresponding Tx PC, and the distributions of dendritic PC values by MET-type.

## Discussion

The relationship between the thousands of transcriptomically defined cell types and other phenotypic properties, including long-range axonal projections, has yet to be described for the vast majority of neurons in the mouse and human brain. Here we used the Patch-seq method to collect a large dataset of mouse visual cortical, excitatory neurons (*n* = 1544). We characterized the morphoelectric and transcriptomic properties of these neurons, mapped them to an established transcriptomic taxonomy^12^, and defined 16 excitatory MET-types. Using those types, we interrogated cross-modal relationships within and across types, in particular those between transcriptomic gradients and other phenotypic variation.

To extend MET-type descriptions to include local and long-range axonal properties, we reconstructed a dataset of 306 visual cortical neurons from whole brain images generated as part of the BICCN^27,31^, then built a multi-step classifier based on projection subclass identity (either transcriptomically or anatomically-defined), and dendritic features common to both datasets, to assign MET-type to WNMs. We characterized the complete axons of predicted MET-types and used a computational model to identify transcriptomically-related phenotypic properties that could best predict specific axonal projection targets.

### Relationships between T-types and MET-types

From our Patch-seq MET-type definitions, we found that multiple T-types were either consolidated into a single MET-type or mapped one-to-one to a MET-type. T-types were consolidated when there was considerable overlap or continuous variation in morphoelectric properties across those types. However, the presence of multiple T-types within a MET-type does suggest transcriptomic heterogeneity within a MET-type, which led us to examine how this variation corresponded with differences in the other modalities measured by the Patch-seq technique (Figs. 2 and 3). Transcriptomic variation (as quantified by transcriptomic principal components) often was correlated with electrophysiological and morphological feature variation, but this does not necessarily translate into strict separability along these feature dimensions between T-types as T-types can be distributed within transcriptomic space in complex ways (Extended Data Fig. 16a). However, characterizing the relationships between transcriptomics, electrophysiology, and morphology, as we have done here, allows us to understand how the properties interrelate in MET-types and T-types.

We also noted transcriptomic variation associated with neuronal activity (Extended Data Fig. 24a-c), which appeared to be responsible for a few T-type definitions in L2/3 IT cells and L6 IT cells, since removing those dimensions led to the merging of T-types in each of those subclasses (Extended Data Fig. 24d). Though recent work has suggested that L2/3 IT T-types may have functional differences^48,49^, two of them (L2/3 IT VISp Rrad and L2/3 IT VISp Adamts2) are primarily distinguished by differential expression of several activity-regulated genes, which could reflect transient variation in activity levels and be subject to experimental perturbation. Indeed, the Patch-seq technique appears to elevate the expression of several of those genes (Extended Data Fig. 24e-f) resulting in proportionately more cells mapping to the higher-activity T-type (L2/3 IT VISp Rrad) than in the reference data set. Other transcriptomic variation within L2/3 IT cells, though, did correlate with electrophysiological and morphological feature differences (Fig. 2a-f), such as action potential shape, depth within the cortex, and the shape of the dendritic arbor. This graded transcriptomic and spatial variation is consistent with that described by others^50^; for example, the gene *Cdh13*, found by those authors to be expressed more strongly by L2/3 IT neurons near the pia, had strong weights in both Tx PC-2 (correlated with AP shape) and Tx PC-3 (correlated with VISpm/VISal projection signature), which were both also correlated with depth in the cortex.

### Characterization of distinctive MET-types

Excitatory MET-types that mapped one-to-one with a T-type, like L5/L6 IT Car3 and L5 ET-1 Chrna6 neurons, were particularly interesting candidates for cross-data set integration. In our Patch-seq experiments, L5/L6 IT Car3 neurons look like deep spiny stellate neurons, usually found in L4, which are typically thought to be a local neuron^38^. However, the WNM data showed that these neurons have the most elaborate long-range projections within the IT group 5. Many of these neurons also have extensive contralateral projections to multiple VIS structures.

Neurons of another one-to-one type, L5 ET-1 Chrna6, are wider thick-tufted neurons that, compared with L5 ET-3 Stac neurons, project to a similar set of targets but with lower probability. Unlike L5 ET-3 Stac neurons, however, the L5 ET-1 Chrna6 cells rarely fire bursts of APs. Differences in bursting could have interesting functional implications for these deep brain projecting neurons (e.g.,^51^). These different ET types were first characterized transcriptomically by Tasic *et al.* [12], and our study is the first detailed description of their interareal projections. In motor cortex, different L5 ET types have been demonstrated to have functional differences^30^, and it will be interesting to determine if that is also true for VIS L5 ET types.

The multiple L5 IT MET-types identified with Patch-seq will also be interesting to investigate further with respect to WNM. There were hints that L5 IT-1 and/or L5IT-2 Pld5 may have very different axonal profiles than the L4/L5 IT neurons, but additional data are needed. With advances in whole brain imaging, data processing, and automated reconstruction, much more WNM data will become available in the near future^52,53^.

These more novel T-types have already been identified in multiple cortical regions and may be considered worthy of consideration in the canonical cortical circuit, especially L5/L6 IT Car3, which provides strong projections to contralateral cortex and may play an important role in interhemispheric, multi-sensory processing.

### Cross-modality predictions

We used logistic regression to model how well multi-modal cell properties could predict specific projection targets. We found that the location of the projecting cell within the cortex was important for some targets (e.g., HVAs, CP), while the MET-type of the cell and/or its transcriptomically-correlated dendritic properties were important for others (e.g., PG). The MET-type-specific, transcriptomically-correlated dendritic PC creates a useful link between the transcriptomic taxonomy and dendritic properties, which helps to narrow the transcriptomic space into which we can map neurons from other data sets. In addition, neurons on one side of certain dendritic continua can display preferential targeting compared to the other (Fig. 6h, Fig. 7), suggesting that there could be genes that encode and/or give access to “MET-P” types (MET-types that project to specific brain regions). Genes given high weights in the transcriptomic principal component analysis (Extended Data Fig. 16b, Fig. 7c) could be potential candidates for that: for example, CP-targeting L5 ET Stac cells might be selected by identifying cells of that type with high *Cdh13* or *Crym* expression. These analyses thus allow us to go beyond transcriptomic descriptions of major projection subclasses (e.g., ET, IT, CT) by characterizing other factors that link them to specific projection targets.

### Future directions

In this study, we focused on interareal, axonal targeting patterns. In subsequent studies, axon terminal morphology and subregion innervation patterns will also be of great interest. This may help to refine functional domains of downstream target structures beyond what we have learned from bulk labeling approaches^36^.

The availability of an increasing number of T-type and subclass specific viral and transgenic tools^54,55^ is essential for validating our findings and for future whole brain morphology studies. These tools offer critical benchmarks for feature registration across datasets and can offset the need for difficult, post-hoc, spatial transcriptomic studies to directly test these relationships.

The flexibility and extensibility of the classifier needs further testing on other data sets, as well as other brain regions and species. Thus far, we have successfully mapped mouse visual cortex inhibitory Sst MET-types and inhibitory subclasses from Patch-seq to neurons in the MICrONS EM dataset^4,56^ and found that Sst MET-types have differential connectivity^57^. Ongoing studies at our and other institutes aim to extend these results to the entire MICrONS volume. It will be of particular interest to understand the relationship between MET-types and connectivity-defined types^58^.

Although our models of projection target probability characterized how several neuronal properties influence projection patterns in cortex, a high amount of variance in projection patterns remains unexplained for many target regions (see Fig. 6f). It is possible that additional variance could be related to developmental processes regulating these projections which are inaccessible in this study or could reflect stochastic processes that may underlie an individual neuron’s targeting choices. Future models built from larger data sets that more fully characterize projection pattern relationships may better explain the variance, as well.

### Concluding remarks

With this work, we add to existing data^13^ to produce a comprehensive taxonomy of neuronal morpho-electictranscriptomic (MET) types in mouse visual cortex. We extend these findings beyond previous Patch-seq studies of excitatory cortical neurons^15^ to characterize the local and long-range axons of excitatory MET-types. We also create computational approaches for morphology-based, cross-data set mapping and cross-modality predictions, including the relationship to long-range projections. The transcriptome provides one of the most powerful links for translating cell types across species. Wherever we can draw close associations between it and other phenotypic neuronal properties, it will help us to better understand cell types and their contribution to brain organization and function across animal species, including our own.

## Methods

### Animal care and use

Experimental procedures that involved the use of mice were all conducted with approved protocols in accordance with NIH guidelines. They were also approved by the Allen Institute for Brain Science Institutional Animal Care and Use Committee (IACUC). Mice were housed ≤ 5 mice per cage and were maintained on a 12-hour light/dark cycle, in a humidity- and temperature-controlled room with water and food available ad libitum.

### Transgenic mice and sparse labeling

Transgenic driver and reporter mice used in Patch-seq and WNM studies are listed in Supplementary Data Table 2. Characterization of the expression pattern of many of the transgenic mouse lines can be found in the AIBS Transgenic Characterization database (http://connectivity.brain-map.org/transgenic/search/basic). Many of the brains used for WNM studies were described in a previous manuscript^27^. Additional brains were sparsely and robustly labeled for WNMs studies using Supernova virus, which was provided as a generous gift of pAAV-TRE-fDIO-GFP-IRES-tTA (Addgene plasmid # 118026;http://n2t.net/addgene:118026; RRID:Addgene 118026) by Minmin Luo, and variants.

### Tissue processing and slicing procedure

For preparation of acute brain slices, adult male and female mice (ages P45-P70) were first fully anesthetized by 5% isoflurane inhalation. Intracardiac perfusion was then performed with 25-50 mL of ice-cold cutting artificial cerebrospinal fluid (ACSF; 0.5 mM calcium chloride (dehydrate), 25 mM D-glucose, 20 mM HEPES buffer, 10 mM magnesium sulfate, 1.25 mM sodium phosphate monobasic monohydrate, 3 mM myo-inositol, 12 mM N-acetyl-L-cysteine, 96 mM N-methyl-D-glucamine chloride (NMDG-Cl), 2.5 mM potassium chloride, 25 mM sodium bicarbonate, 5 mM sodium L-ascorbate, 3 mM sodium pyruvate, 0.01 mM taurine, and 2 mM thiourea (pH 7.3), which had been continuously bubbling with a mixture of 95%*O*_2_/5%*CO*_2_). 350 µm sections were sliced on a vibrating microtome (Compresstome VF-300 vibrating microtome, Precisionary Instruments or VT1200S Vibratome, Leica Biosystems), either coronally or at a 17° angle from the coronal plane. For visual cortex, this latter slice angle helps to maximize the integrity of neuronal processes. In order to optimize registration to the CCFv3, a block-face image was collected before each section was cut (Mako G125B PoE camera with custom integrated software). Immediately after slicing, brain slices were placed in warm (34 °C) oxygenated cutting ACSF for 10 minutes, then allowed to further recover in holding ACSF (2 mM calcium chloride (dehydrate), 25 mM D-glucose, 20 mM HEPES buffer, 2 mM magnesium sulfate, 1.25 mM sodium phosphate monobasic monohydrate, 3 mM myo-inositol, 12.3 mM N-acetyl-L-cysteine, 84 mM sodium chloride, 2.5 mM potassium chloride, 25 mM sodium bicarbonate, 5 mM sodium L-ascorbate, 3 mM sodium pyruvate, 0.01 mM taurine, and 2 mM thiourea (pH 7.3)), bubbling with a mixture of 95%*O*_2_/5%*CO*_2_ at room temperature until transferred to the microscope for recordings.

### Patch-clamp recording

Slices were bathed in warm (34 °C) recording ACSF (2 mM calcium chloride (dehydrate), 12.5 mM D-glucose, 1 mM magnesium sulfate, 1.25 mM sodium phosphate monobasic monohydrate, 2.5 mM potassium chloride, 26 mM sodium bicarbonate, and 126 mM sodium chloride (pH 7.3)) and continuously bubbled with 95% O_2_/5% CO_2_. The bath solution contained blockers of fast glutamatergic (1 mM kynurenic acid) and GABAergic synaptic transmission (0.1 mM picrotoxin). Thick-walled borosilicate glass (Warner Instruments, G150F-3) electrodes were manufactured (Narishige PC-10) with a resistance of 4–5 MΩ. Before recording, the electrodes were filled with ∼1.0 to 1.5 µL of internal solution with biocytin (110 mM potassium gluconate, 10.0 mM HEPES, 0.2 mM ethylene glycol-bis (2-aminoethylether)-N,N,N*^′^*,N*^′^*-tetraacetic acid, 4 mM potassium chloride, 0.3 mM guanosine 5*^′^*-triphosphate sodium salt hydrate, 10 mM phosphocreatine disodium salt hydrate, 1 mM adenosine 5*^′^*-triphosphate magnesium salt, 20 µg/mL glycogen, 0.5U/µL RNAse inhibitor (Takara, 2313A) and 0.5% biocytin (Sigma B4261), pH 7.3). The pipette was mounted on a Multiclamp 700B amplifier headstage (Molecular Devices) fixed to a micromanipulator (PatchStar, Scientifica).

Electrophysiology signals were recorded using an ITC-18 Data Acquisition Interface (HEKA). Commands were generated, signals processed, and amplifier metadata were acquired using MIES (https://github.com/AllenInstitute/MIES/), written in Igor Pro (Wavemetrics). Data were filtered (Bessel) at 10 kHz and digitized at 50 kHz. Data were reported uncorrected for the measured (Neher, 1992) liquid junction potential of −14 mV between the electrode and bath solutions. Prior to data collection, all surfaces, equipment and materials were thoroughly cleaned in the following manner: a wipe down with DNA away (Thermo Scientific), RNAse Zap (Sigma-Aldrich), and finally with nuclease-free water.

After formation of a stable seal and break-in, the resting membrane potential of the neuron was recorded (typically within the first minute). A bias current was injected, either manually or automatically using algorithms within the MIES data acquisition package, for the remainder of the experiment to maintain that initial resting membrane potential. Bias currents remained stable for a minimum of 1 s before each stimulus current injection.

To be included in the analysis, neurons needed to have a >1 GΩ seal recorded before break-in and an initial access resistance <20 MΩ and <15% of the *R*_input_. To stay below this access resistance cut-off, cells with a low input resistance were targeted with larger electrodes. For an individual sweep to be included, the following criteria were applied: (1) the bridge balance was <20 MΩ and <15% of *R*_input_; (2) bias (leak) current within ±100 pA; and (3) root mean square noise measurements in a short window (1.5 ms, to gauge high frequency noise) and longer window (500 ms, to measure patch instability) <0.07 mV and <0.5 mV, respectively.

After electrophysiological recording, the pipette was centered on the soma or placed near the nucleus (if visible). A small amount of negative pressure was applied (∼ −30 mbar) to begin cytosol extraction and to attract the nucleus to the tip of pipette. After approximately one minute, the soma visibly shrank and/or the nucleus was near the tip of the pipette. While maintaining negative pressure, the pipette was slowly retracted; slow, continuous movement was maintained while monitoring the pipette seal. Once the pipette seal reached >1 GΩ and the nucleus was visible on the tip of the pipette, the speed was increased to remove the pipette from the slice. The pipette containing internal solution, cytosol, and the nucleus was removed from pipette holder, and its contents were expelled into a PCR tube containing the lysis buffer (Takara, 634894). Metadata for all Patch-seq neurons including in this study are located in Supplementary Data Table 3.

### Electrophysiology feature analysis

Electrophysiological features were measured from responses elicited by short (3 ms) current pulses and long (1 s) current steps as previously described^13,21^. Action potentials (APs) were detected, and the threshold, peak, fast trough, and width (at half-height) were calculated for each AP along with the ratio of the peak upstroke dV/dt to the peak downstroke dV/dt (upstroke/downstroke ratio). Several voltage trajectories (the initial AP elicited by the lowest-amplitude current pulses and steps, the derivatives of those APs, and the interspike interval) were analyzed as previously. AP features across responses to long current steps were averaged in time bins and concatenated across step amplitudes; bins without APs had interpolated values from their neighbors. This was done for steps starting at a given cell’s rheobase and increasing at 10 pA intervals. Sweeps from intervals without data were interpolated from sweeps at neighboring intervals. Subthreshold responses to hyperpolarizing current steps were analyzed as before by downsampling to 10 ms bins and concatenating responses from different stimulus amplitudes (ranging from −90 pA to −10 pA). Sparse principal component analysis was performed separately on data from each of these categories (e.g., AP waveform, AP features across current steps), and sparse principal components (sPCs) that exceeded 1% adjusted explained variance were kept. This yielded 61 sPCs in total from twelve data categories. The components were z-scored and combined to form the reduced dimension electrophysiology feature matrix.

### cDNA amplification and library construction

For Patch-seq experiments, we reverse transcribed the collected nuclear and cytosolic mRNA, and sequenced the resulting cDNA using the SMART-Seq v4 method described in Tasic *et al.* [12]. We used the SMART-Seq v4 Ultra Low Input RNA Kit for Sequencing (Takara, 634894) to reverse transcribe poly(A) RNA and amplify full-length cDNA according to the manufacturer’s instructions. We performed reverse transcription and cDNA amplification for 20 PCR cycles in 0.65 mL tubes, in sets of 88 tubes at a time. At least 1 control 8-strip was used per amplification set, which contained 4 wells without cells and 4 wells with 10 pg control RNA. Control RNA was either Mouse Whole Brain Total RNA (Zyagen, MR-201) or control RNA provided in the SMART-Seq v4 kit. All samples proceeded through Nextera XT DNA Library Preparation (Illumina FC-131-1096) using either Nextera XT Index Kit V2 Set A-D (FC-131-2001,2002,2003,2004) or custom dual-indexes provided by IDT (Integrated DNA Technologies). Nextera XT DNA Library prep was performed according to manufacturer’s instructions except that the volumes of all reagents including cDNA input were decreased either to 0.4x or to 0.2× by volume. Each sample was sequenced to approximately 500,000 - 1 million reads.

### Sequencing data processing

Fifty-base-pair paired-end reads were aligned to mm10 GENCODE vM23/Ensembl 98 reference genome, downloaded from 10X cell ranger (refdata-cellranger-arc-mm10-2020-A-2.0.0). Sequence alignment was performed using STAR aligner (v2.7.1a) with default settings. PCR duplicates were masked and removed using STAR option “bamRemoveDuplicates.” Only uniquely aligned reads were used for gene quantification. Gene counts were computed using the R Genomic Alignments package^59^ summarizeOverlaps function using “IntersectionNotEmpty” mode for exonic and intronic regions separately. Exonic and intronic reads were added together to calculate total gene counts; this was done for both the reference dissociated cell data set and the Patch-seq data set. Data were analyzed as counts per million reads (CPM).

### Transcriptomic mapping and analysis

We followed the procedures previously used by Gouwens *et al.* [13] to assign transcriptomic types to Patch-seq neurons by mapping Patch-seq transcriptomes to a reference dataset of scRNA-seq transcriptomes obtained from dissociated cells collected by Tasic *et al.* [12]. We used the same reference taxonomy here as in Gouwens *et al.* [13], starting with the 24,411 dissociated cells from VISp and ALM regions and 4,020 differentially expressed genes from Tasic *et al.* [12], but keeping only neuronal cells from the VISp region and their corresponding T-types (13,464 cells encompassing 93 cell types). We note that the T-types and subclasses that used the “PT” nomenclature in the original study have been renamed “ET” here to be consistent with a recently generated whole-brain taxonomy^3^.

#### Mapping to the VISp reference taxonomy

We mapped the transcriptomes of Patch-seq samples to the reference taxonomy above using the methods described previously for inhibitory neurons^13^. Briefly, for each Patch-seq transcriptome, we traversed the reference hierarchical transcriptomic tree, computing the correlation of its expression of select marker genes at each branch point of the tree with the expression profile of the reference dissociated cell types below that branch point. We chose the more correlated branch and repeated the process until the leaves (i.e., T-types) of the hierarchical tree were reached. This procedure was bootstrapped with 100 iterations at each branch point using a random subsampling (70%) of markers and reference cells. We defined a mapping probability based on the fraction of times that a cell mapped to a leaf or node of the reference taxonomy. The T-type with the highest mapping probability was assigned to that Patch-seq cell.

#### Mapping to the whole brain reference taxonomy

We also mapped Patch-seq cells to a recently generated whole-brain taxonomy^3^ and examined the correspondence between transcriptomic types assigned from this taxonomy and the VISp-derived reference taxonomy. Here we used the Hierarchical Approximate Nearest Neighbor (HANN) method implemented in the scrattch-mapping package (https://github.com/alleninstitute/scrattch-mapping). This method involved traversing the WB taxonomy hierarchy, selecting offspring node-differentiating marker genes at each node, and finding the approximate nearest neighbor T-type using marker gene correlation as the distance metric.

#### Assessing mapping quality

As in Gouwens *et al.* [13], we evaluated the T-type mappings by considering the confidence with which a Patch-seq transcriptome mapped to one or more reference T-types, and the expected level of ambiguity between reference T-types. We classified mapping quality measures (based on the correlation and the Kullback-Leibler (KL) divergence between the mapping probability distributions of Patch-seq cells and the reference mapping probability distribution; see Gouwens *et al.* [13]) into “highly consistent,” “moderately consistent” and “inconsistent” categories. 1,544 excitatory neurons that passed our QC criteria for both electrophysiological and transcriptomic data were included in this study. 1,290 of these cells mapped to T-types with “high consistency”, a similar fraction to what we found for inhibitory neurons using the same method^13^. In this study, we excluded cells with inconsistent mapping from further analyses.

#### Visualization of reference cells and Patch-seq cells

For visual comparison of reference dissociated and Patch-seq cells, we selected the 7,339 dissociated FACS-sorted neurons from primary visual cortex from the Tasic *et al.* [12] reference data set that were within the glutamatergic branch of the hierarchy (32 T-types) and used 1,398 differentially expressed (DE) genes (the top 50 DE genes in each direction for all pairwise cluster comparisons within only those excitatory types). The log_2_(CPM + 1) values of these DE genes were combined across the Patch-seq and reference cells and reduced to 20 components with PCA. Three “technical bias” PCs were removed as they were found to be correlated (Pearson’s *r* =0.65, 0.45, and 0.45) with the collection method. We visualized the variation in the remaining 17 PCs in two dimensions using Uniform Manifold Approximation and Projection (UMAP)^37^.

#### Dimensionality reduction for continuous transcriptomic variation

Because Patch-seq transcriptomes are known to suffer from increased contamination and gene dropout^11,13,60^, we defined transcriptomic dimensions from reference dissociated cells collected from mouse visual cortex. For each transcriptomic subclass (L2/3 IT, L4 & L5 IT, L6 IT, L5/L6 IT Car3, L5 ET, L5 NP, L6 CT, and L6b), we identified highly variable genes using Brennecke’s method (https://github.com/AllenInstitute/scrattch.hicat/). We then performed PCA and omitted PCs with a tolerance below 0.01 (i.e., PCs with standard deviations ≤ 0.01 times the standard deviation of the first principal component), which resulted in 3–7 principal components per subclass. We then projected data from Patch-seq cells assigned to the different MET-type groups into this lower dimensional space using the gene loadings from PCA.

#### Gene ontology analysis of gene modules

To validate the transcriptomic dimensions derived from PCA and infer biological functions from the continuous variation within transcriptomically-defined subclasses, we performed a dimensionality reduction with weighted gene co-expression network analysis (WGCNA)^41^. Unlike PCA, WGCNA identifies modules of genes that are co-expressed, which could be advantageous for gene ontology analysis. We projected the transcriptomic data from Patch-seq cells assigned to their respective MET-type groups into both the PCA and WGCNA spaces defined from reference cells and computed the Spearman correlation of each principal component and WGCNA-derived eigengene for that group of cells. We found that many of the PCA-derived transcriptomic dimensions were highly correlated with at least one eigengene (Extended Data Figs. 17 and 18).

To find functional associations with transcriptomic principal components, we performed a gene ontology analysis on the set of genes in any gene module that was correlated with a transcriptomic PC (> 0.85 Spearman correlation) using g:Profiler with g:SCS multiple testing^61^. We applied a significance threshold of 0.05. For visualization purposes, we dropped associations with very large terms (> 5, 000 associations), which tended to be non-specific in nature; we also limited the number of displayed associations to at most 15 terms (Extended Data Figs. 17 and 18).

#### Calculation of VISpm-projecting vs VISal-projecting transcriptomic signature

In order to examine whether the genes identified as differentially expressed between VISpm-projecting and VISal-projecting L2/3 cells^42^ were related to other cellular properties, we projected L2/3 IT Patch-seq data into a PC space derived from these genes. We first identified the cells in the Kim *et al.* [42] study with the best transcriptomic quality, selecting cells identified as “L23 AL” or “L23 PM” with “highly consistent” or “moderately consistent” quality when mapped to the same reference VISp taxonomy as Patch-seq cells. We limited analysis to genes with a log fold change > 1 and adjusted p-threshold < 0.05, combining the genes from both Zinbwave-EdgeR or Zinbwave-DESeq2 analyses in the study. We then performed PCA, reducing the log-transformed expression of the resulting 838 differentially expressed genes in 345 upper cortical layer neurons to 20 features (total explained variance = 0.21). We projected Patch-seq data mapping to L2/3 IT T-types onto this common PC space for further comparison of cellular properties with this HVA projection-associated transcriptomic signature.

#### Differential gene expression analysis

Differentially expressed ion channels were identified using the scrattch.hicat package (https://github.com/AllenInstitute/scrattch.hicat/) as previously described^12^, except that the proportions of cells expressing the gene in each type were not required to differ by ≥ 0.7, as we did not want to limit our identified genes to only those expressed in an on/off manner. Only genes that were identified as being differentially expressed in both the reference and Patch-seq data sets were included.

### Morphological reconstruction

#### Biocytin histology

Neurons were filled with biocytin via the patch pipette. To visualize the label, a horseradish peroxidase (HRP) enzyme reaction using diaminobenzidine (DAB) as the chromogen was used after the electrophysiological recording. 4,6-diamidino-2-phenylindole (DAPI) stain was also used identify cortical layers as described previously^21^.

#### Imaging

Slices from Patch-seq experiments were mounted on slides and imaged as described previously^21^. Briefly, operators captured images on an upright AxioImager Z2 microscope (Zeiss, Germany) equipped with an Axiocam 506 monochrome camera and 0.63x optivar lens. Two-dimensional tiled overview images were also captured (Zeiss Plan-NEOFLUAR 20X/0.5) in brightfield transmission and fluorescence channels. Higher resolution image stacks of individual cells were acquired in the transmission channel only for the purpose of morphological reconstruction. Light was transmitted using an oil-immersion condenser (1.4 NA). High-resolution, multi-tile image stacks were captured (Zeiss Plan-Apochromat 63x/1.4 Oil or Zeiss LD LCI Plan-Apochromat 63x/1.2 Imm Corr) at an interval of 0.28 µm (1.4 NA objective) or 0.44 µm (1.2 NA objective) along the Z axis. Image tiles were stitched in ZEN software and exported as single-plane TIFF files.

#### Anatomical location of Patch-seq cells

Layer and anatomical location were determined based on DAPI stained overview images mentioned above. The soma position of reconstructed neurons, as well as the pia, white matter, and L1–L6b borders (using DAPI for reconstructed neurons) were drawn and used in subsequent analyses. Individual cells were manually aligned to the Allen Mouse Common Coordinate Framework version 3 (CCFv3) by matching the overview image of the slice with a “virtual” slice at an appropriate location and orientation within the CCFv3. Laminar locations were calculated by finding the path connecting pia and white matter that passed through the cell’s coordinate, identifying its distance to pia and white matter as well as position within its layer, then aligning those values to an average set of layer thicknesses.

#### Computer-assisted morphological reconstruction of Patch-seq neurons

Dendritic reconstructions were performed for a subset of neurons with good quality transcriptomics, electrophysiology, and labeling. Reconstructions were generated based on 63X image stacks described above. Stacks were run through a Vaa3D-based image processing and reconstruction pipeline^62^. An automated reconstruction of the neuron was produced using TReMAP^63^. Alternatively, initial reconstructions were created manually using the reconstruction software PyKNOSSOS (Ariadne-service) or through the citizen neuroscience game Mozak (Mozak.science)^64^. Automated or manually initiated reconstructions were then extensively manually corrected and extended using a range of tools (e.g., virtual finger, polyline) in the Mozak extension (Zoran Popovic, Center for Game Science, University of Washington) of Terafly tools^65,66^ in Vaa3D. Where possible, the local axon was also reconstructed. After 3D reconstruction, morphological features were calculated as previously described^13,21^.

#### Automated morphological representations

All neurons that were eligible for reconstruction were automatically segmented and post-processed to produce a quantifiable neuron reconstruction using the approach described in Gliko *et al.* [53]. These automated reconstructions were used to make inferred MET-type assignments for cells from T-types that split across different MET-types (see below).

### MET-type definition

We defined MET-types from our Patch-seq data set as previously described^13^. Briefly, we first used electrophysiological and morphological features to define ME-clusters by several methods and defined consensus clusters from the combined results^13,21^. We used the per-cell cross-T-type mapping probabilities (see “Mapping to the reference data set” above) and cross-ME-cluster mapping probabilities (by subsampled random forest classification) to construct the edges of a graph in which the nodes represented cells with specific T-type/ME-type combinations. We then used the Leiden community detection algorithm^67^ to group strongly-connected nodes into MET-types (see Extended Data Figure 11b). The analysis was performed on the 384 neurons with electrophysiological data, transcriptomic data, and morphological features from a manually-curated reconstruction.

We observed that nearly all T-types were strongly associated with a single MET-type; therefore, T-types were used to infer MET-type labels for an additional 1,090 Patch-seq neurons that lacked a manually-curated reconstruction. For the handful of T-types that split across two MET-types (L6 IT VISp Penk Col27a1, L6 IT VISp Penk Fst, L6 IT VISp Col23a1 Adamts2, L5 ET VISp Lgr5), we used morphological features from automated morphological reconstructions to assign the final MET-type label (78 neurons from those T-types lacked an automated reconstruction and therefore were not assigned an inferred MET-type).

### fMOST imaging

As described previously^27^, resin-embedded, GFP-labeled brains underwent chemical reactivation to recover GFP fluorescence and facilitate wide-field or two-photon block-face imaging^34,68^. For the entire mouse brain, a 15–20 TB dataset containing 10,000 coronal planes of 0.2 to 0.3 µm X-Y resolution and 1 µm Z sampling rate was generated within 2 weeks. Tissue was prepared and imaged as previously described^69,70^. A 40X water-immersion lens with NA 0.8 was used to provide an optical resolution (at 520 nm) of 0.35 µm in XY axes and voxel size of 0.35 x 0.35 x 1.0 µm, appropriate for neuron reconstruction. GFP was imaged with an excitation wavelength of 488 nm and a bandpass emission filter of 510–550 nm.

### Whole neuron morphology (WNM) reconstruction and analysis

Vaa3D-TeraVR was used for WNM reconstructions of fMOST images. All dendrites and the complete local and long-range axonal arbor was traced using the virtual finger or polyline tool. Special care was taken to mark all putative axonal terminals, which were identified based on a large, well-labeled bouton, for secondary review by an experienced annotator. For this quality control step, the entire reconstruction was reviewed using Tera-VR. At high magnifications, the axon proximal to the soma or the main branches of distal axon collaterals were carefully examined for missed branches. Post-processing steps were run on completed reconstructions to ensure that there were no errors (i.e., breaks or loops).

#### fMOST image registration to CCF

Whole brain fMOST images were registered to the average mouse brain template of CCFv3 by one of two methods: BrainAligner^27^ or DeepMAPI^71^. For the DeepMAPI method, reconstruction data was supplied for several neurons labeled across individual brains and registration was performed iteratively as previously described. For the BrainAligner method, in brief, images were down-sampled by 64 × 64 × 16 (X, Y, Z), and outer contours were affine-aligned using the Robust Landmark points Matching algorithm (RLM). Intensity was then normalized by matching the local average intensity of raw fMOST images to that of the CCFv3, and local alignment was then iteratively deformed. As a final step, mBrainAligner was used, as necessary, to manually or semi-automatically adjust the boundaries of brain regions. With either method, once images were CCF-aligned, the reconstructed neurons were transformed into the CCFv3 space using the generated deformation fields. DeepMAPI-registered reconstructions were used in the WNM analyses presented throughout the paper. With this method, VIS WNM projection targets largely agree with what has previously been described in the literature for population studies. Where there are differences, they may result from issues with registration accuracy, particularly for smaller structures (e.g., SCig), differences in the location and/or type of neurons labeled, and/or the type of labeling method used.

#### Calculating the WNM projection matrix

CCF-registered reconstructions were translated such that all somas were positioned in the left hemisphere. SWC files were subsequently resampled to ensure uniform spacing between nodes. To quantify the pattern of axonal projection targets, a projection matrix was derived based on reconstruction node counts per anatomical target structure (see structure list below). Target regions were represented in both ipsilateral and contralateral hemispheres. To reduce the size of the projection matrix, only regions containing a branch or tip node were included.

#### Nomenclature and abbreviations in the Allen Mouse Brain Common Coordinate Framework, version 3 (CCFv3) ontology of brain regions referred to in this study

**Isocortex**: frontal pole (FRP), primary motor area (MOp), secondary motor area (MOs), primary somatosensory area (SSp), supplemental somatosensory area (SSs), gustatory area (GU), visceral area (VISC), dorsal auditory area (AUDd), primary auditory area (AUDp), posterior auditory area (AUDpo), ventral auditory area (AUDv), primary visual area (VISp), anterolateral visual area (VISal), anteromedial visual area (VISam), lateral visual area (VISl), posterolateral visual area (VISpl), posteromedial visual area (VISpm), laterointermediate area (VISli), postrhinal area (VISpor), anterior cingulate area, dorsal part (ACAd), anterior cingulate area, ventral part (ACAv), prelimbic area (PL), infralimbic area (ILA), orbital area, lateral part (ORBl), orbital area, medial part (ORBm), orbital area, ventrolateral part (ORBvl), agranular insular area, dorsal part (AId), agranular insular area, posterior part (AIp), agranular insular area, ventral part (AIv), retrosplenial area, lateral agranular part (RSPagl), retrosplenial area, dorsal part (RSPd), retrosplenial area, ventral part (RSPv), posterior parietal association area (PTLp), anterior area (VISa), rostrolateral visual area (VISrl), temporal association area (TEa), perirhinal area (PERI), ectorhinal area (ECT).

**Olfactory areas (OLF)**: piriform area (PIR).

**Hippocampal formation (HPF)**: hippocampal region (HIP), fields CA1, CA2, CA3, dentate gyrus (DG), entorhinal area, lateral part (ENTl), entorhinal area, medial part (ENTm), parasubiculum (PAR), postsubiculum (POST), presubiculum (PRE), subiculum (SUB), prosubiculum (ProS).

**Cortical subplate (CTXsp)**: claustrum (CLA), endopiriform nucleus, dorsal part (EPd), endopiriform nucleus, ventral part (EPv), lateral amygdalar nucleus (LA), basolateral amygdalar nucleus (BLA), basomedial amygdalar nucleus (BMA).

**Cerebral nuclei (CNU)**: caudoputamen (CP), nucleus accumbens (ACB), fundus of striatum (FS), central amygdalar nucleus (CEA), medial amygdalar nucleus (MEA), globus pallidus, external segment (GPe), globus pallidus, internal segment (GPi), bed nuclei of the stria terminalis (BST). STR-unspecified (STR) corresponds to areas of the striatum that have not be assigned to a child structure.

**Thalamus (TH)**: ventral anterior-lateral complex (VAL), ventral medial nucleus (VM), ventral posterolateral nucleus (VPL), ventral posterolateral nucleus, parvicellular part (VPLpc), ventral posteromedial nucleus (VPM), ventral posteromedial nucleus, parvicellular part (VPMpc), posterior triangular thalamic nucleus (PoT), medial geniculate complex, dorsal part (MGd), medial geniculate complex, ventral part (MGv), medial geniculate complex, medial part (MGm), lateral geniculate complex, dorsal part (LGd), lateral posterior nucleus (LP), posterior complex (PO), anteromedial nucleus (AM), interanterodorsal nucleus (IAD), lateral dorsal nucleus (LD), mediodorsal nucleus (MD), submedial nucleus (SMT), paraventricular nucleus (PVT), nucleus of reuniens (RE), central medial nucleus (CM), paracentral nucleus (PCN), central lateral nucleus (CL), parafascicular nucleus (PF), reticular nucleus (RT). Thalamus-unspecified (TH) corresponds to areas of the thalamus that have not be assigned to a child structure.

**Hypothalamus (HY)**: subthalamic nucleus (STN), zona incerta (ZI).

**Midbrain (MB)**: substantia nigra, reticular part (SNr), midbrain reticular nucleus (MRN), superior colliculus, motor related (SCm), periaqueductal grey (PAG), anterior pretectal nucleus (APN), red nucleus (RN), pedunculopontine nucleus (PPN), dorsal nucleus raphe (DR). Midbrain-unspecified (MB) corresponds to areas of the midbrain that have not be assigned to a child structure.

**Pons (P)**: parabrachial nucleus (PB), pontine grey (PG), pontine reticular nucleus, caudal part (PRNc), tegmental reticular nucleus (TRN), pontine reticular nucleus (PRNr), locus ceruleus (LC). Pons-unspecified (P) corresponds to areas of the pons that have not been assigned to a child structure.

#### Generating local morphology

To analyze the local axon of the WNM, axon nodes that were more than 500 µm from the soma in the x-z dimensions were excised. Any orphaned segments were also removed. In 115 cells, a fraction of superficial axon nodes were registered outside of the cortex. To correct this, the cell was translated along the streamline passing nearest to the soma until the stopping criterion was met. The stopping criterion was that either all superficial axon nodes were in the cortex or that the soma was at the L6b–white matter boundary.

#### Calculating morphological features in WNM

To extract local features for WNM data, a CCF driven protocol was developed to replicate the patch-seq laminar annotations. A 2-dimensional slice was drawn through the CCF which passed through a given cell’s soma. The slice was drawn such that it minimized the curvature of the cortex at both the pial and white matter surfaces. Cortical layers were annotated on the 2-dimensional slice using the CCF structure annotations.

From here, morphological features were extracted as outlined earlier for Patch-seq cells. However, in this study, we did not differentiate between apical and basal dendrites when calculating interaction features with local axons, such as the percentage of overlap.

Summary features were derived from the projection matrix, including the total number of projection targets, total projection length, total length in visual cortex, total length in ipsilateral visual cortex, the length of axon within the soma structure, total number of targets in visual cortex, total number of targets in contralateral visual cortex, and proportion of the total axon length within the soma structure.

#### Morphological feature alignment

Reconstructions from the WNM data set were uprighted and cut to imitate the slicing that occurs when preparing Patch-seq samples from visual cortex. The slice thickness and average soma depth-in-slice for patch-seq were 350 µm and 48.2±12.6 µm, respectively. To imitate this, WNM reconstructions were positioned 48.2 µm into a 350 µm wide rostral-caudal bounding box. Any dendrites extending beyond this rostral-caudal bounding box were excised. (Step 1a Extended Data Fig. 15).

To further align the Patch-seq and WNM data sets, the chamfer distance was minimized between two-dimensional feature point clouds. For each feature, a depth by feature point cloud was created for each data set. These point clouds were subsequently aligned by imposing a linear transformation on the feature, which reduced the chamfer distance between the respective point clouds. (Step 1b Extended Data Fig. 15).

#### Multi-step MET-type prediction

A systematic multi-step approach was developed to predict MET-types within the WNM dataset. The method aimed to first predict a projection subclass label—namely IT-NP-L6b, ET, or CT—and subsequently channel the data to a specialized classifier that exclusively predicted MET-types based on a designated subclass.

The first step involved predicting projection subclass (IT-NP-L6b, ET and CT) using dendritic morphology features. Training data was aggregated from Patch-seq and WNM neurons. Patch-seq subclass labels were derived explicitly from MET-type labels. A projection-derived subclass was found for WNM using high-level projection patterns (Step 2 Extended Data Fig. 15). Together, the Patch-seq and WNM data were shuffled and split into training and testing sets (80%/20%, respectively). The support vector classifier achieved 96% prediction accuracy on the hold-out data set using a radial based kernel function, balanced class weight, and a *C* value of 1. Final dendrite-derived projection subclass labels were predicted for WNM using a leave-one-out approach (Step 3 Extended Data Fig. 15).

In WNM, when projection-derived subclass and dendrite-derived subclass labels were aligned (*n* = 293), cells were routed to the corresponding MET-type classifier. Discrepancies between projection and dendrite derived subclass labels (*n* = 12) were resolved using local axon features. For these twelve cells, silhouette scores were calculated to determine which subclass a cell’s local axon was most consistent with. If the silhouette analysis for the local axon matched either the projection- or dendrite-derived subclass (*n* = 11), cells were then routed to the appropriate MET-type classifier. One cell had a different subclass assignment at each step; this cell was routed to a MET-type classifier using the dendrite-derived subclass.

To predict ET MET-types, a random forest classifier with 25 estimators, a maximum depth of 7, balanced class weights, a minimum of 5 samples per split, and at least 2 samples per leaf node was used. 5-fold cross validation was repeated 20 times and achieved a mean accuracy of 77.2% ± 11.5%. The cumulative confusion matrix was recorded (Step 4 Extended Data Fig. 15). L5 ET-3 Stac was randomly undersampled in each iteration of cross validation to reduce the impact of class size imbalance. For IT-NP-L6b MET-type classification, a random forest classifier with 250 estimators, a maximum depth of 10, balanced class weights, a minimum of 3 samples per split, and at least 4 samples per leaf node was used. 5-fold cross validation was repeated 20 times and achieved a mean accuracy of 91% ± 3.6%. The cumulative confusion matrix was recorded (Step 4 Extended Data Fig. 15). L6b was randomly undersampled in each iteration of cross validation to reduce the impact of class size imbalance. CT cells were directly mapped to the L6 CT MET-type.

Over the course of 500 iterations, the Patch-seq training data was sampled without replacement at 95% with selection probabilities proportional to the MET-type class size. During each iteration, a new classifier was trained on this sub-sampled data set, subsequently predicting MET-types for all WNM cells. The final MET assignment was determined based on the most frequently predicted MET-type label. Prediction probabilities are reported as the fraction of iterations a cell was assigned to the most frequently predicted MET-type label.

Cells predicted to a L6 IT MET-type but having a soma in L6b were reassigned to the L6b class (*n* = 2). A single cell, which exhibited notably sparse apical dendrite obliques given its overall local morphology, was initially categorized as L5 NP. However, due to its prominent long contralateral projections, it was reassigned to L4/L5 IT.

### Logistic regression models

Logistic regression was used to predict VISp vs. HVA locations for Patch-seq neurons using transcriptomic data. Highly variable genes were identified from the Patch-seq data set for the three most populous MET-types (L5 ET-3 Stac, L6 CT, L4/L5 IT). PCA was performed on those sets of genes, and the top 10 PCs were used as predictors for logistic regression. Since Patch-seq VISp neurons substantially outnumbered Patch-seq HVA neurons (L5 ET-3 Stac: *n*=298 VISp, *n*=64 HVA; L6 CT: *n*=255 VISp, *n*=55 HVA; L4/L5 IT: *n*=116 VISp, *n*=25 HVA), we repeatedly subsampled equally from both location types so that the chance accuracy level would be 0.5. We divided the HVA neurons with a 50% train/test split, then selected the same number of VISp neurons for training and test sets, as well. We then trained the regression model and calculated the test accuracy; we repeated this procedure 1000 times and calculated the 95% confidence interval of the mean accuracy across those repetitions.

The probabilities of individual VISp WNM neurons projecting to specific target regions was modeled by logistic regression. Only regions that were targeted by at least 10 VISp cells in the data set were used. For each region, binomial generalized linear models were fit using the cortical surface location, MET-type, and the subclass-specific transcriptomic-correlated dendritic PCs as predictors. Only predicted MET-types with at least 5 cells were included. Note that the transcriptomic correlated dendritic PCs were calculated for each class for every cell so that all cells would have the same set of predictor variables. Models were fit using all the predictors as well as using different subsets (seven model types in all). To select among the model types for each area, the Akaike information criterion (AIC) was calculated for each model, and the model with the lowest AIC was chosen, unless a simpler model using fewer predictors had a comparable AIC (i.e., the difference in the AIC was less than 2^72^). For each selected model, we calculated a pseudo *R*^2^ = 1 − log(*L*_model_)/ log(*L*_null_) where *L*_model_ was the likelihood of the data with the selected model and *L*_null_ was the likelihood of the data with a null model^73^ to estimate the variance explained by the selected model.

To estimate the effects of dendritic PC on target projection probability in Figure 6h, probabilities were calculated at the average cortical location of the neurons in the examined MET-type. For the effects of cortical location on target projection probability in Figure 6i, the probabilities were calculated for a given MET-type (L4/L5 IT for cortical targets, L5 ET-3 Stac for CP, and L6 CT for LD) and used the average dendritic PC values for neurons of that MET-type. For the effect of cortical location plots in Extended Data Figure 39, the probabilities were calculated using the MET-type associated with the highest odds ratio and used average dendritic PC values for all cells in the data set. Prediction error rates were estimated based on the predicted probabilities with a threshold of *p* = 0.5 and are reported both from the full training data set and from a leave-one-out cross validation (LOOCV) procedure.

### Statistics and research design

No statistical methods were used to predetermine sample sizes, but the sample sizes here are similar to those reported in previous publications. No randomization was used during data collection as there was a single experimental condition for all acquired data. The different stimulus protocols were not presented in a randomized order. Data collection and analyses were not performed blind to the conditions of the experiments as there was a single experimental condition for all acquired data.

Correlations were measured by the non-parametric Spearman rank correlation coefficient unless otherwise noted. Kruskal-Wallis tests followed by post hoc Dunn’s tests were used to identify significant differences across multiple groups. The p-values of multiple comparisons (e.g., correlations between all Tx PCs and electrophysiology/morphology features) were adjusted by the Benjamini-Hochberg method^74^ for an family-wise error rate of 0.05.

## Supporting information

Extended Data Figure 1 all_the_morphs5-5-9

Extended Data Figure 1 all_the_morphs0-4

## Data and software availability

Transcriptomic data supporting the findings of this study will be available at the NeMO archive upon publication. Similarly, electrophysiological data for this study will be available at the DANDI archive and morphological reconstructions from this study, as well as the fMOST whole brain images used for WNM reconstructions, will be available at the BIL archive.

The electrophysiology data acquisition software (MIES) used for this study is available at https://github.com/alleninstitute/mies. The morphological reconstruction software (Vaa3D-TeraFLY-Mozak is freely available at http://home.penglab.com/proj/vaa3d/home/index.html and the code is available at https://github.com/Vaa3D. The code for electrophysiological and morphological feature analysis and clustering is available as part of the open-source Allen SDK repository (https://github.com/AllenInstitute/AllenSDK), skeleton-keys repository (https://skeleton-keys.readthedocs.io/en/latest/), pyropractor repository (https://github.com/AllenInstitute/pyropractor/), IPFX repository (https://github.com/alleninstitute/ipfx), and DRCME repository (https://github.com/alleninstitute/drcme).

## Acknowledgments

We are grateful to the Transgenic Colony Management, Neurosurgery & Behavior, Lab Animal Services, Molecular Biology, Histology, and Imaging teams at the Allen Institute for technical support. We thank Nuno da Costa for helpful feedback on the manuscript. We thank Adrian Wanner and colleagues for providing reconstruction services through Ariadne and Roy Szeto for facilitating the reconstruction work contributed by Mozak.science. Finally, we thank the Mozak citizen-scientists for their valuable contribution. The research was funded by multiple grant awards from institutes under the National Institutes of Health (NIH), including award number, U01MH105982 from the National Institute of Mental Health and Eunice Kennedy Shriver National Institute of Child Health and Human Development, and U19MH114830 from the National Institute of Mental Health to H.Z and Q. Luo. The work was also funded by STI2030-Major Projects (2021ZD0201000 to H.G.) and NSFC grant (T2122015 to A.L.). The content is solely the responsibility of the authors and does not necessarily represent the official views of NIH and its subsidiary institutes. The Wenzhou Medical University reconstruction team received funding from the EPFL–Blue Brain Project. This work was also supported by the Allen Institute for Brain Science. The authors thank the Allen Institute founder, Paul G. Allen, for his vision, encouragement, and support.

