## Extended Data Figures 2-39 for "Connecting single-cell transcriptomes to projectomes in mouse visual cortex"

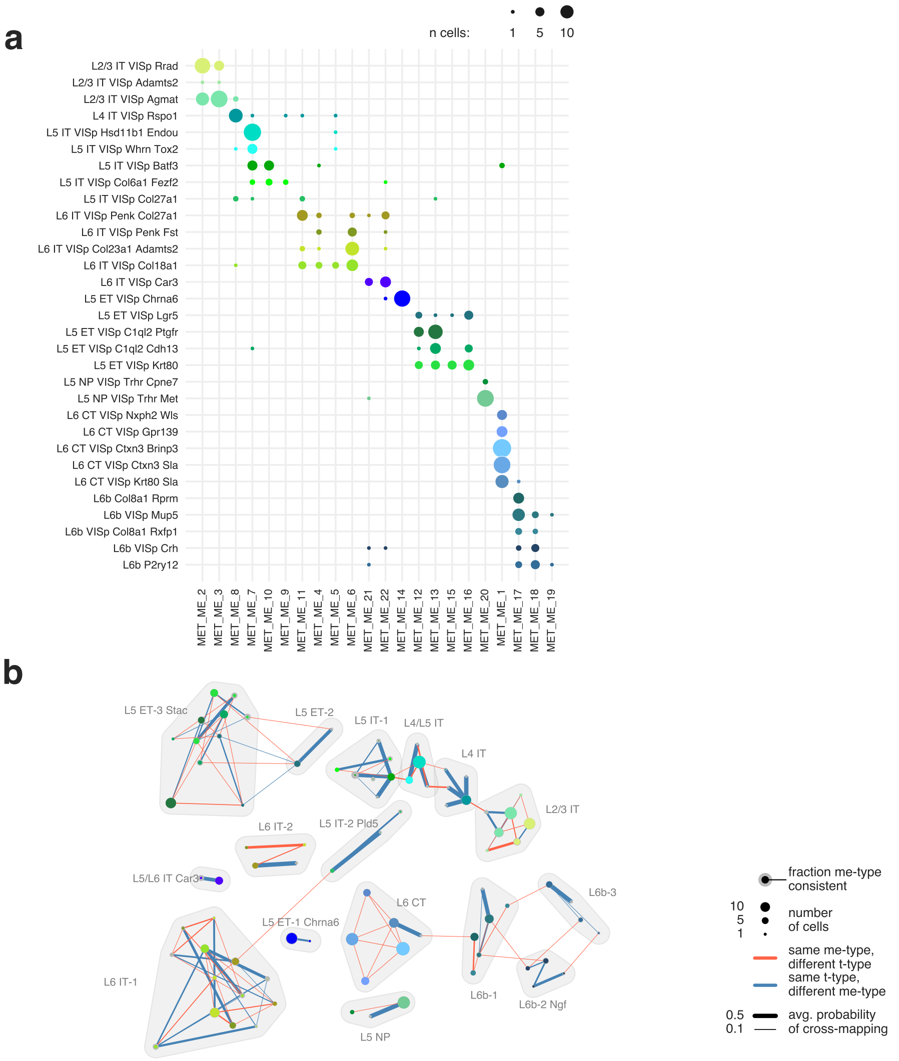


**Extended Data Figure 11: Definition of excitatory MET-types.** **a**, Relationship between T-types and ME-types. **b**, Community-detection grouping of T- and ME-type combinations into MET-types. Nodes are cells with a specific combination of ME-type and T-type (as in (**a**)). The size of the node indicates the number of cells with that combination of types. Edges represent cross-mapping probabilities of cells that could map to another T-type (orange line) or ME-type (blue line); thicker lines indicate higher probabilities. Shaded regions show the groupings of the ME-type/T-type combinations into MET-types.

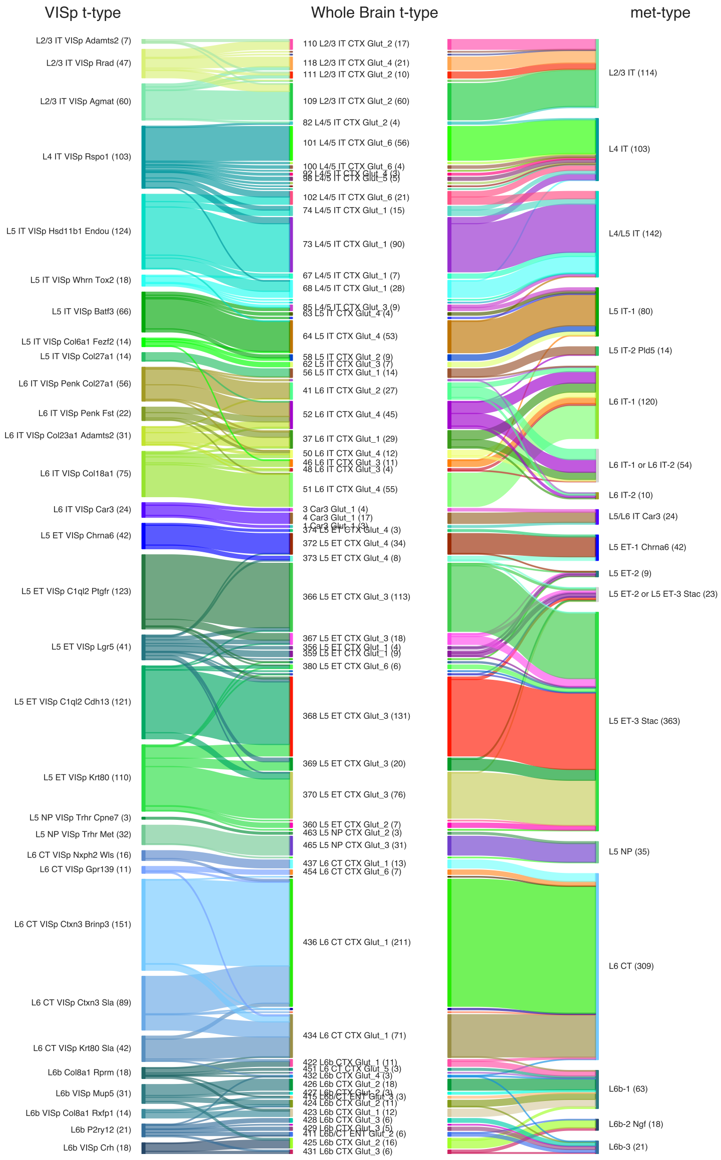


**Extended Data Figure 12:** Comparison of the mapping of Patch-seq neurons to the Tasic *et al.* [12] VISp taxonomy and to the Yao *et al.* [3] whole-brain taxonomy.

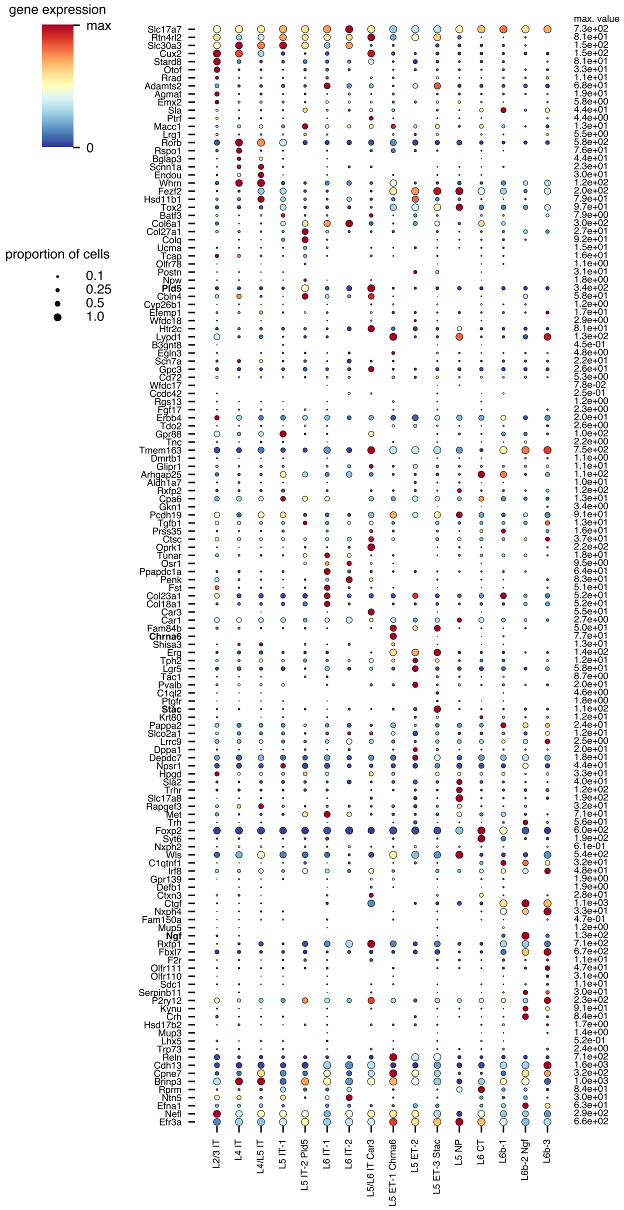

**Extended Data Figure 13: Marker gene expression across MET-types.** Color indicates average gene expression level within a MET-type (scaled to the maximum value, listed at right). Dot size indicates the proportion of cells within the MET-type that expressed the gene. The marker gene list for excitatory cortical neurons was taken from Extended Data Figure 5b in Tasic *et al.* [12]. Bold gene names indicate those used as part of MET-type names.

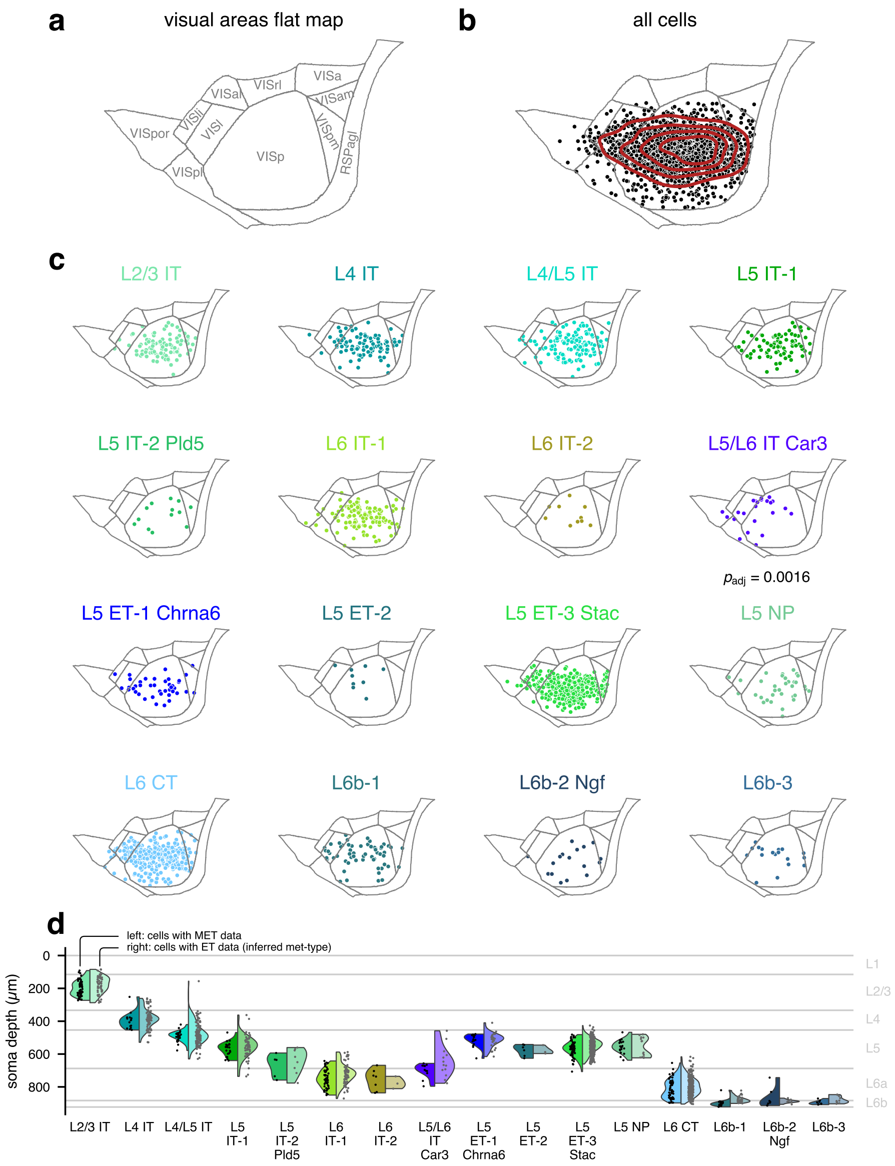
a
visual areas flat map
VISa
VISrl
VISal
VISam
VISli
VISl
VISpor
VISp
VISpm
RSPagl
VISpl
b
all cells
c
L2/3 IT
L4 IT
L4/L5 IT
L5 IT-1
L5 IT-2 Pld5
L6 IT-1
L6 IT-2
L5/L6 IT Car3
$p_{adj}$ = 0.0016
L5 ET-1 Chrna6
L5 ET-2
L5 ET-3 Stac
L5 NP
L6 CT
L6b-1
L6b-2 Ngf
L6b-3
d
left: cells with MET data
right: cells with ET data (inferred met-type)
soma depth (μm)
0
200
400
600
800
L1
L2/3
L4
L5
L6a
L6b
L2/3 IT
L4 IT
L4/L5 IT
L5 IT-1
L5 IT-2 Pld5
L6 IT-1
L6 IT-2
L5/L6 IT Car3
L5 ET-1 Chrna6
L5 ET-2
L5 ET-3 Stac
L5 NP
L6 CT
L6b-1
L6b-2 Ngf
L6b-3

**Extended Data Figure 14 *(preceding page)*: Surface and depth locations by MET-type.** **a**, Flat map visualization of visual cortical areas. **b**, Positions of all Patch-seq cells on the cortical surface. Red lines show contours of a kernel density estimate (KDE) of the total distribution. **c**, Distributions of MET-types on the cortical surface. A permutation test of the log-likelihoods for each MET-type distribution arising from the all-cell KDE (**b**) was used to estimate p-values. Only L5/L6 IT Car3 exhibited a significant difference from the all-cell distribution. **d**, Soma depth distributions of MET-types. Cortical depths were normalized to a standard set of layer thicknesses. For each MET-type, cells with all three data modalities (morphology, electrophysiology, and transcriptomics) are shown on the left, and those with only electrophysiology and transcriptomics data are shown on the right. The MET-types for the latter cells were inferred by T-type and, when available, auto-traced morphologies (see Methods).

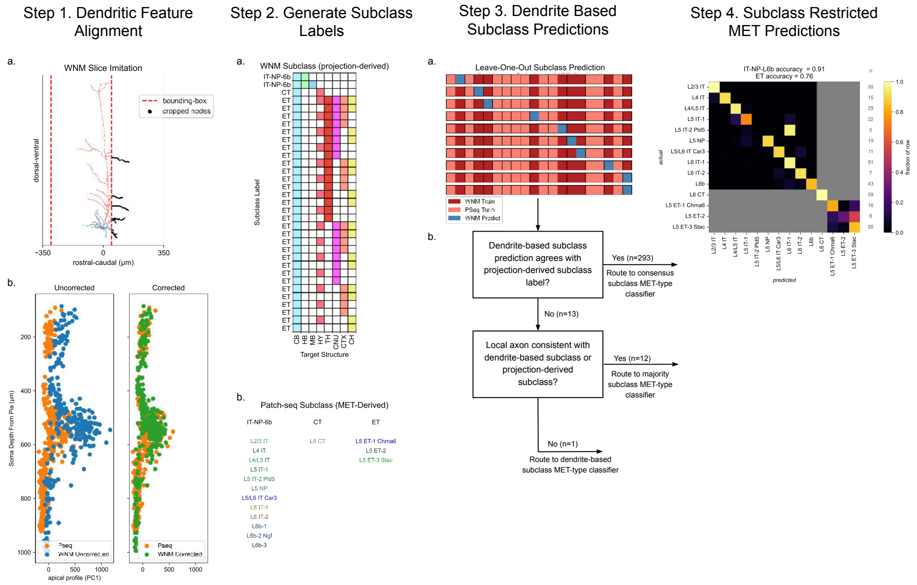


**Extended Data Figure 15: MET prediction workflow.** **Step 1a**, Uprighted WNM dendrites were cropped in the rostral-caudal dimension to replicate sample slicing in the Patch-seq data set. **Step 1b**, Features were aligned using chamfer distance minimization. For each feature, a depth by feature point cloud was created for each data set. These point clouds were subsequently aligned by imposing a linear transformation on the feature to minimize the chamfer distance between the respective point clouds. **Step 2a**, WNM projection subclass training labels (e.g., IT/NP/L6b, CT, ET) were derived from the projection targets of each individual neuron. **Step 2b**, Patch-seq projection subclass labels (e.g., IT/NP/L6b, CT, ET) were derived from MET-type labels (e.g., L5 ET-1 Chrna6). **Step 3a**, Leave-one-out subclass prediction protocol using dendrite morphology features. **Step 3b**, Flow chart for mapping samples to the correct MET-type classifier. **Step 4a**, Confusion matrix for MET-type classifiers. Cross classifier indices are grayed out.

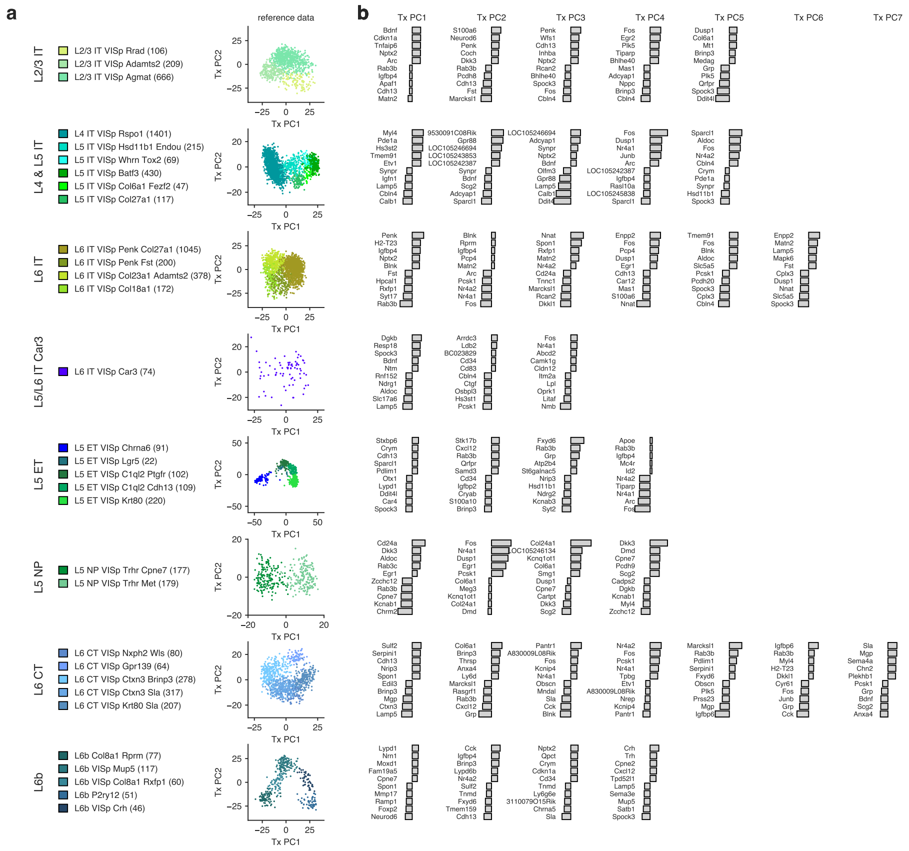

**Extended Data Figure 16: Continuous variation within transcriptomic subclasses.** **a**, Scatter plots showing values in the top 2 transcriptomic PC dimensions for reference FACS neurons within each subclass grouping. Cells are colored by their T-type. **b**, Bar plots of the top 5 positive and top 5 negative weights contributing to a given transcriptomic dimension derived from the subclass-specific PCA.

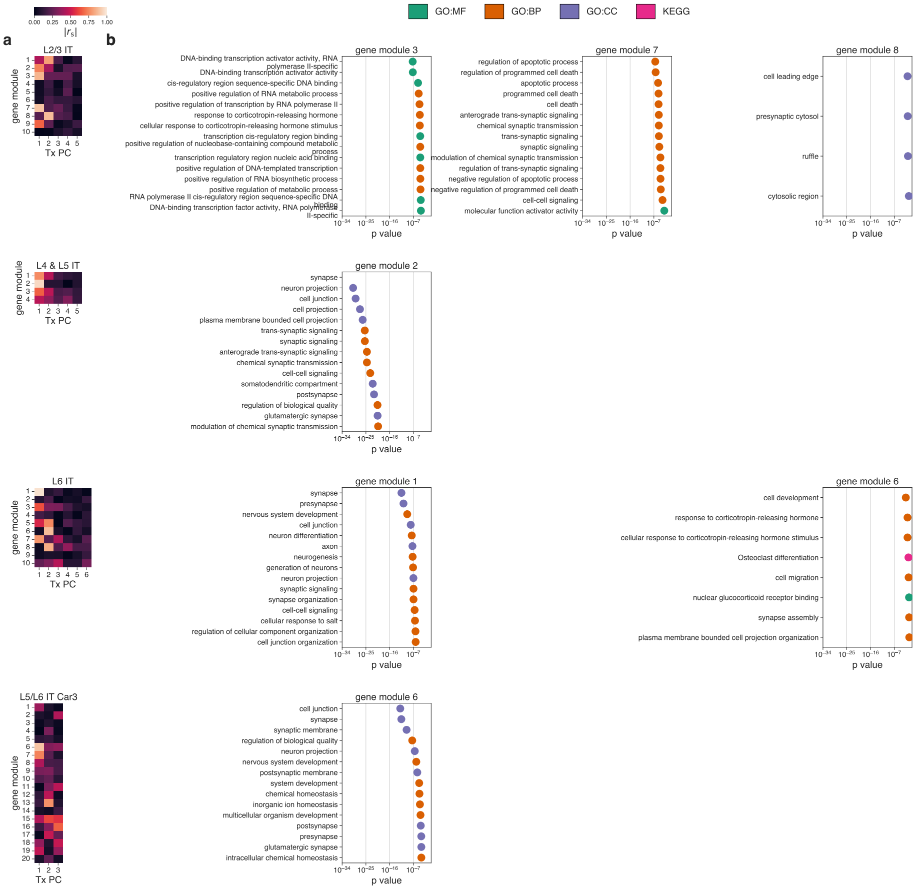


**Extended Data Figure 17: Functional associations of genes in WGCNA modules derived from IT subclasses.** **a**, Heat maps of Spearman correlations between WGNCA eigengenes and transcriptomic principal components. Both methods were performed on reference cells for each transcriptomic subclass. **b**, Gene ontology analyses were performed on gene modules that were highly correlated with a transcriptomic PC in (**a**). Dot plots show the 15 most significant terms from gene ontology analysis. Subontology (biological process, cellular component, molecular function, Kyoto Encyclopedia of Genes and Genomes (KEGG) pathway database) is indicated by color.

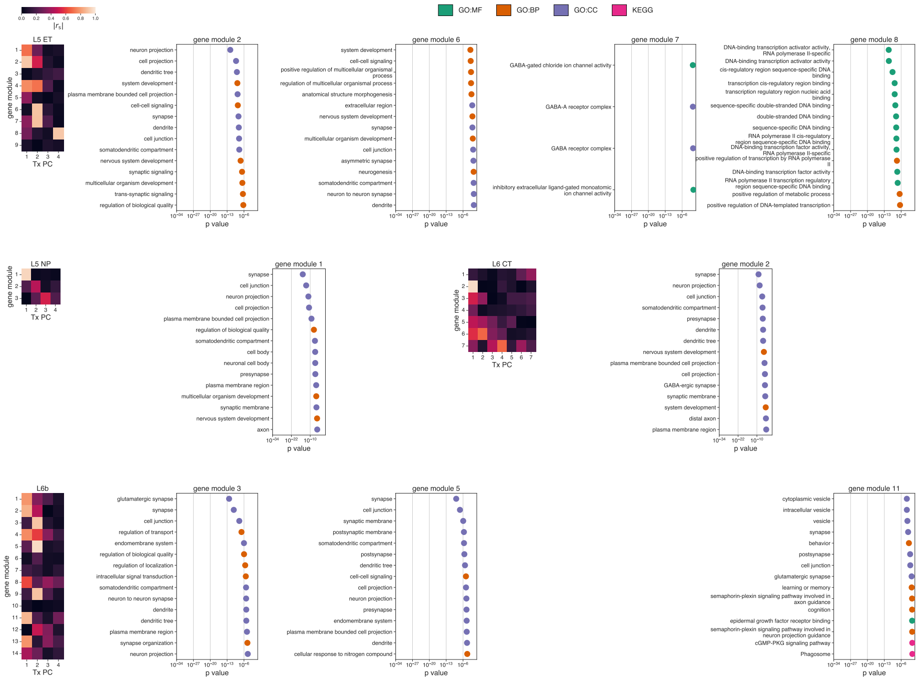


**Extended Data Figure 18: Functional associations of genes in WGCNA modules derived from ET, NP, CT and L6b subclasses.** Heat maps show Spearman correlations between WGNCA eigengenes and transcriptomic principal components. Both methods were performed on reference cells for each transcriptomic subclass. Gene ontology analyses were performed on gene modules that were highly correlated with a transcriptomic PC. Dot plots show the 15 most significant terms from gene ontology analysis. Sub-ontology (biological process, cellular component, molecular function, Kyoto Encyclopedia of Genes and Genomes (KEGG) pathway database) is indicated by color.

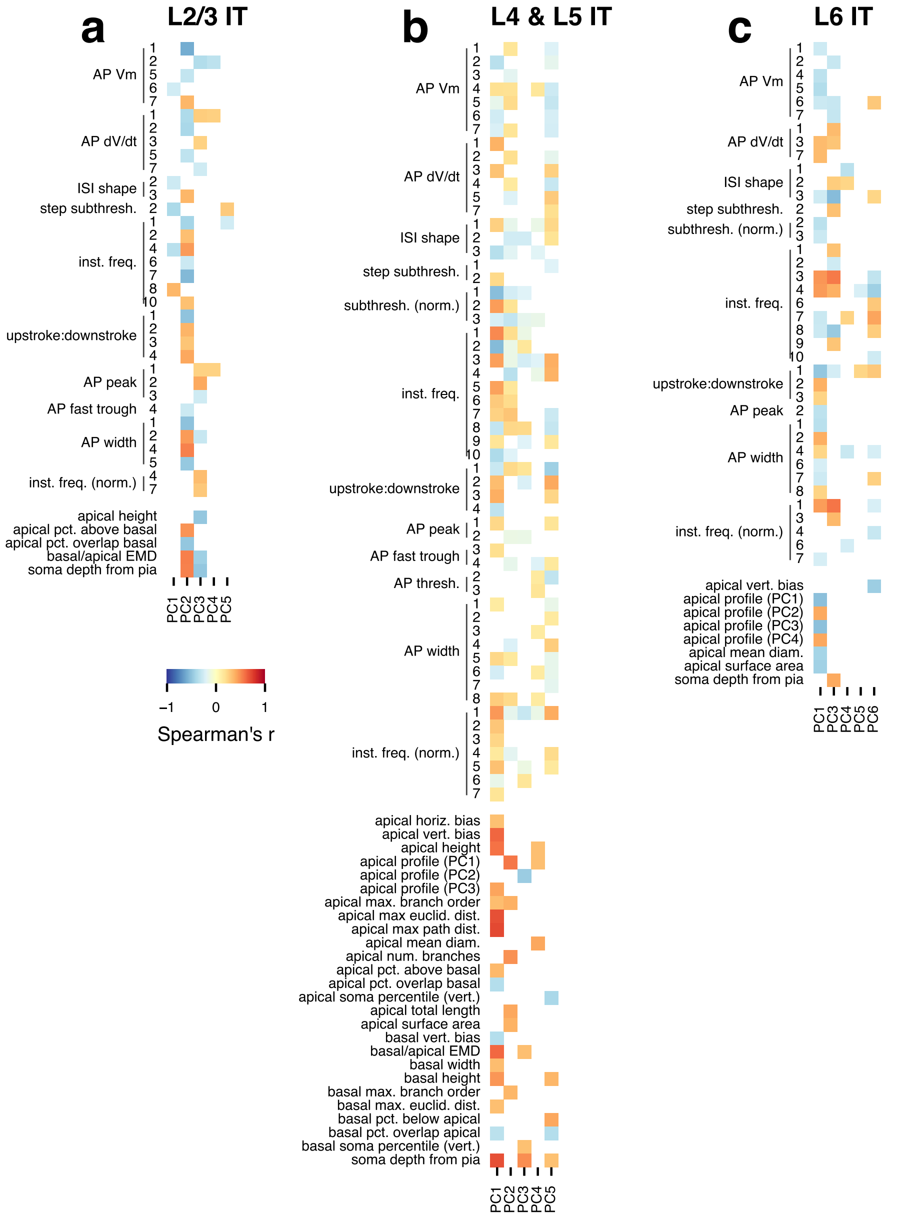


**Extended Data Figure 19: Cross-modal correlations within IT subclasses.** Statistically significant Spearman correlations between transcriptomic principal components (PCs) and electrophysiological (upper) and morphological (lower) features are shown in the heat maps for the L2/3 IT (**a**), L4 and L5 IT (**b**), and L6 IT (**c**) subclasses. Only PCs and features with at least one significant correlation are shown.

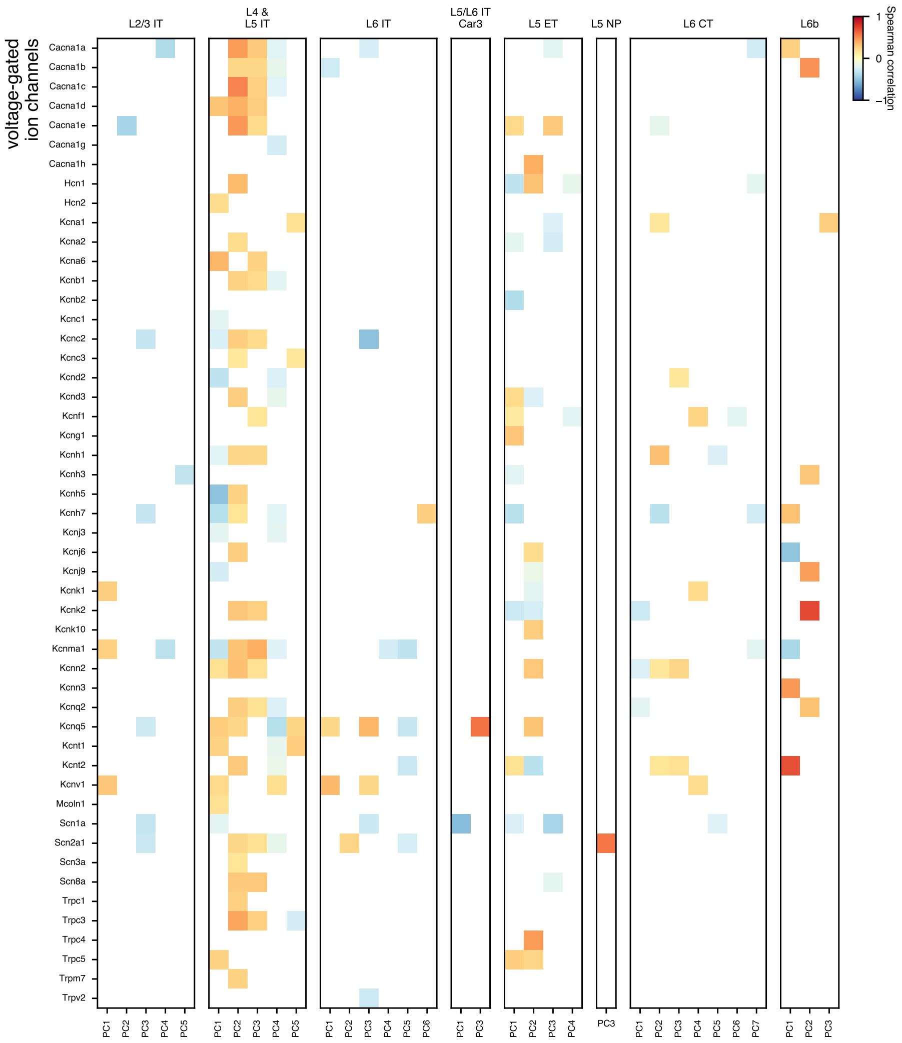


**Extended Data Figure 20: Correlations between voltage-gated ion channel gene expression and transcriptomic PCs.** Spearman correlations were measured between the transcriptomic PCs and voltage-gated ion channel gene expression (list from Földy *et al.* [17]). Correlations are shown for the Patch-seq data set. Only statistically significant correlations found in both the Patch-seq and reference FACS data sets are shown. Significance was assessed after multiple comparison correction of p-values. PCs and genes with no significant correlations are omitted.

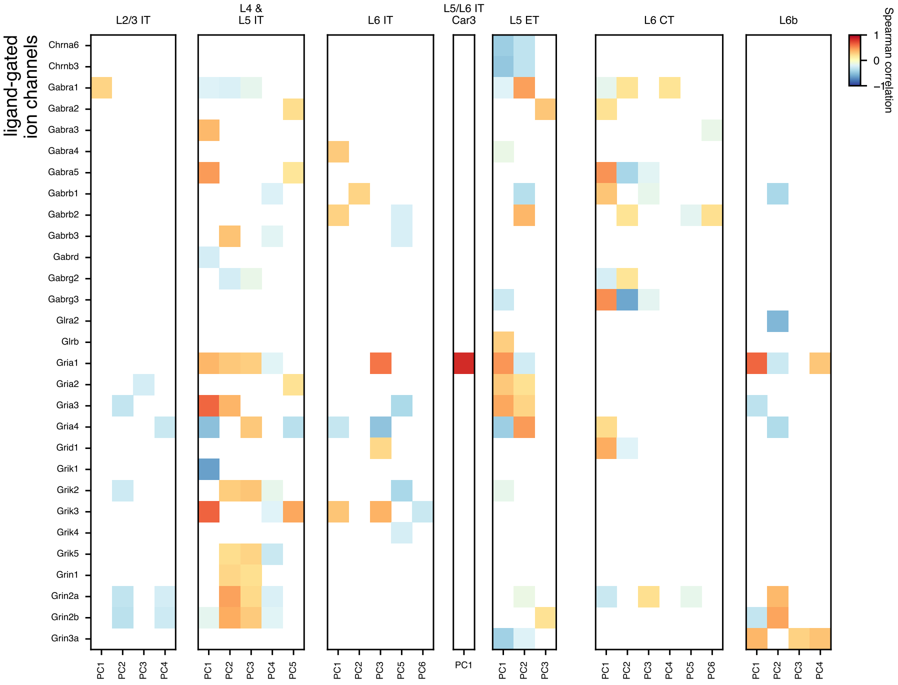


**Extended Data Figure 21: Correlations between ligand-gated ion channel gene expression and transcriptomic PCs.** Spearman correlations were measured between the transcriptomic PCs and ligand-gated ion channels gene expression (list from Földy *et al.* [17]). Correlations are shown for the Patch-seq data set. Only statistically significant correlations found in both the Patch-seq and reference FACS data sets are shown. Significance was assessed after multiple comparison correction of p-values. PCs and genes with no significant correlations are omitted.

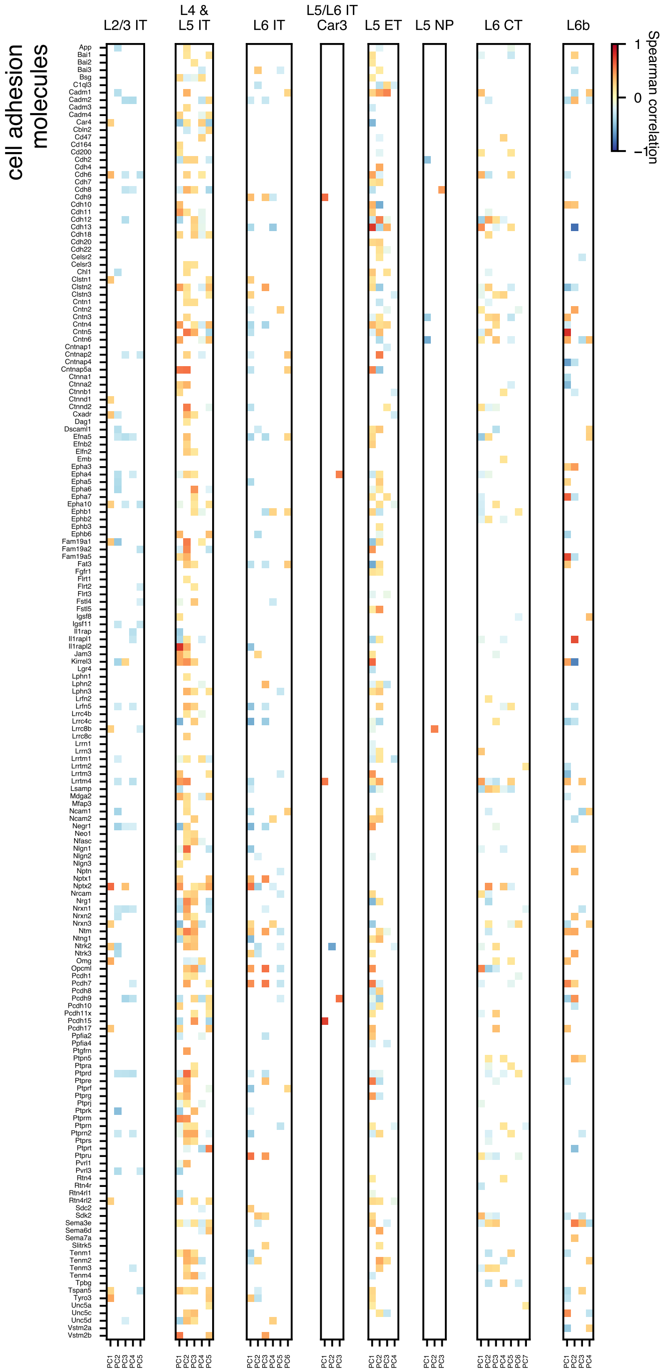
cell adhesion molecules
L2/3 IT
L4 & L5 IT
L6 IT
L5/L6 IT Car3
L5 ET
L5 NP
L6 CT
L6b
Spearman correlation
1
0
−1

**Extended Data Figure 22** ***(preceding page)*****: Correlations between cell adhesion molecule gene expression and transcriptomic PCs.** Spearman correlations were measured between the transcriptomic PCs and cell adhesion molecule gene expression (list from Földy *et al.* [17]). Correlations are shown for the Patch-seq data set. Only statistically significant correlations found in both the Patch-seq and reference FACS data sets are shown. Significance was assessed after multiple comparison correction of p-values. PCs and genes with no significant correlations are omitted.

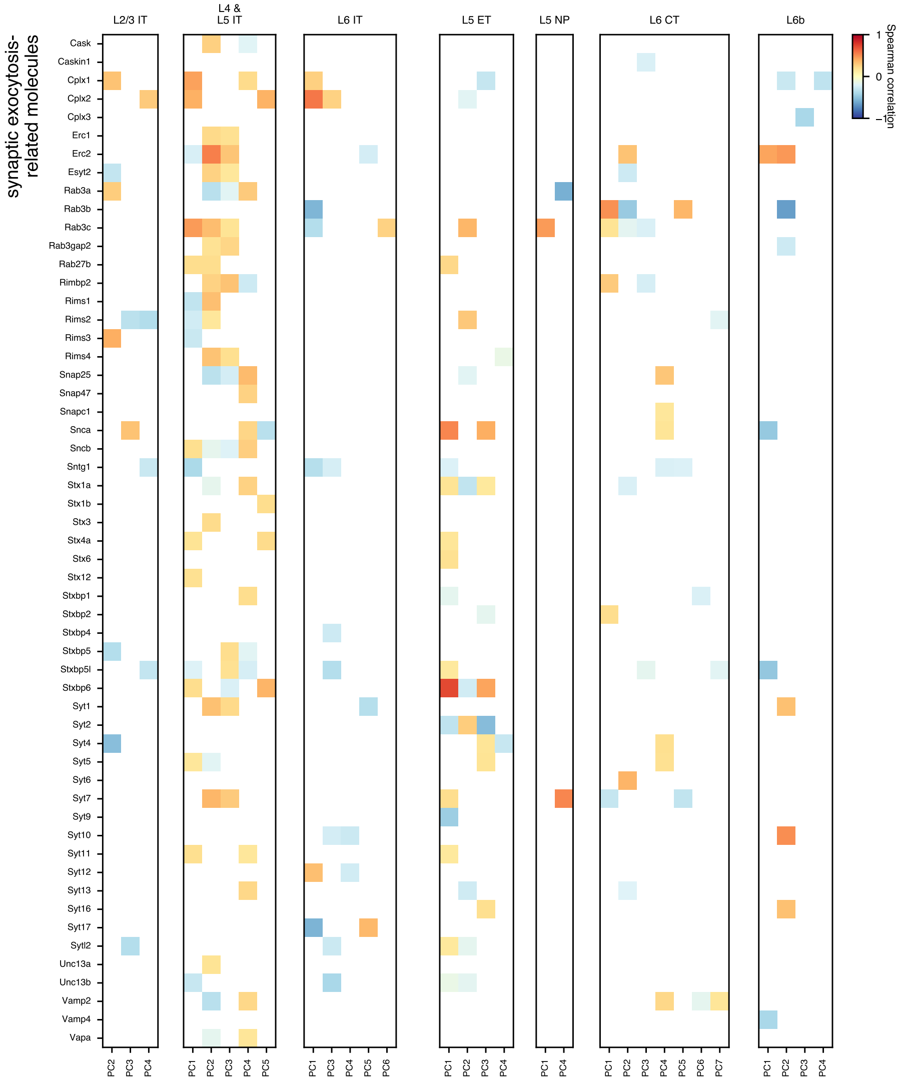


**Extended Data Figure 23: Correlations between synaptic exocytosis-related gene expression and transcriptomic PCs.** Spearman correlations were measured between the transcriptomic PCs (Tx PCs) and synaptic exocytosis-related gene expression (list from Földy *et al.* [17]). Correlations are shown for the Patch-seq data set. Only statistically significant correlations found in both the Patch-seq and reference FACS data sets are shown. Significance was assessed after multiple comparison correction of p-values. PCs and genes with no significant correlations are omitted.

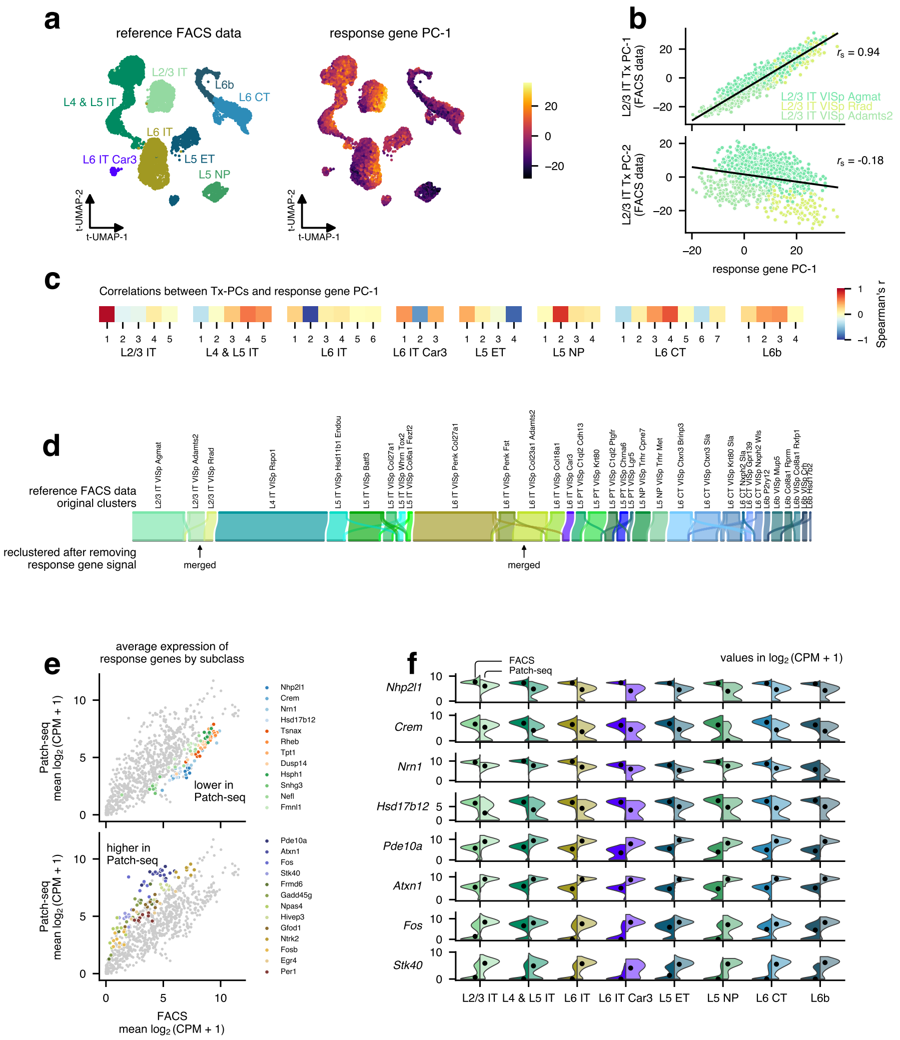
a
reference FACS data
response gene PC-1
L2/3 IT
L4 & L5 IT
L6b
L6 CT
L6 IT
L6 IT Car3
L5 ET
L5 NP
t-UMAP-2
t-UMAP-1
b
L2/3 IT Tx PC-1 (FACS data)
L2/3 IT Tx PC-2 (FACS data)
$r_s$ = 0.94
$r_s$ = -0.18
L2/3 IT VISp Agmat
L2/3 IT VISp Rrad
L2/3 IT VISp Adamts2
response gene PC-1
c
Correlations between Tx-PCs and response gene PC-1
L2/3 IT
L4 & L5 IT
L6 IT
L6 IT Car3
L5 ET
L5 NP
L6 CT
L6b
Spearman's r
d
reference FACS data original clusters
reclustered after removing response gene signal
merged
merged
e
average expression of response genes by subclass
Patch-seq mean $\log_2$(CPM + 1)
lower in Patch-seq
higher in Patch-seq
Nhp2l1
Crem
Nrn1
Hsd17b12
Tsnax
Rheb
Tpt1
Dusp14
Hsph1
Snhg3
Nefl
Fmnl1
Pde10a
Atxn1
Fos
Stk40
Frmd6
Gadd45g
Npas4
Hivep3
Gfod1
Ntrk2
Fosb
Egr4
Per1
FACS mean $\log_2$(CPM + 1)
f
FACS
Patch-seq
values in $\log_2$(CPM + 1)
Nhp2l1
Crem
Nrn1
Hsd17b12
Pde10a
Atxn1
Fos
Stk40
L2/3 IT
L4 & L5 IT
L6 IT
L6 IT Car3
L5 ET
L5 NP
L6 CT
L6b

**Extended Data Figure 24** ***(preceding page)*****: Activity-dependent genes contribute to transcriptomic variation within subclasses.** **a**, UMAP plots of the reference FACS data showing transcriptomic subclasses (left) and variation in activity-dependent genes summarized by a response gene principal component. The response gene first principal component (PC-1) was calculated from the expression across all visual cortex excitatory cells of 108 early- and late-response genes identified as being increased after visual stimulus by Hrvatin *et al.* [75]. Variation in this activity-related signal is evident in several subclasses, such as L2/3 IT and L6 IT. **b**, Correlations between the first (top) and second (bottom) transcriptomic principal components in the L2/3 IT subclass and the response gene PC-1. The first component is highly correlated, with the T-type L2/3 IT VISp Rrad exhibiting higher values and L2/3 IT VISp Adamts2 exhibiting lower values. **c**, Correlations between transcriptomic principal components across all subclasses. **d**, Results of reclustering the reference FACS data after removing transcriptomic dimensions that exhibit high correlation (>0.7) with the response gene PC-1. Note that two L2/3 IT T-types and two L6 IT T-types are merged by this reclustering. **e**, Comparison of early- and late-response gene expression between the FACS and Patch-seq data sets. Each point represents the mean gene expression of one of the 108 genes in one of the eight transcriptomic subclasses. Several genes exhibited lower expression consistently across subclasses in the Patch-seq cells (top) while others exhibited higher expression (bottom) in the Patch-seq data set (see Methods). **f**, Example response gene expression in FACS and Patch-seq data sets by transcriptomic subclass.

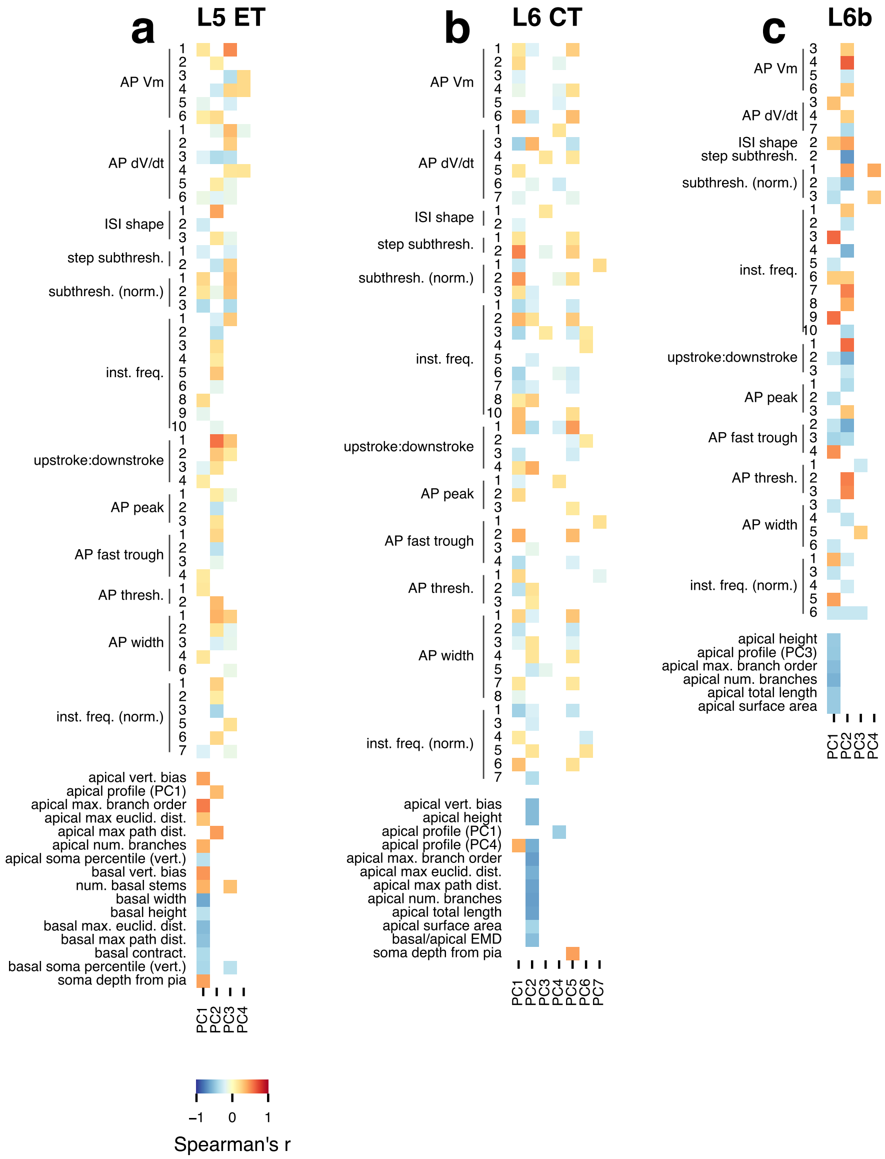


**Extended Data Figure 25: Cross-modal correlations within ET, CT, and L6b subclasses.** Statistically significant Spearman correlations between transcriptomic principal components (PCs) and electrophysiological (upper) and morphological (lower) features are shown in the heat maps for the L5 ET (**a**), L6 CT (**b**), and L6b (**c**) subclasses. Only PCs and features with at least one significant correlation are shown.

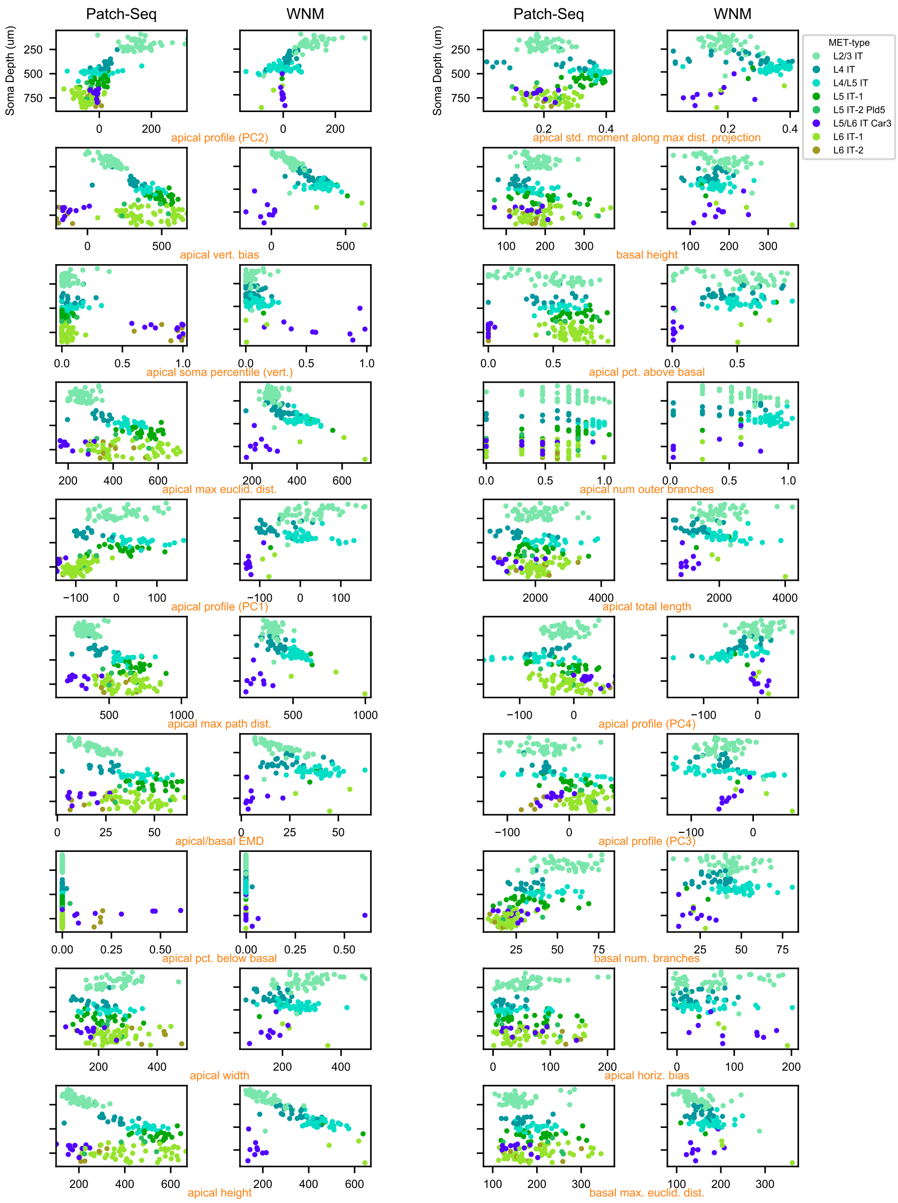


**Extended Data Figure 26: Comparison of dendritic morphology features between Patch-seq and WNM IT MET-types**. The graphs depict the relationship between soma depth from pia and the twenty most informative features for distinguishing IT MET-types, as determined by the Gini index.

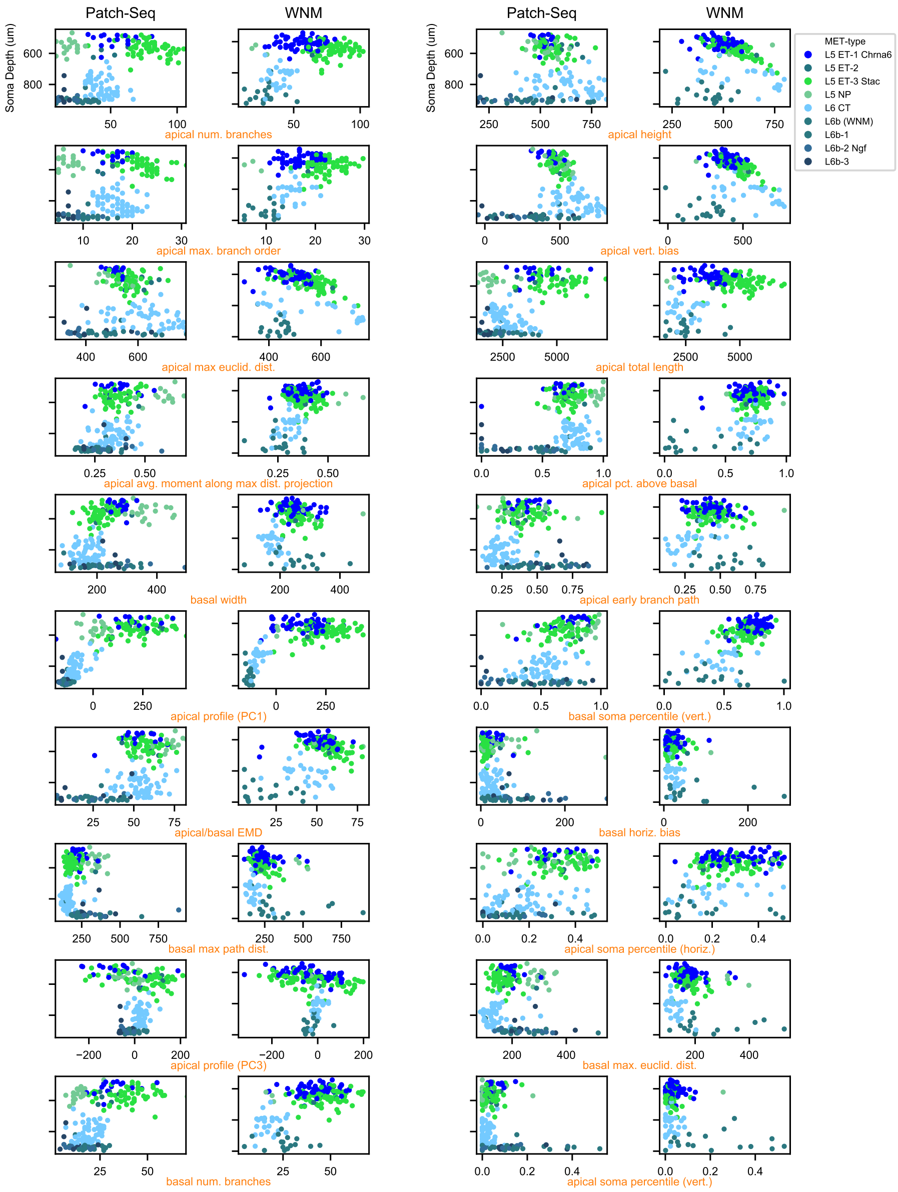


**Extended Data Figure 27: Comparison of dendritic morphology features between Patch-seq and WNM non-IT MET-types**. The graphs depict the relationship between soma depth from pia and the twenty most informative features for distinguishing non-IT MET-types, as determined by the Gini index.

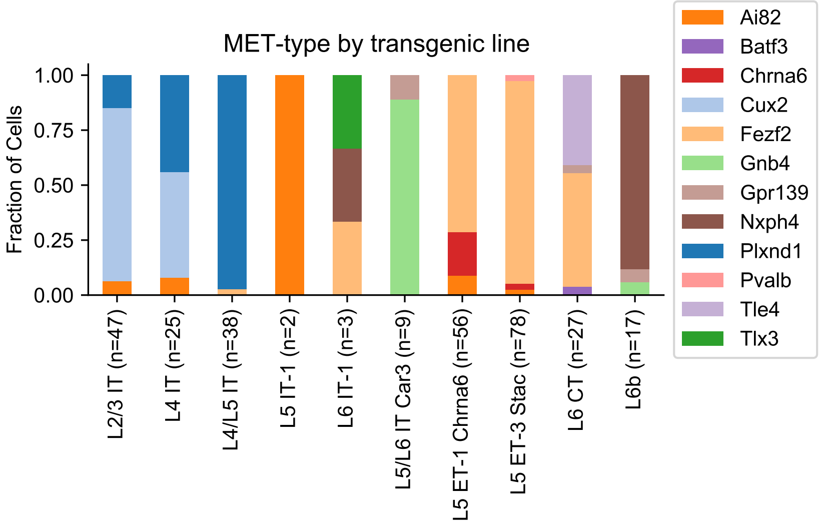


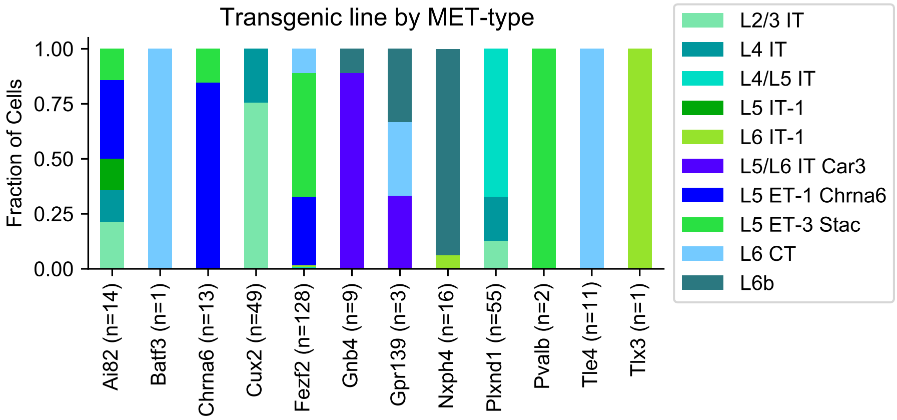


**Extended Data Figure 28: Predicted MET-type by transgenic line and transgenic line by predicted MET-type**

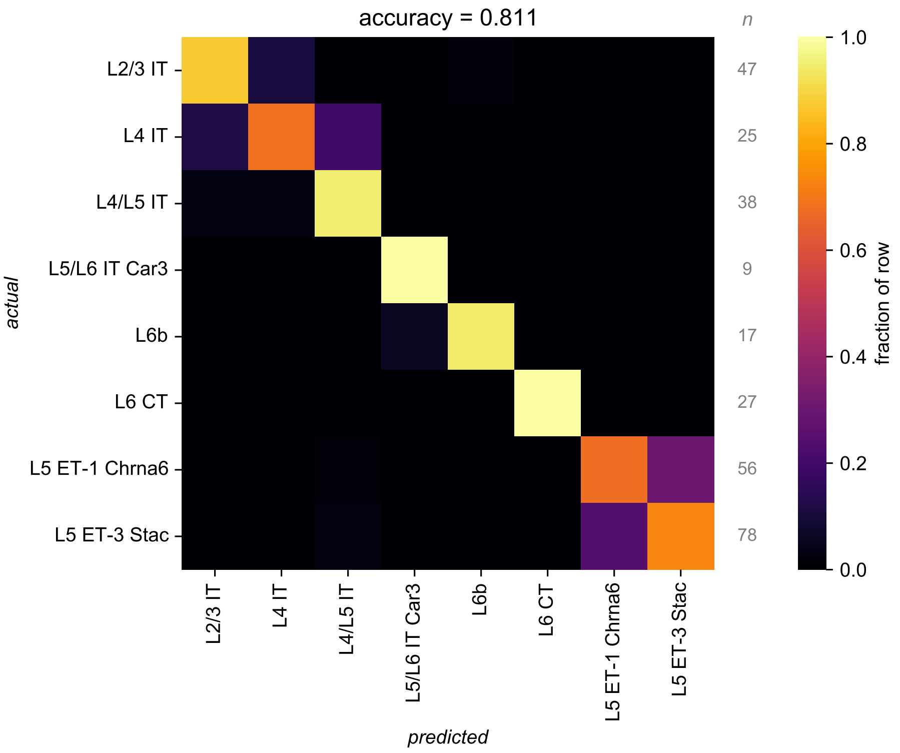


**Extended Data Figure 29: Local axon predicts predicted MET-type.** Local axon features were used to train a random forest classifier to classify the predicted MET-type labels.

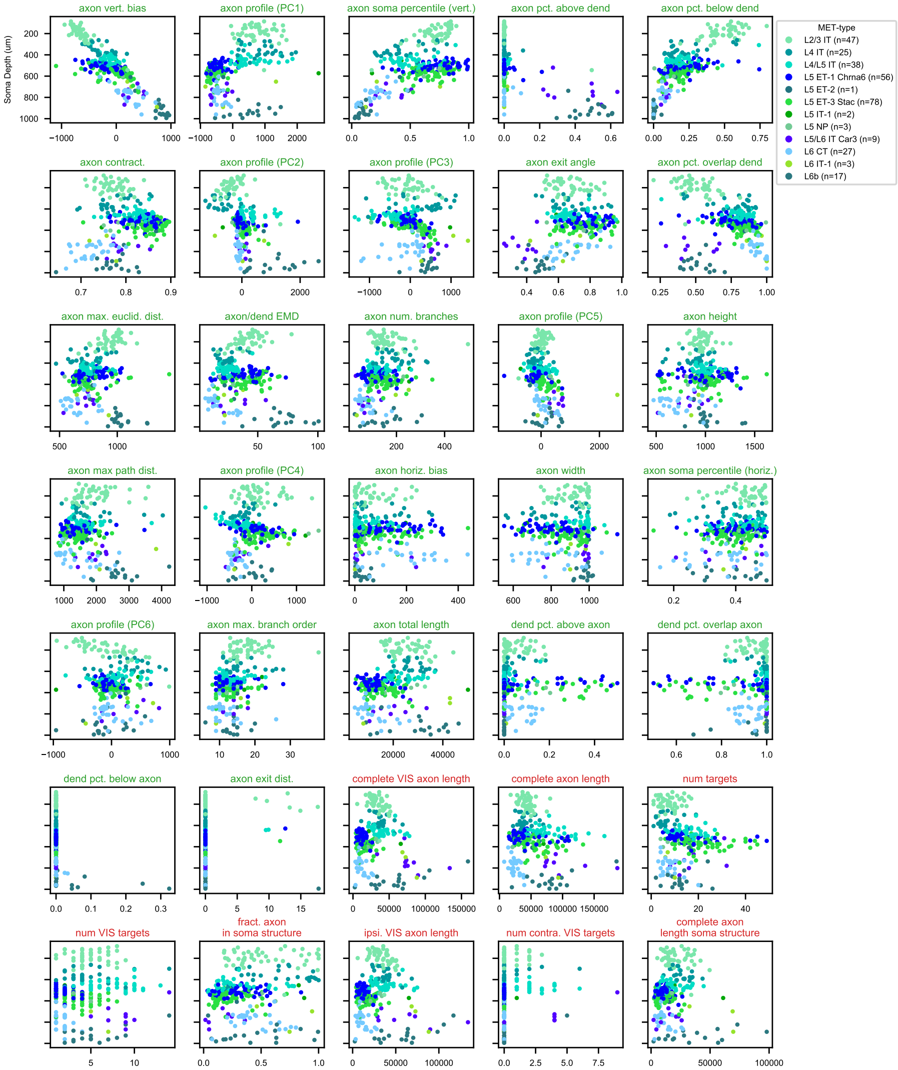


**Extended Data Figure 30: Comparison of local (green titles) and whole (red titles) axon morphology features across predicted MET-types.** The graphs depict the relationship between soma depth from pia and each feature.

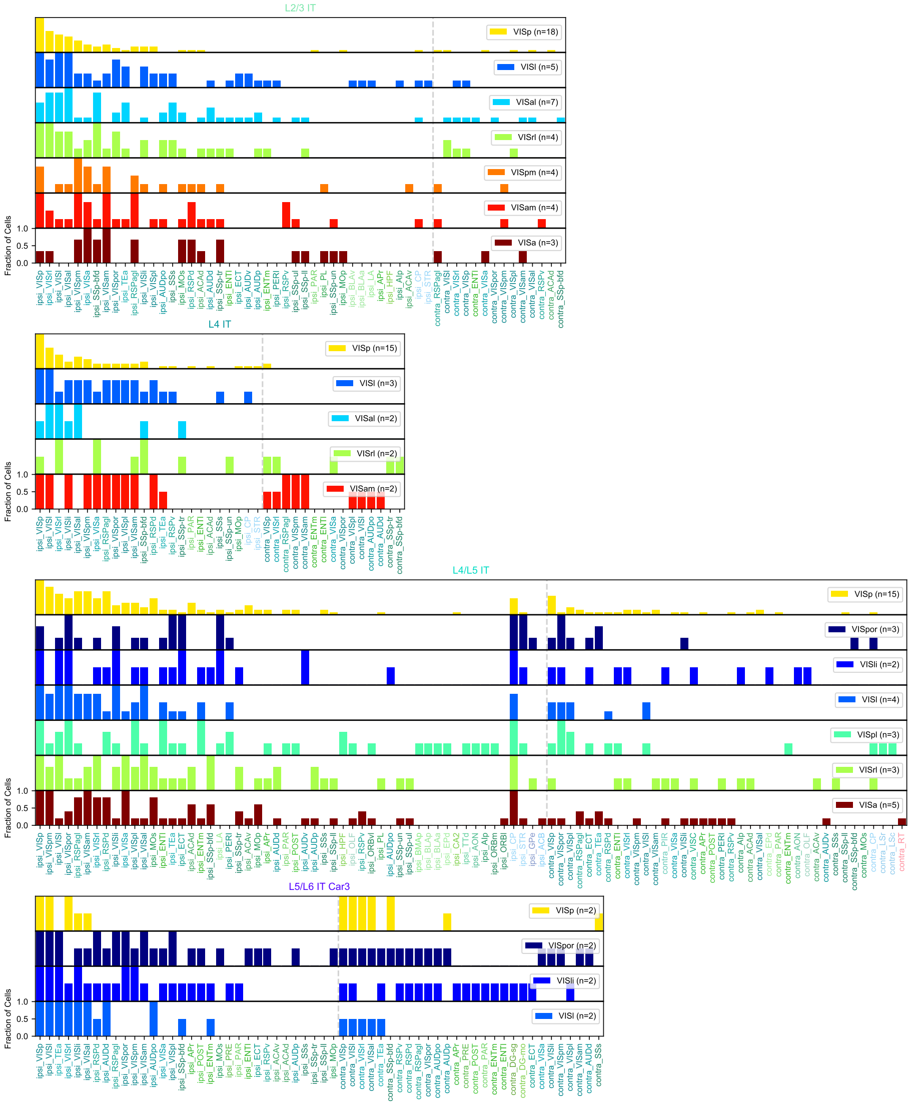

**Extended Data Figure 31: IT predicted MET-type projection summaries**. Fraction of cells that project to target regions separated by MET-type and soma structure.

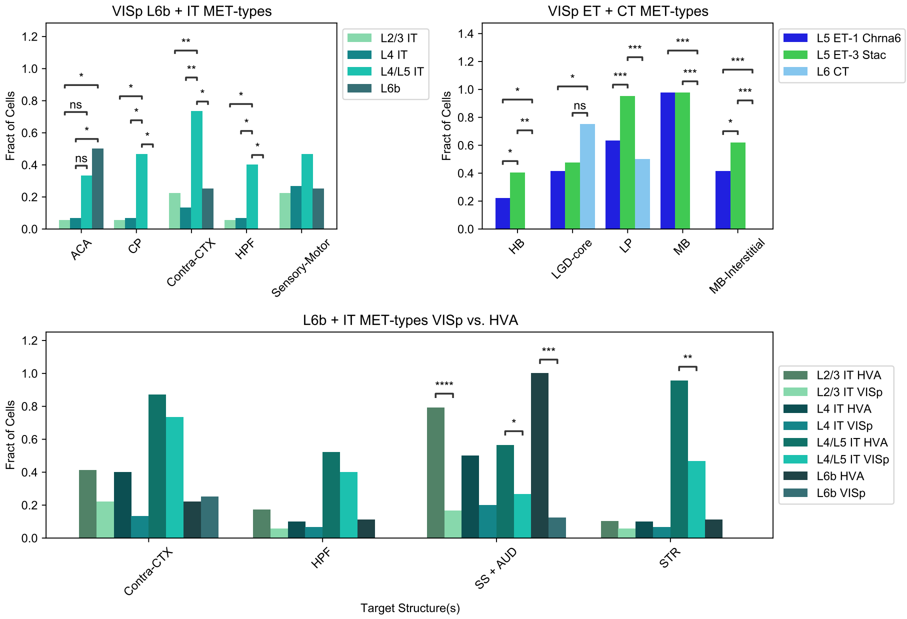


**Extended Data Figure 32: Differences in projections at the population level**. Monte Carlo permutation tests were run over 10,000 iterations to determine significant differences in the fraction of cells that project to specified target regions from VISp (top). Within MET-types, differences across VISp and HVA populations were observed (bottom).

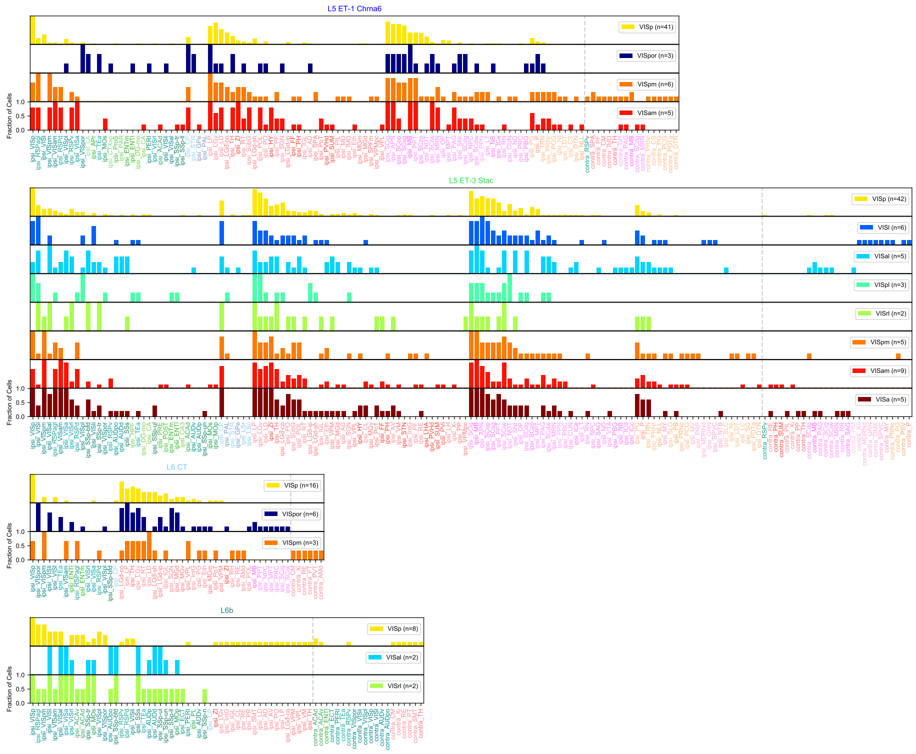


**Extended Data Figure 33: Non-IT predicted MET-type projection summaries**. Fraction of cells that project to target regions separated by MET-type and soma structure.

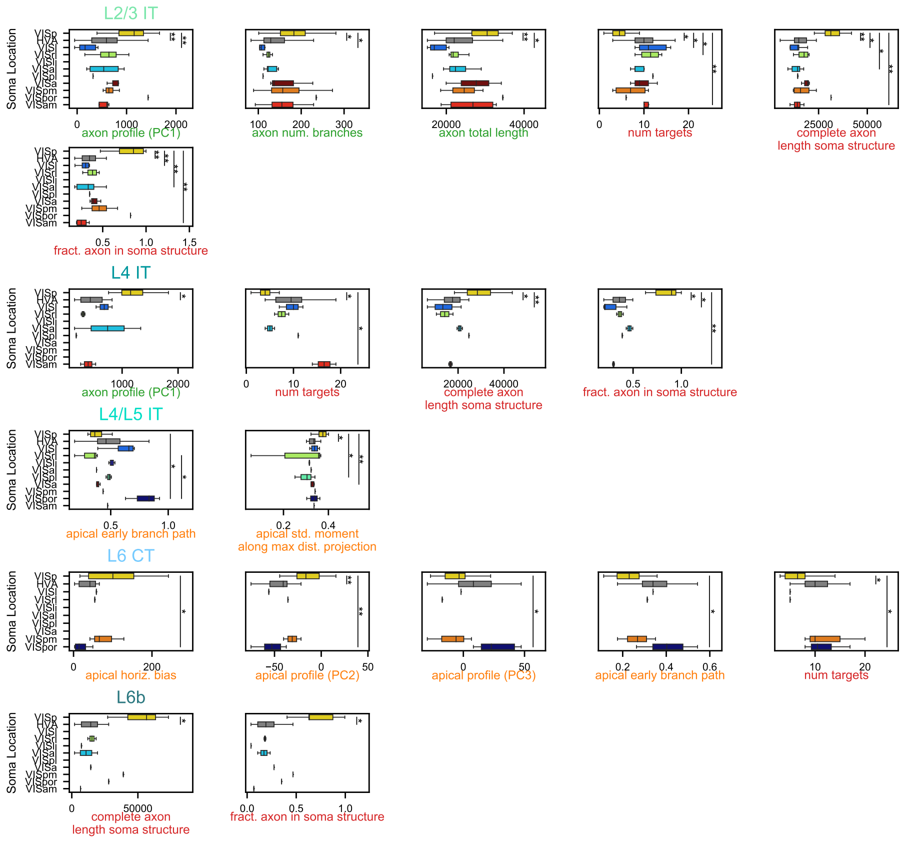

**Extended Data Figure 34: Local axon (green), complete/whole neuron axon (red) and dendrite (orange) features with observed differences across VIS regions in non-ET MET-types.** Significant differences were found using the Kruskal-Wallis test followed by Benjamini-Hochberg corrected post-hoc Dunn tests.

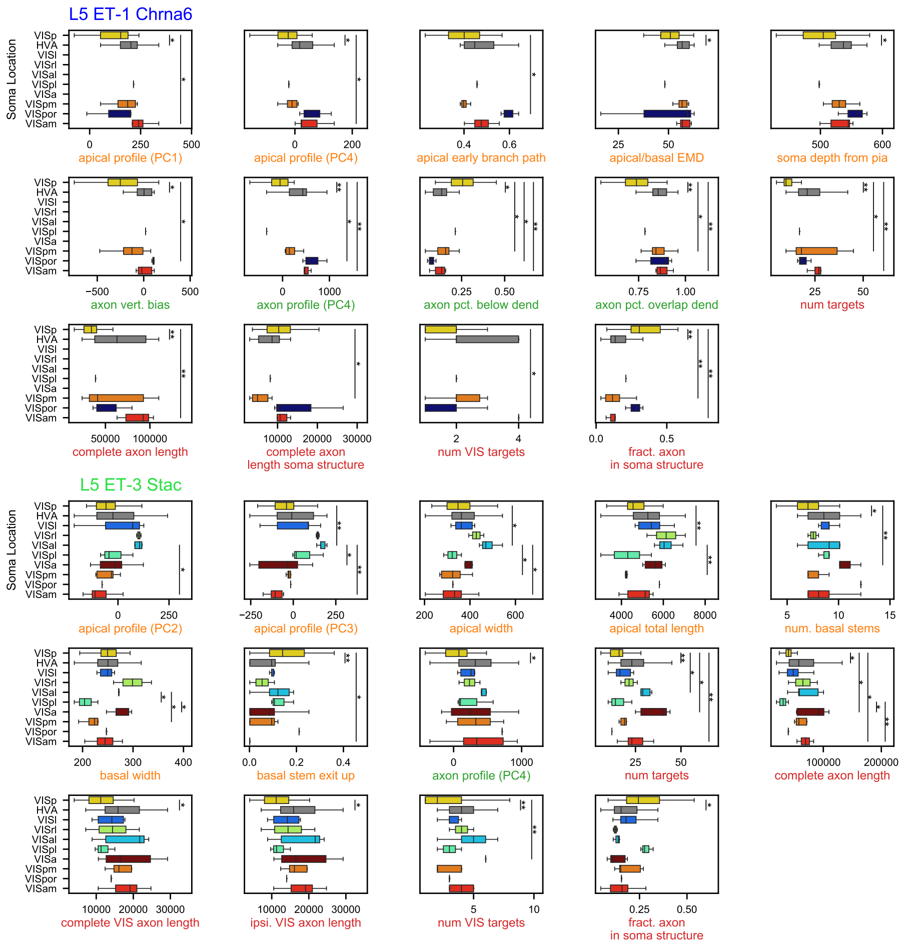


**Extended Data Figure 35: Local axon (green), complete/whole neuron axon (red) and dendrite (orange) features with observed differences across VIS regions in ET MET-types.** Significant differences were found using the Kruskal-Wallis test followed by Benjamini-Hochberg corrected post-hoc Dunn tests.

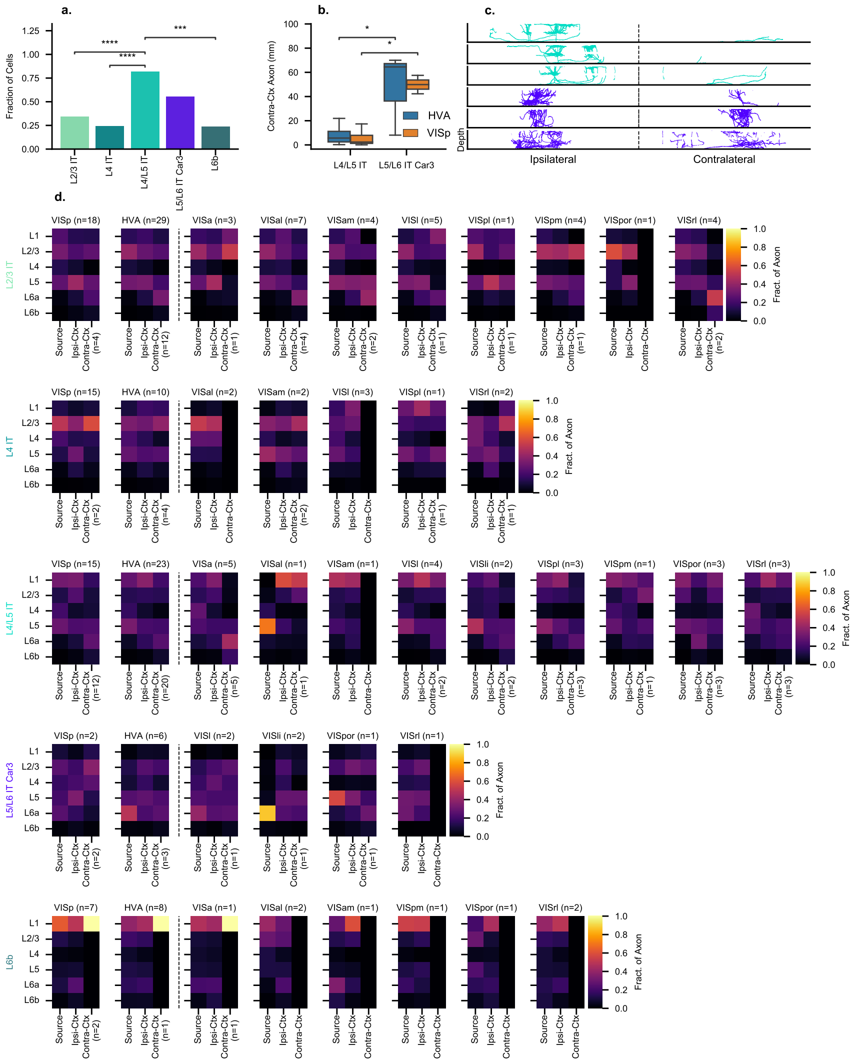

**Extended Data Figure 36: Summary of cortical projections across IT and L6b MET-types**. **a**, Monte Carlo permutation tests were run over 10,000 iterations to determine significant differences in the fraction of cells that target contralateral cortex. **b**, Kruskal-Wallis and post hoc Dunn tests were conducted to show significant differences in total contralateral axon length. **c**, Views of contralaterally projecting morphologies from L4/L5 IT and L5/L6 IT Car3 through the cortical depth. **d**, Laminar projection summaries are displayed for cells from each MET-type and soma structure. The Source label represents the proportion of axon within the structure containing a cell's soma. Ipsi-Ctx represents the proportion of axon in the ipsilateral cortex that lies outside of the source structure. Contra-Ctx represents the fraction of axon in the contralateral cortex when present.

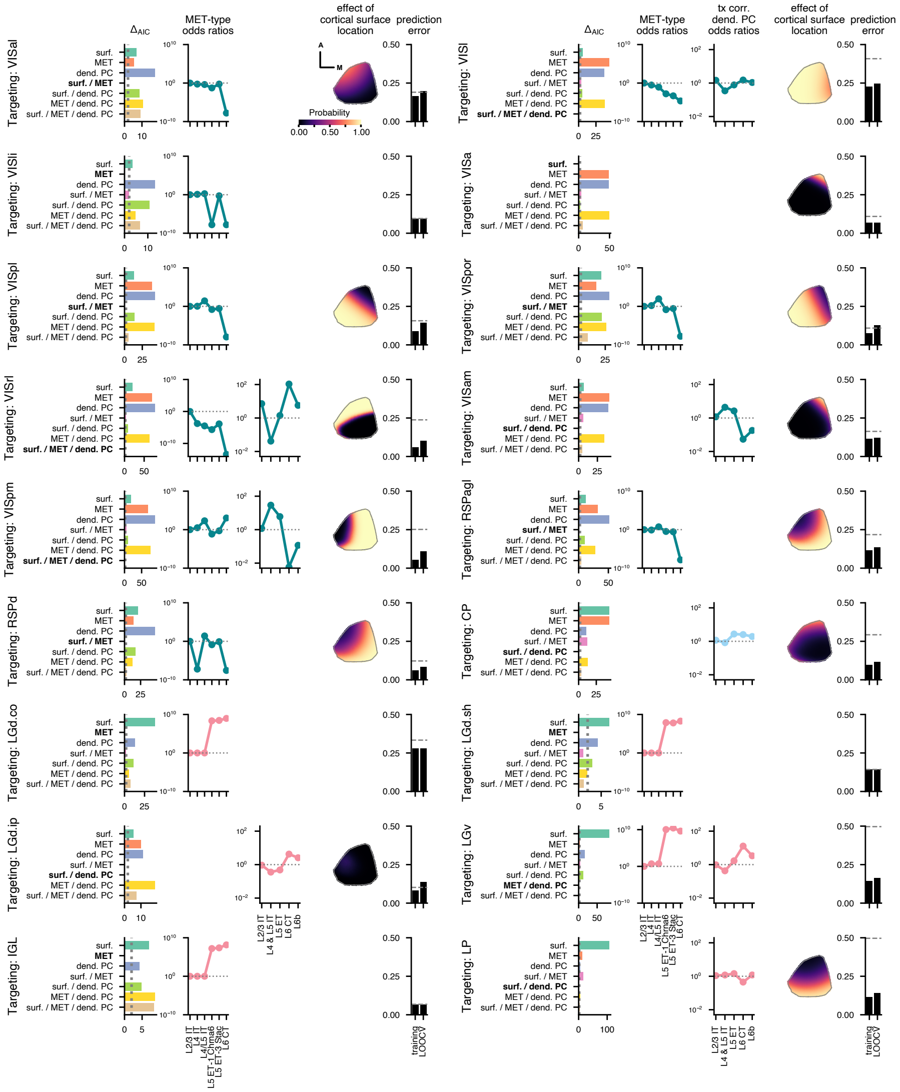
Targeting: VISal
Targeting: VISli
Targeting: VISpl
Targeting: VISrl
Targeting: VISpm
Targeting: RSPd
Targeting: LGd.co
Targeting: LGd.ip
Targeting: IGL
Targeting: VISl
Targeting: VISa
Targeting: VISpor
Targeting: VISam
Targeting: RSPagl
Targeting: CP
Targeting: LGd.sh
Targeting: LGv
Targeting: LP
ΔAIC
MET-type odds ratios
tx corr. dend. PC odds ratios
effect of cortical surface location
prediction error
surf.
MET
dend. PC
surf. / MET
surf. / dend. PC
MET / dend. PC
surf. / MET / dend. PC
Probability
0.00 0.50 1.00
A
M
training
LOOCV
L2/3 IT
L4 IT
L4/L5 IT
L5 ET-1 Chrna6
L5 ET-3 Stac
L6 CT
L4 & L5 IT
L5 ET
L6b

**Extended Data Figure 37** ***(preceding page)*****: Logistic regression models for predicted brain region targets, part 1.** For each target region, the Akaike information criterion (AIC) was used for model selection; the model with the fewest parameters and a $\Delta_{\text{AIC}}$ value below 2 (dotted line) was chosen (bold text label in leftmost plot). When used for a given model, the odds ratios for the predicted MET-type are shown. Also when used, the odds ratios for the transcriptomic-correlated dendritic PCs are shown (odds ratios indicate the change in the odds ratio per unit of the given PC). If surface location is used, the effects of location within VISp on targeting probability is displayed; the probabilities were calculated for the MET-type with the highest odds ratio and the average dendritic PC values across all cells (if part of the model). Prediction errors are calculated for the training data and by leave-one-out cross-validation (LOOCV) with a probability threshold of 0.5. The prediction error for a naive classifier that always chooses the more likely option (targeting or not targeting) is shown by the dashed line.

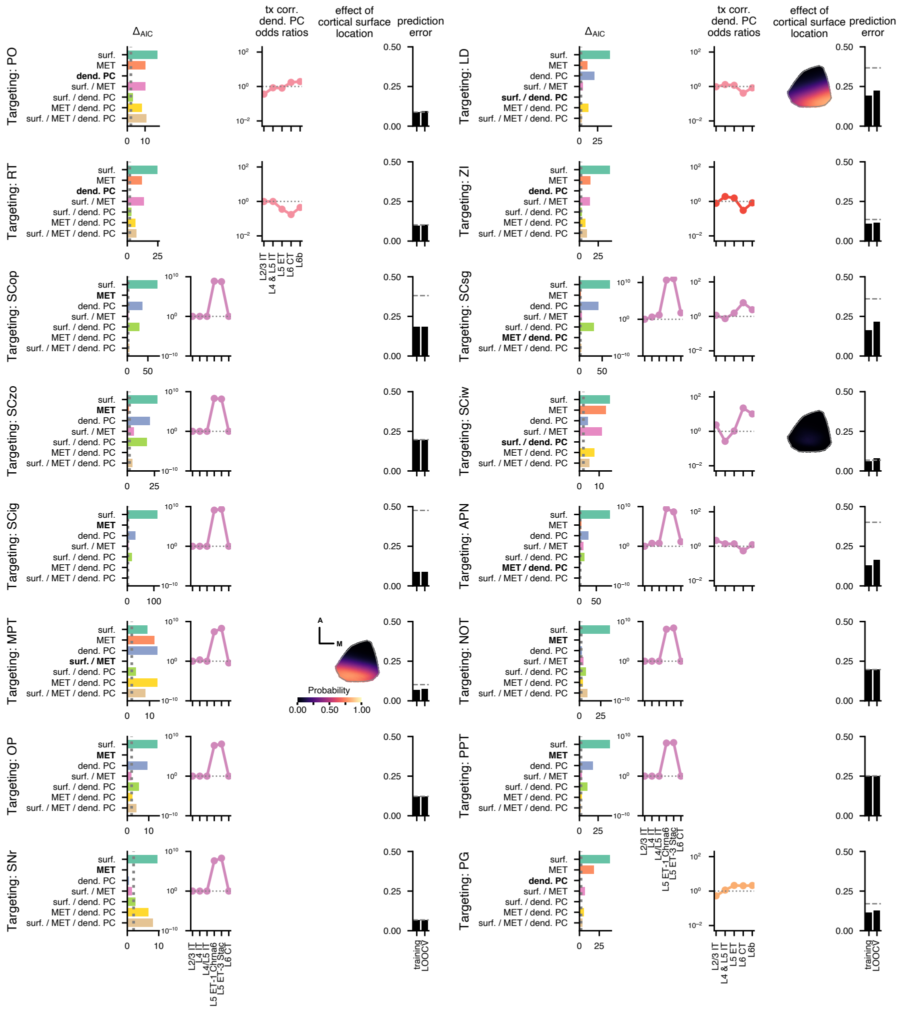

**Extended Data Figure 38 *(preceding page)*: Logistic regression models for predicted brain region targets, part 2.** For each target region, the Akaike information criterion (AIC) was used for model selection; the model with the fewest parameters and a $\Delta_{AIC}$ value below 2 (dotted line) was chosen (bold text label in leftmost plot). When used for a given model, the odds ratios for the predicted MET-type are shown. Also when used, the odds ratios for the transcriptomic-correlated dendritic PCs are shown (odds ratios indicate the change in the odds ratio per unit of the given PC). If surface location is used, the effects of location within VISp on targeting probability is displayed; the probabilities were calculated for the MET-type with highest probability and the average dendritic PC values across all cells (if part of the model). Prediction errors are calculated for the training data and by leave-one-out cross-validation (LOOCV) with a probability threshold of 0.5. The prediction error for a naive classifier that always chooses the more likely option (targeting or not targeting) is shown by the dashed line.

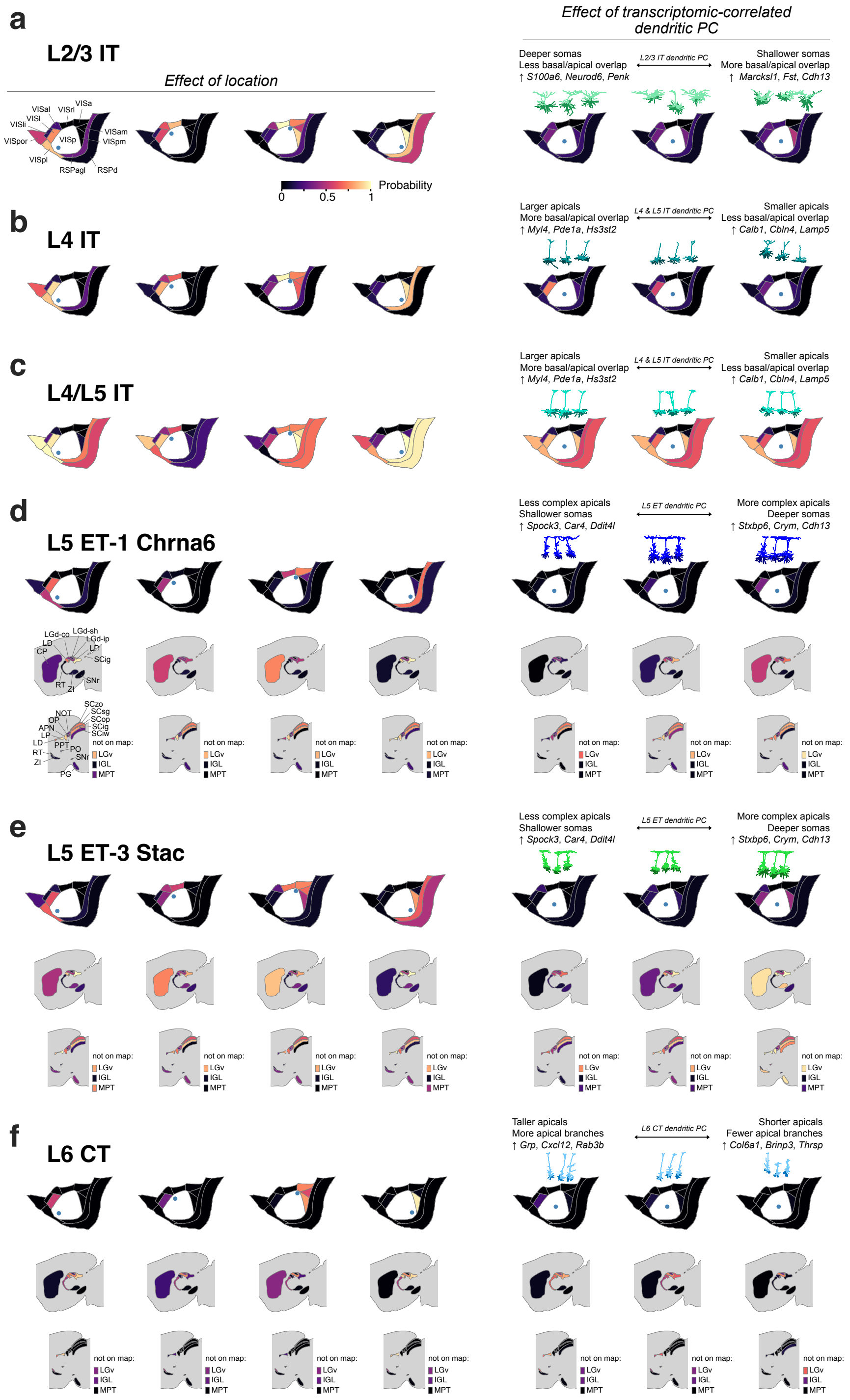
a
L2/3 IT
Effect of location
Effect of transcriptomic-correlated dendritic PC
Probability
b
L4 IT
c
L4/L5 IT
d
L5 ET-1 Chrna6
e
L5 ET-3 Stac
f
L6 CT

**Extended Data Figure 39** ***(preceding page)*****: Predictions of target projections by MET-type. a-c**, Predicted probabilities of projecting to cortical targets from several VISp locations (left) and transcriptomic-correlated dendritic PC values (right) for L2/3 IT (**a**), L4 IT (**b**), and L4/L5 IT (**c**) MET-types. Example neurons (right, above) were chosen at low, medium, and high values of the dendritic PC range for projection probability calculations. Morphological feature variations associated with the dendritic PC and genes inferred to vary based on the corresponding Tx PC are described as well. **d-f**, Predicted probabilities of projecting to cortical and subcortical targets for L5 ET-1 Chrna6 (**d**), L5 ET-3 Stac (**e**), and L6 CT (**f**) MET-types. Effects of location and dendritic PC value are shown as in (**a**)-(**c**).
