## Extended Data Figure 1 all_the_morphs5-5-9 for "Connecting single-cell transcriptomes to projectomes in mouse visual cortex"

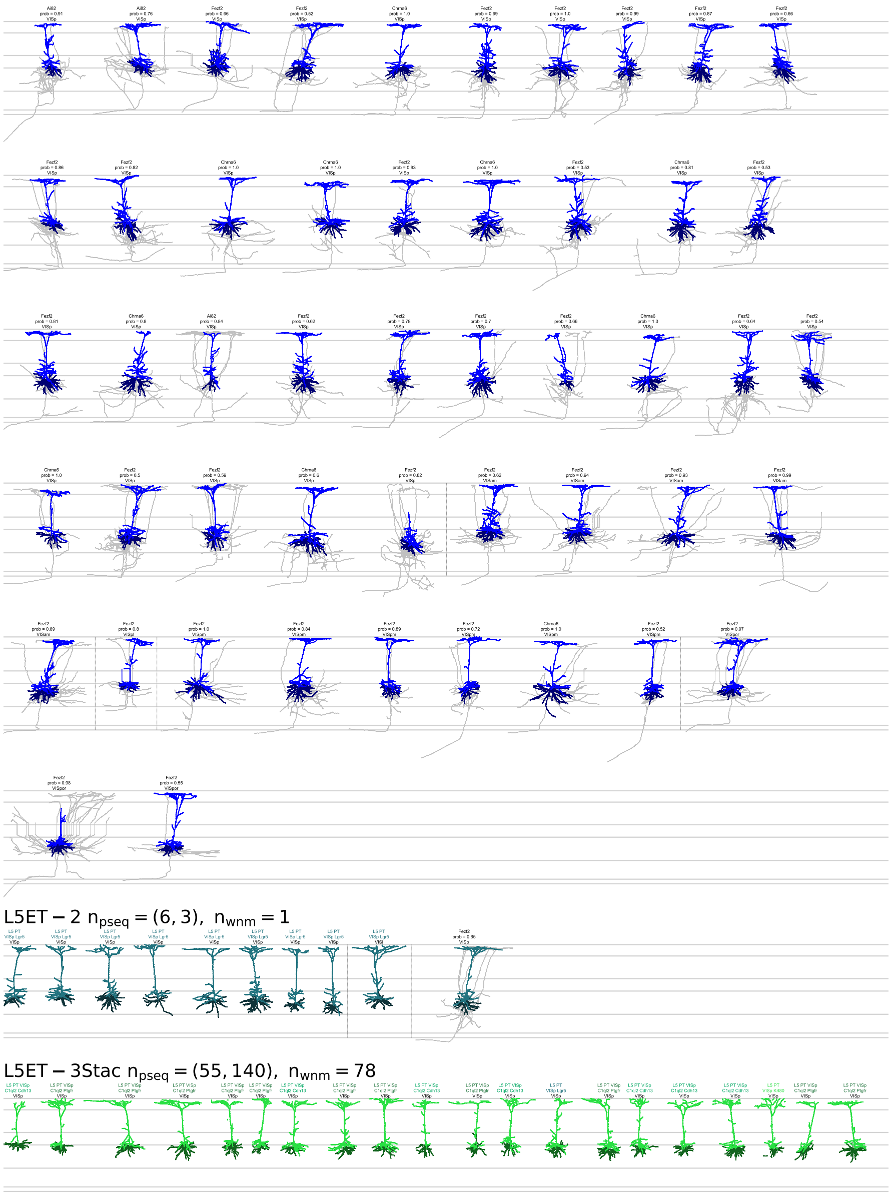


**Extended Data Figure 6: Morphological reconstructions of MET-types (6 of 10)**

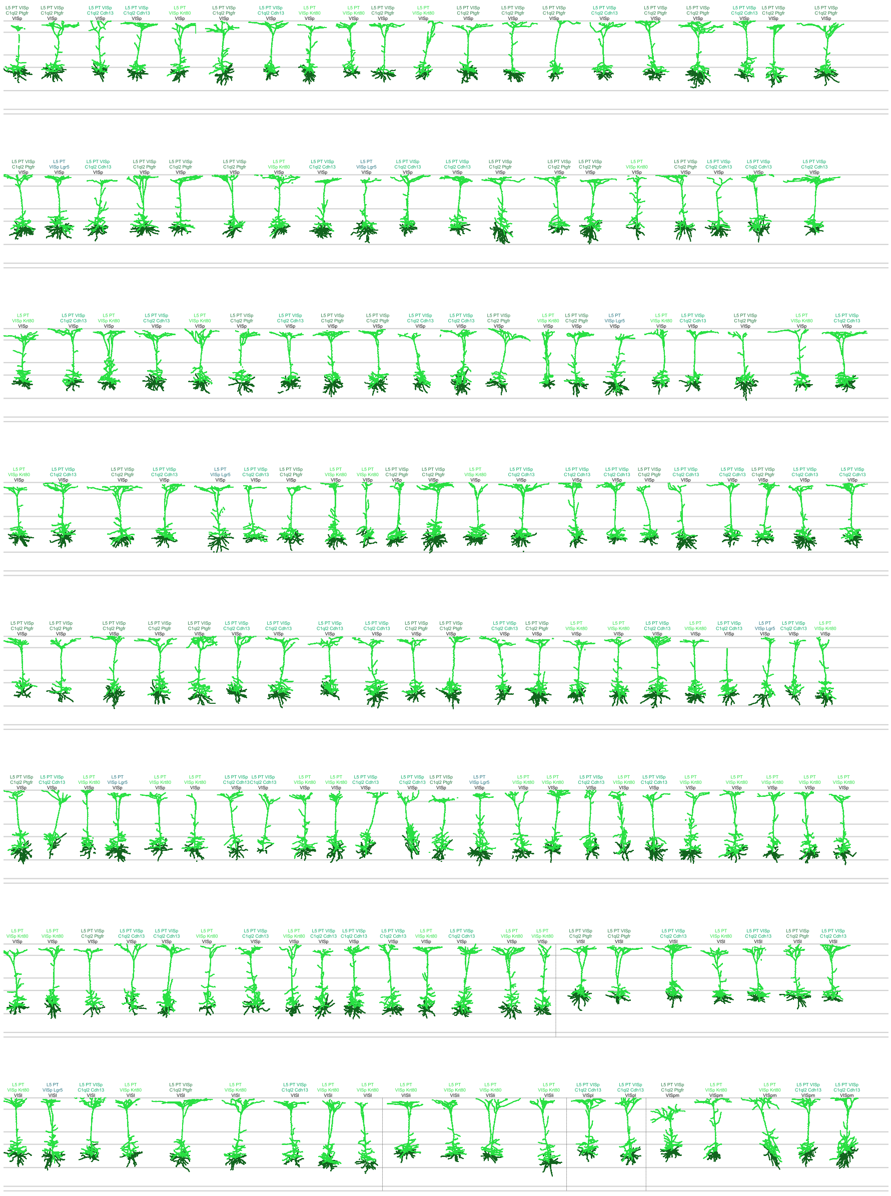

**Extended Data Figure 7: Morphological reconstructions of MET-types (7 of 10)**

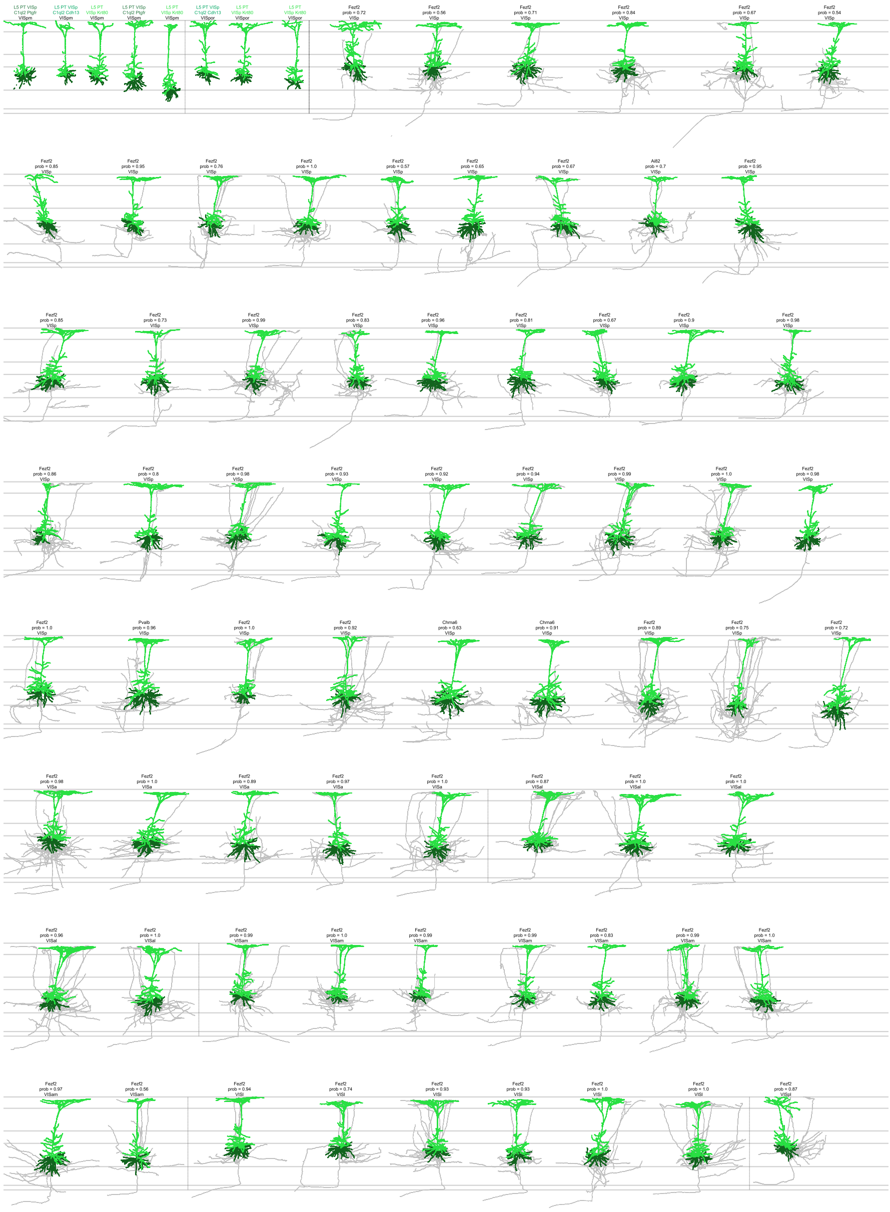

**Extended Data Figure 8: Morphological reconstructions of MET-types (8 of 10)**

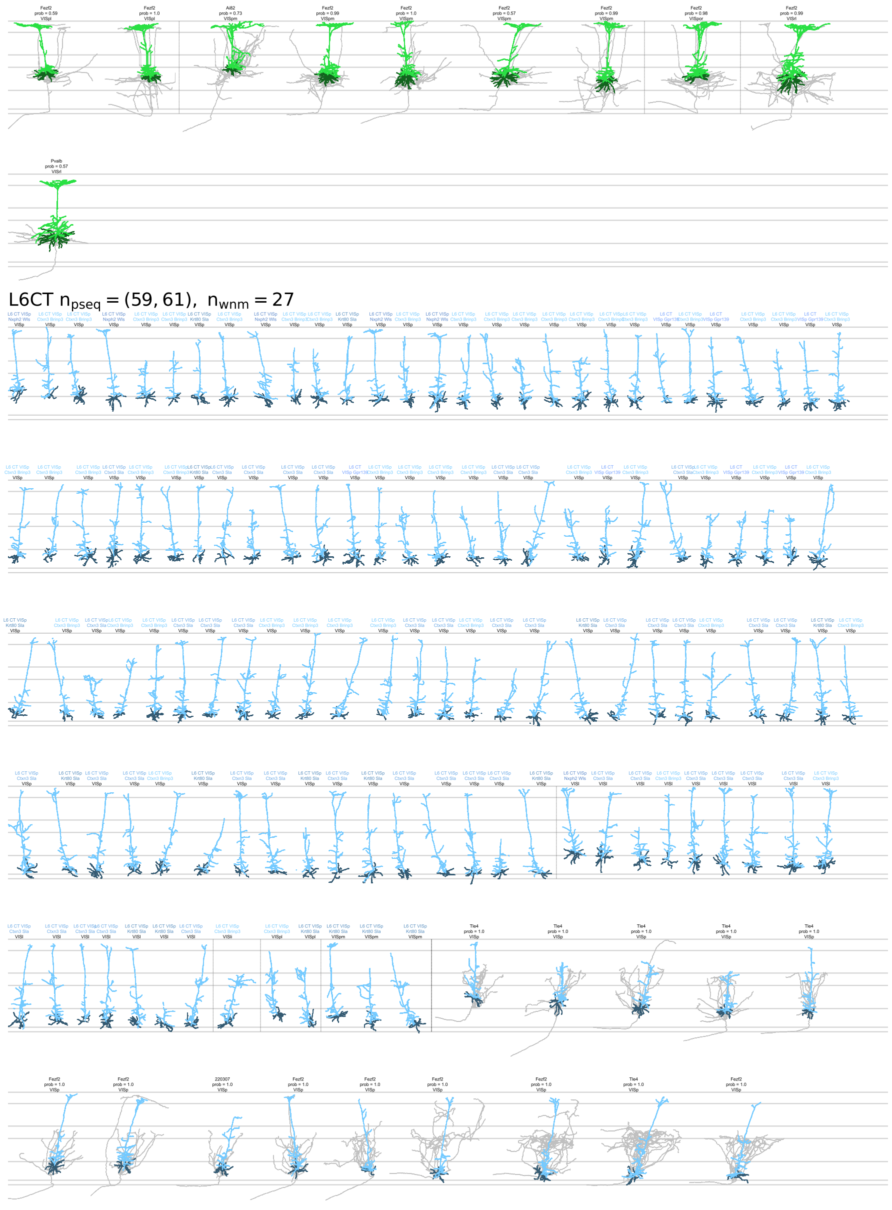


**Extended Data Figure 9: Morphological reconstructions of MET-types (9 of 10)**

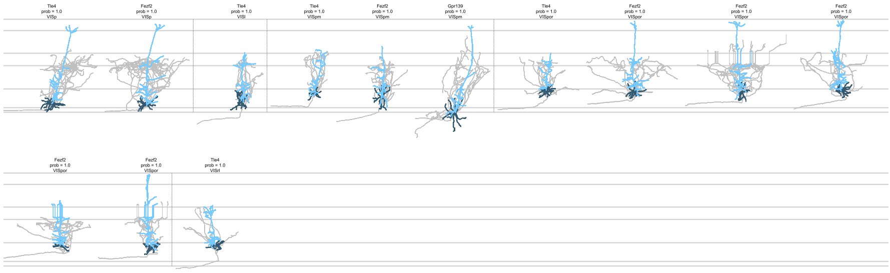


**Extended Data Figure 10: Morphological reconstructions of MET-types (10 of 10)**
