## Extended Data Figure 1 all_the_morphs0-4 for "Connecting single-cell transcriptomes to projectomes in mouse visual cortex"

L2/3IT $n_{pseq} = (50, 2)$, $n_{wnm} = 47$

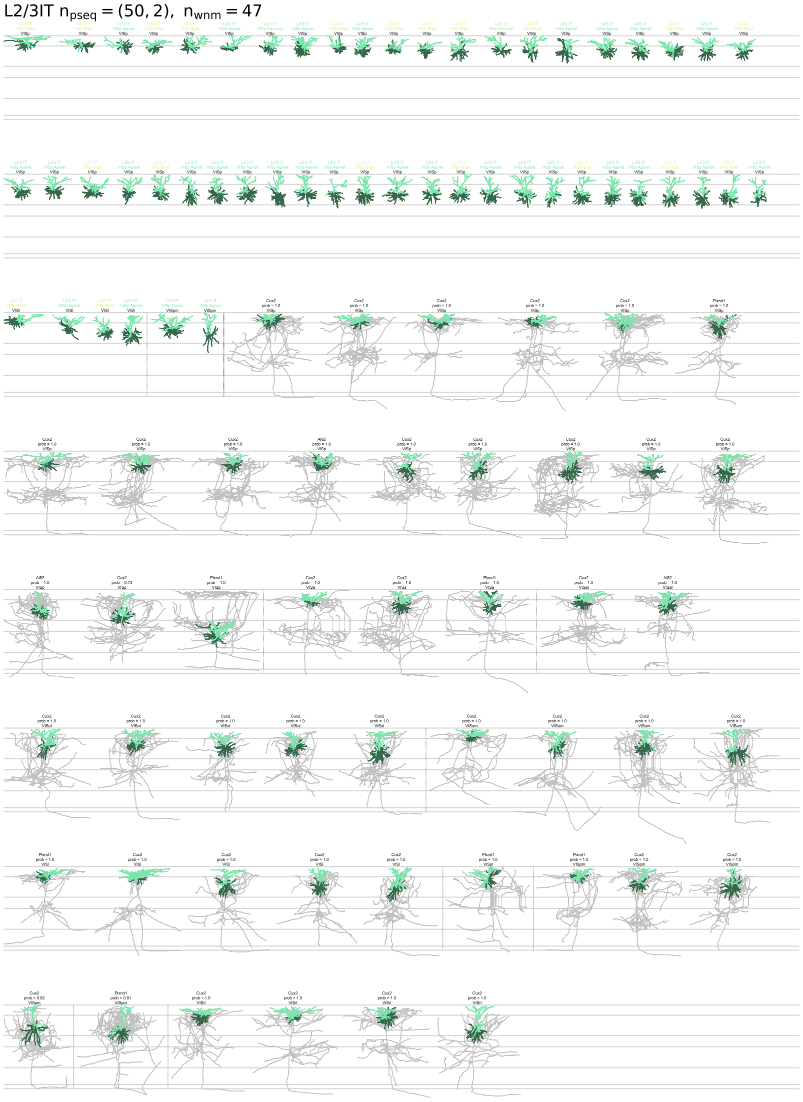

**Extended Data Figure 1: Morphological reconstructions of MET-types (1 of 10)**. Sample sizes indicate the number of Patch-seq neurons (generated manually and automatically respectively) and the number of WNM neurons. Dark dashed lines separate Patch-seq and WNM morphologies, while lighter dashed lines differentiate soma structures. Patch-seq cells include transcriptomic type and soma structure annotations. WNM cells include transgenic line, MET-type prediction probability, and soma structure annotations. Apical dendrites are shaded lighter and local axon is in grey for WNM neurons.

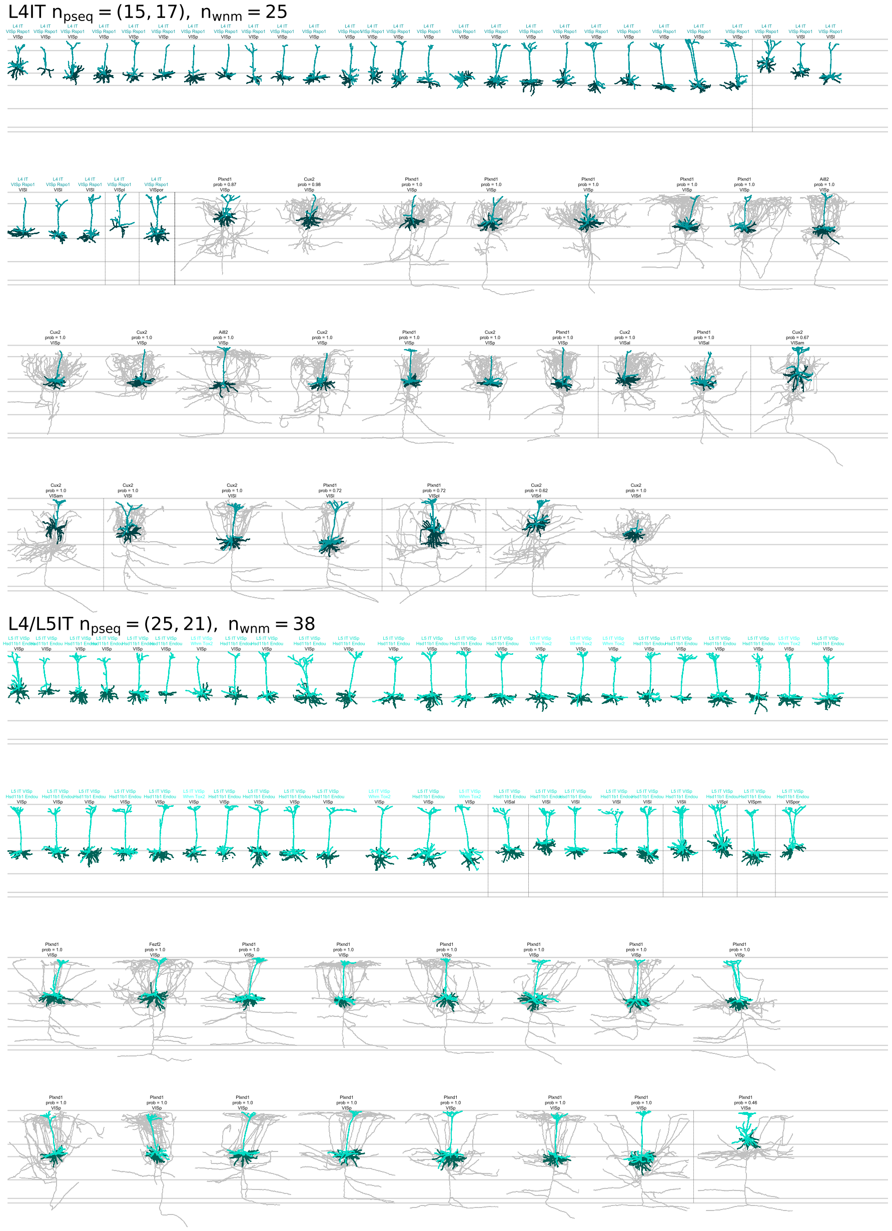


**Extended Data Figure 2: Morphological reconstructions of MET-types (2 of 10)**

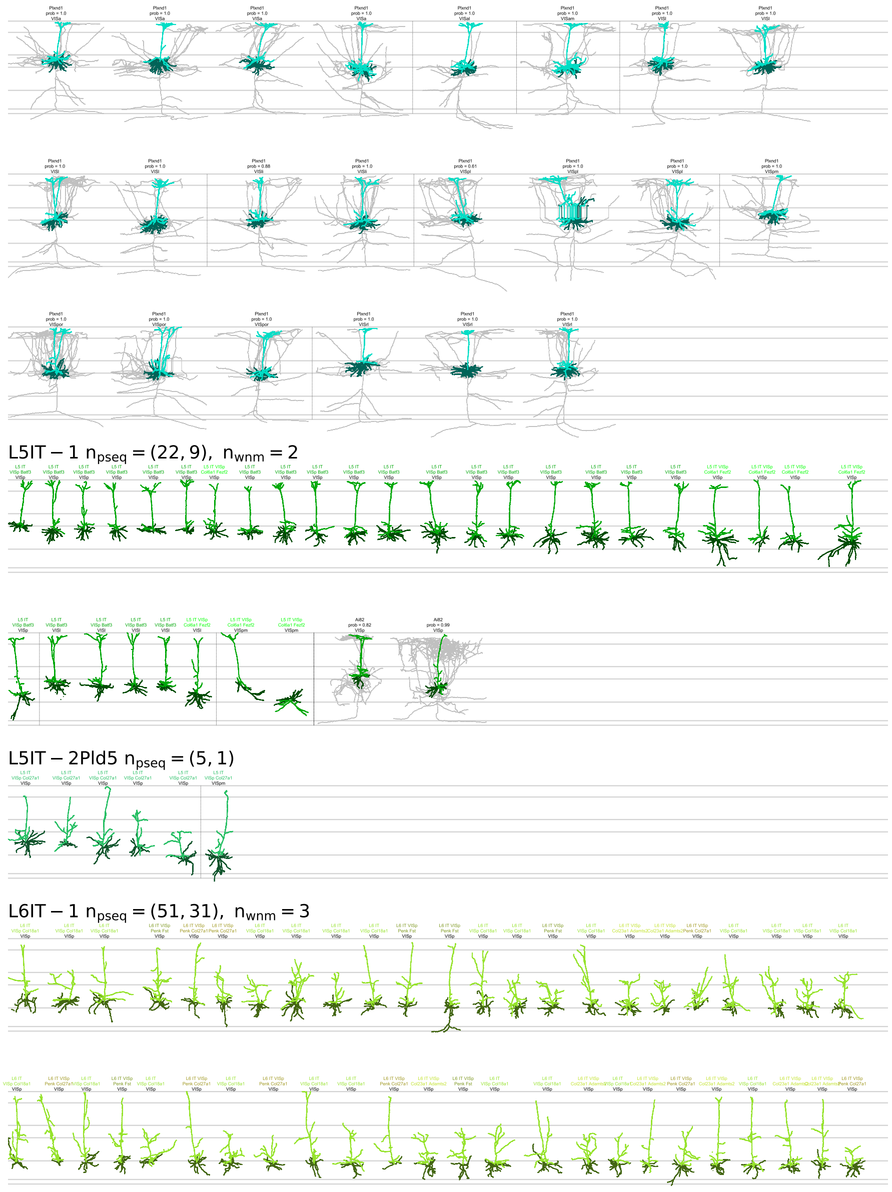


**Extended Data Figure 3: Morphological reconstructions of MET-types (3 of 10)**

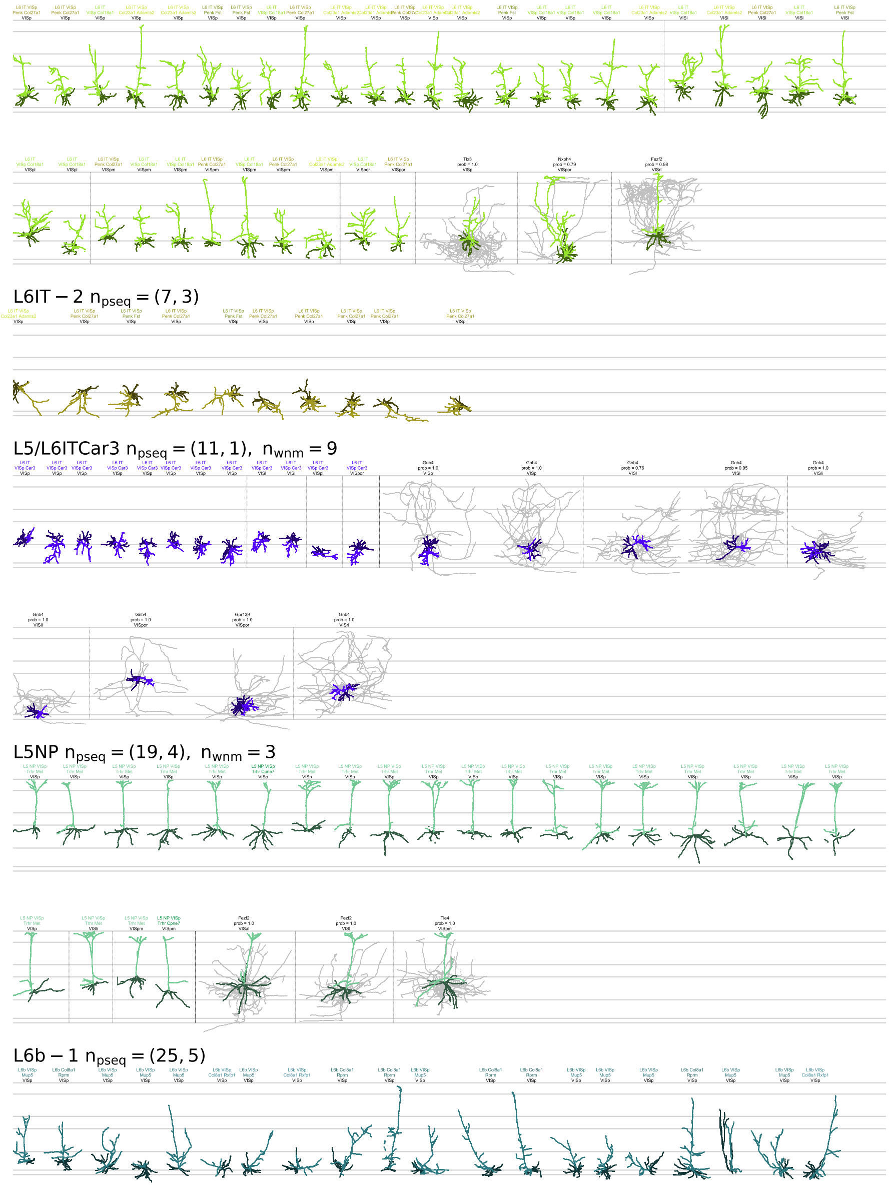


**Extended Data Figure 4: Morphological reconstructions of MET-types (4 of 10)**

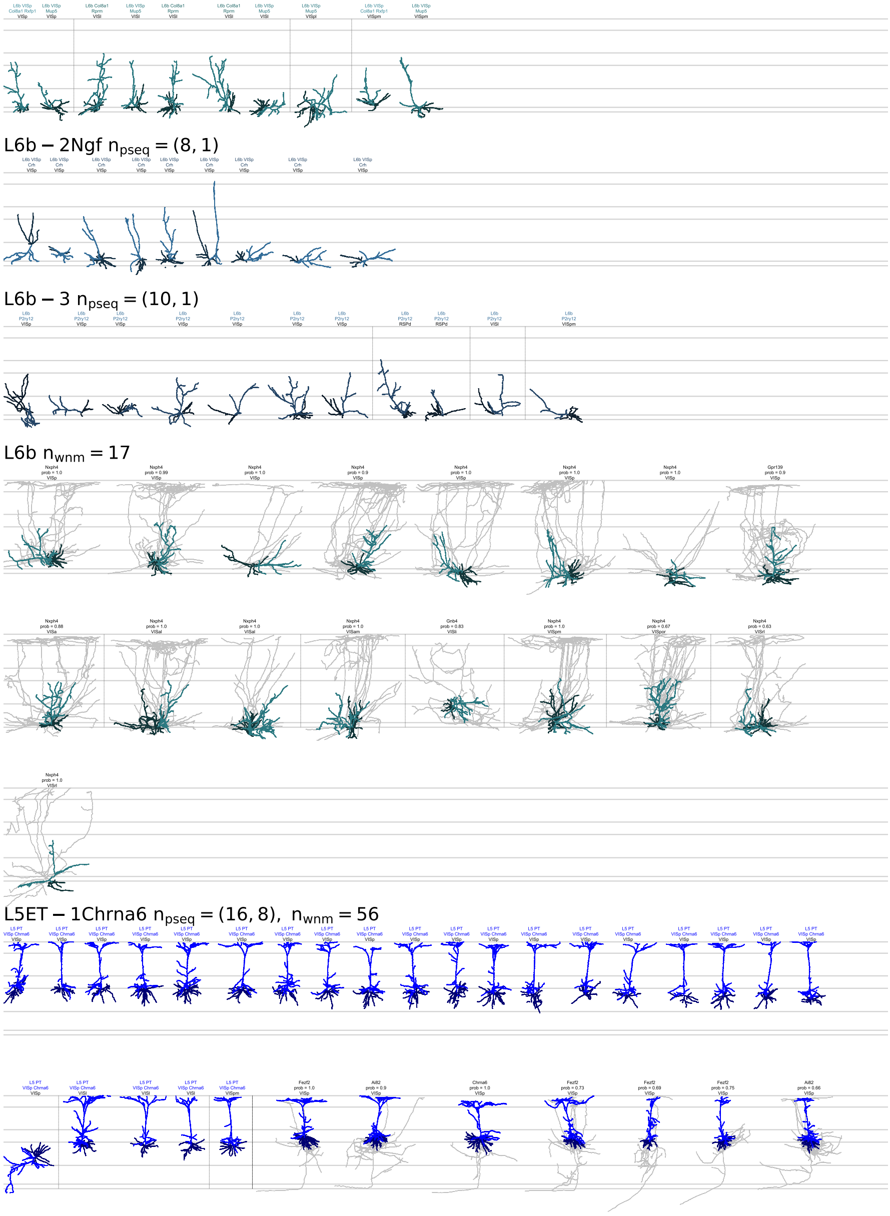


**Extended Data Figure 5: Morphological reconstructions of MET-types (5 of 10)**
